# Benchmarking Uncertainty Quantification for Protein Engineering

**DOI:** 10.1101/2023.04.17.536962

**Authors:** Kevin P. Greenman, Ava P. Amini, Kevin K. Yang

**Affiliations:** Department of Chemical Engineering, Massachusetts Institute of Technology, Cambridge, MA, USA; Microsoft Research New England, Cambridge, MA, USA

**Keywords:** uncertainty quantification, protein engineering, active learning

## Abstract

Machine learning sequence-function models for proteins could enable significant advances in protein engineering, especially when paired with state-of-the-art methods to select new sequences for property optimization and/or model improvement. Such methods (Bayesian optimization and active learning) require calibrated estimations of model uncertainty. While studies have benchmarked a variety of deep learning uncertainty quantification (UQ) methods on standard and molecular machine-learning datasets, it is not clear if these results extend to protein datasets. In this work, we implemented a panel of deep learning UQ methods on regression tasks from the Fitness Landscape Inference for Proteins (FLIP) benchmark. We compared results across different degrees of distributional shift using metrics that assess each UQ method’s accuracy, calibration, coverage, width, and rank correlation. Additionally, we compared these metrics using one-hot encoding and pretrained language model representations, and we tested the UQ methods in a retrospective active learning setting. These benchmarks enable us to provide recommendations for more effective design of biological sequences using machine learning.

## 1 Introduction

Machine learning (ML) has already begun to accelerate the field of protein engineering by providing low-cost predictions of phenomena that require time- and resource-intensive labeling by experiments or physics-based simulations (*1*). It is often necessary to have an estimate of model uncertainty in addition to the property prediction, as the performance of an ML model can be highly dependent on the domain shift between its training and testing data (*2*). Because protein engineering data is often collected in a manner that violates the independent and identically distributed (i.i.d.) assumptions of many ML approaches, (*3*), tailored ML methods are required to guide the selection of new experiments from a protein landscape. Uncertainty quantification (UQ) can inform the selection of experiments in order to improve a ML model or optimize protein function through active learning or Bayesian optimization.

In chemistry and materials science, several studies have benchmarked common UQ methods against one another on standard datasets and have used or developed appropriate metrics to quantify the quality of these uncertainty estimates (*4*–*9*). These works have illustrated that the best choice of UQ method can depend on the dataset and other considerations such as representation and scaling. While some protein engineering work has leveraged uncertainty estimates, these studies have been mostly limited to single UQ methods such as convolutional neural network (CNN) ensembles (*10*) or Gaussian processes (GPs) (*11*, *12*).

Gruver et al. compared CNN ensembles to GPs (using traditional representations and pre-trained BERT (*13*) language model embeddings) in Bayesian optimization (BO) tasks (*14*). They found that CNN ensembles are often more robust to distribution shift than other types of models. Additionally, they report that most model types have more poorly calibrated uncertainties on out-of-domain samples. However, more comprehensive study of CNN UQ methods other than ensembles against GPs using a variety of uncertainty quality metrics has not yet been done. A comparison of uncertainty methods on different protein representations (e.g., one-hot encodings or embeddings from protein language models) in an active learning setting is also lacking.

In this work, we used a set of standardized, public protein datasets to evaluate a panel of UQ methods for protein sequence-function prediction (Figure 1). Our chosen datasets included splits with varied degrees of domain extrapolation, which enabled method evaluation in a setting similar to what might be experienced while collecting new experimental data for protein engineering. We assessed each model using a variety of metrics that captured different aspects of desired performance, including accuracy, calibration, coverage, width, and rank correlation. Additionally, we compared the performance of the UQ methods on one-hot encoded sequence representations and on embeddings computed from the ESM-1b protein masked language model (*15*). We find that the quality of UQ estimates are dependent on the landscape, task, and embedding, and that no single method consistently outperforms all others. Finally, we evaluated the UQ methods in an active learning setting with several acquisition functions, and demonstrated that uncertainty-based sampling often outperforms random sampling (especially in later stages of active learning), although better calibrated uncertainty does not necessarily equate to better active learning. The understanding gained from this work will enable more effective application of UQ techniques to machine learning in protein engineering.

**Figure 1:**
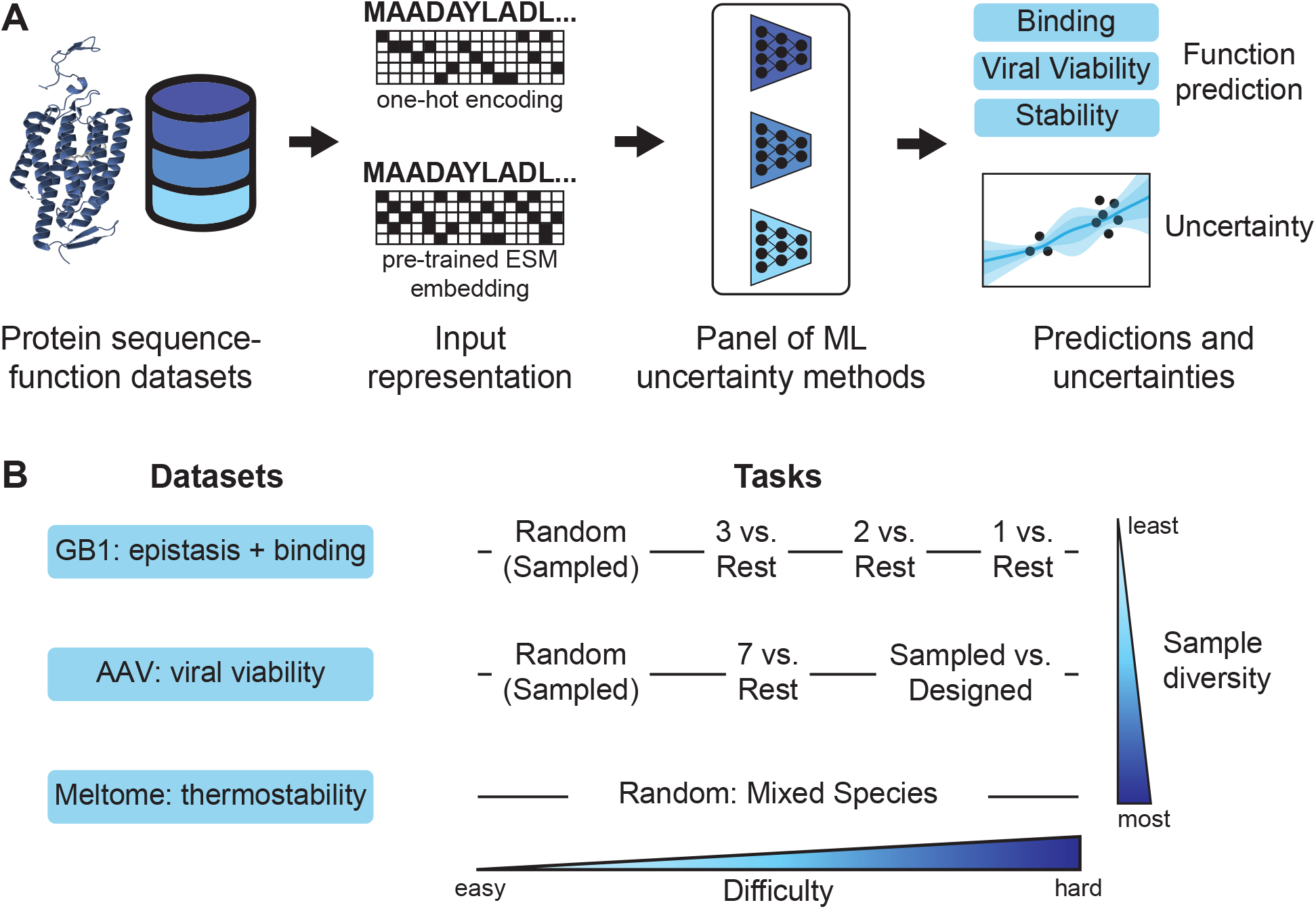
(A) Schematic of the approach for benchmarking uncertainty quantification (UQ) in machine learning for protein engineering. A panel of UQ methods were evaluated on protein fitness datasets to assess the quality of the uncertainty estimates and their utility in active learning. (B) Our study utilized three protein datasets/landscapes and different train-validation-test split tasks within each dataset. These datasets and tasks covered a range of sample diversities and domain shifts (task difficulties).

## 2 Results and Discussion

### 2.1 Uncertainty Quantification

Our first goal was to evaluate the calibration and quality of a variety of UQ methods. We implemented seven uncertainty methods for this benchmark: linear Bayesian ridge regression (BRR) (*16*, *17*), Gaussian processes (GPs) (*18*), and five methods using variations on a convolutional neural network (CNN) architecture. The CNN implementation from FLIP (*3*) provided the core architecture used by our dropout (*19*), ensemble (*20*), evidential (*21*), mean-variance estimation (MVE) (*22*), and last-layer stochastic variational inference (SVI) (*23*) models. Additional model details are provided in Section 4.

The landscapes used in this work were taken from the Fitness Landscape Inference for Proteins (FLIP) benchmark (*3*). These include the binding domain of an immunoglobulin binding protein (GB1), adeno-associated virus stability (AAV), and thermostability (Meltome) data landscapes, which cover a large sequence space and a broad range of protein families. The FLIP benchmark includes several train-test splits, or tasks, for each landscape. Most of these tasks are designed to mimic common, real-world data collection scenarios and are thus a more realistic assessment of generalization than random train-test splits. However, random splits are also included as a point of reference. We chose 8 of the 15 FLIP tasks to benchmark the panel of uncertainty methods. We selected these tasks to be representative of several regimes of domain shift – random sampling with no domain shift (AAV/Random, Meltome/Random, and GB1/Random); the highest (and most relevant) domain-shift regimes (AAV/Random vs. Designed and GB1/1 vs. Rest); and less aggressive domain shifts (AAV/7 vs. Rest, GB1/2 vs. Rest and GB1/3 vs. Rest). The Datasets section of the Methods provides notes on the nomenclature used for these tasks.

We trained the seven models on each of the eight tasks described above and evaluated their performance on the test set using the metrics described in Section 4.7. We compare model calibration and accuracy in Figure 2 and the percent coverage versus average width relative to range in Figure 3. These figures illustrate the results for models trained on the embeddings from a pretrained ESM language model (*15*); the corresponding results using one-hot encodings are shown in Figures S1 and S2.

**Figure 2:**
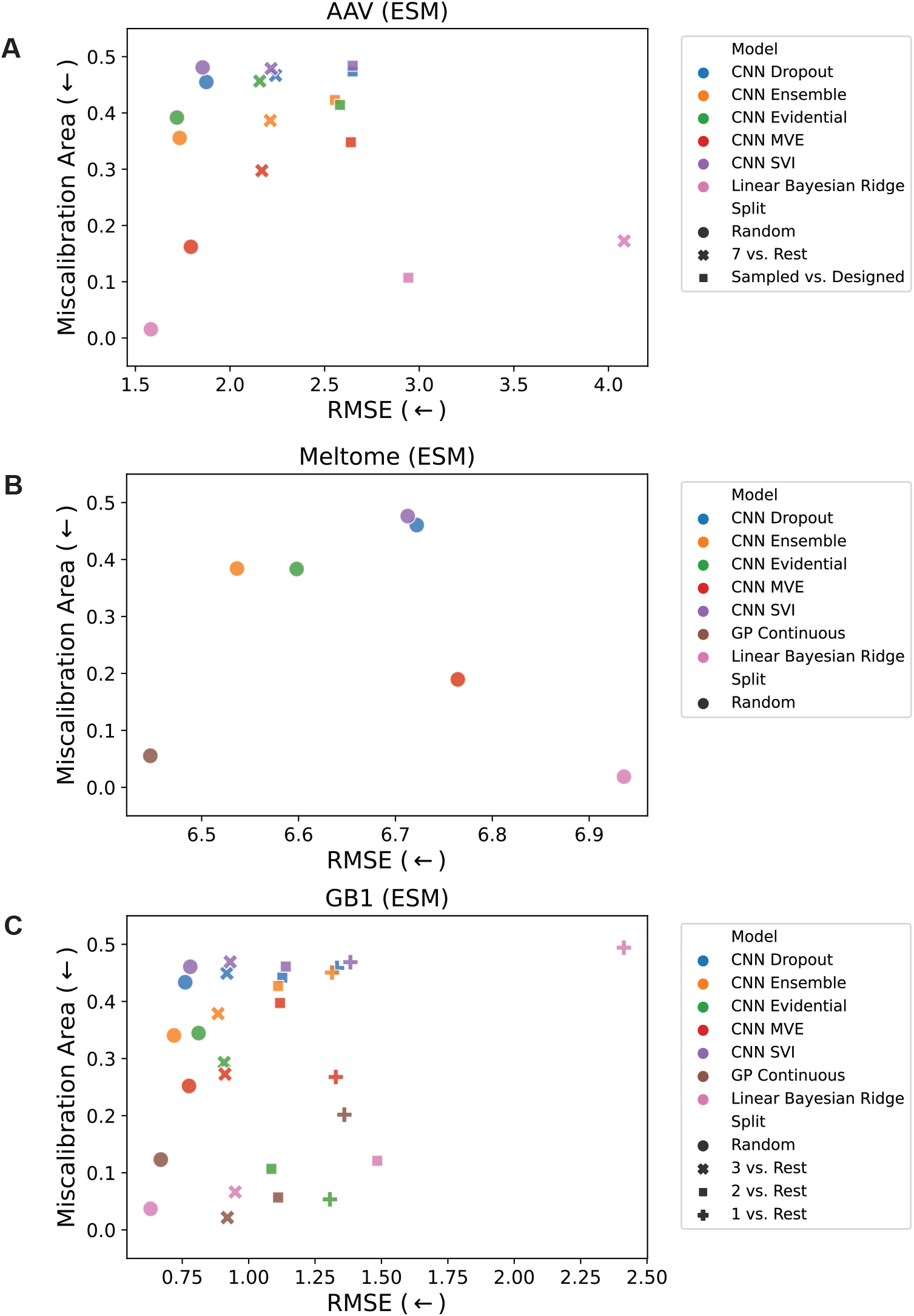
Miscalibration area vs. root mean square error (RMSE) for the (A) AAV, (B) Meltome, and (C) GB1 landscapes. Miscalibration area (also called the area under the calibration error curve or AUCE) quantifies the absolute difference between the calibration plot and perfect calibration. It is desirable to have a model that is both accurate and well-calibrated, so the best performing points are those closest to the lower left corner of the plots. Each point represents an average of 5 models trained using different random seeds for initialization of the CNN parameters and batching / stochastic gradient descent. The GP Continuous model is not shown for the AAV landscape due to memory constraints for training these models. Figure S1 shows the corresponding results for the OHE representation.

**Figure 3:**
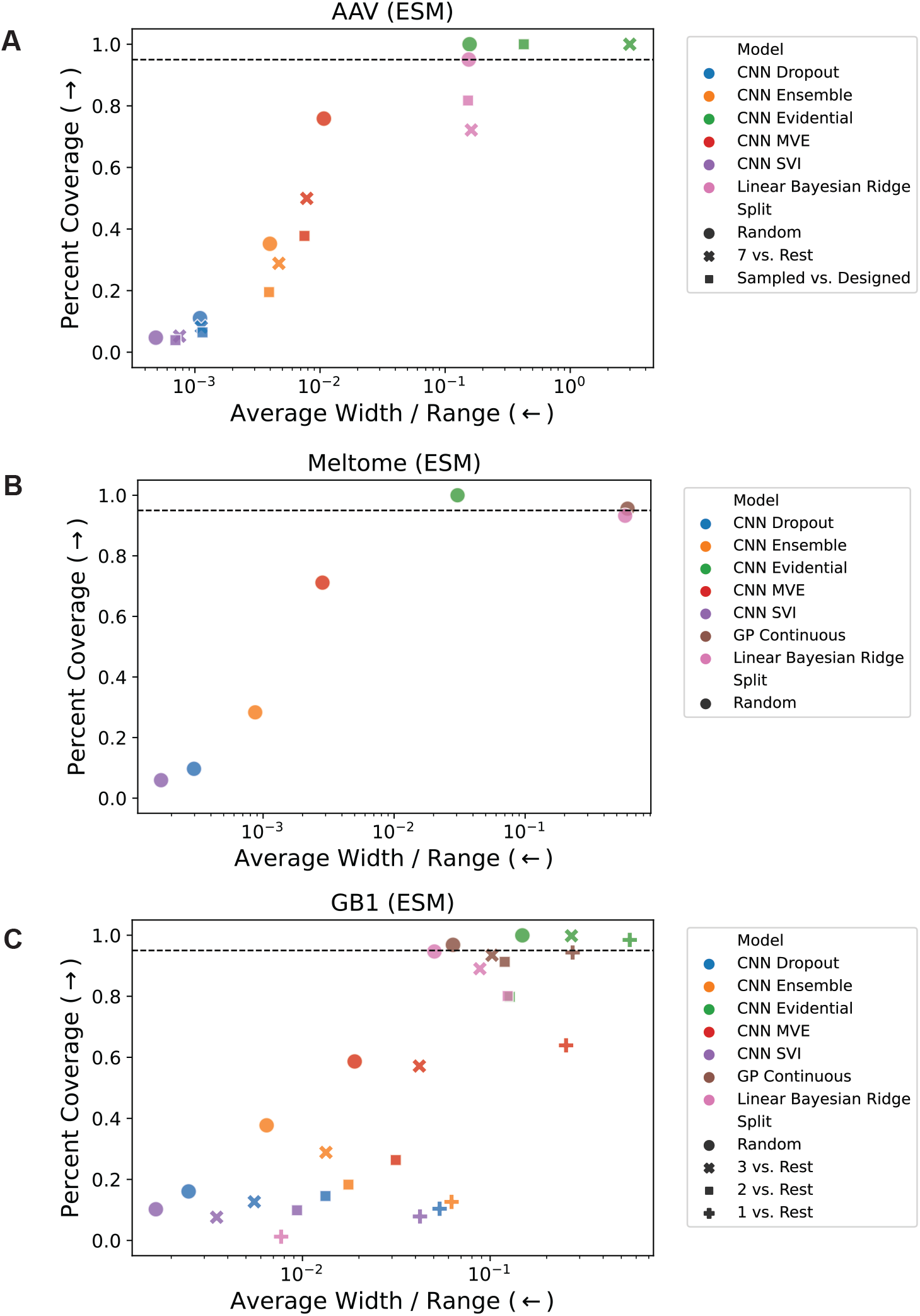
Coverage vs. average width / range for the (A) AAV, (B) Meltome, and (C) GB1 landscapes. Coverage is the percentage of true values that fall within the 95% confidence interval (2σ) of each prediction, and the width is the size of the 95% confidence region relative to the range of the training set (4σ/R where R is the range of the training set). A good model exhibits high coverage and low width, which corresponds to the upper left of each plot. The horizontal dashed line indicates 95% coverage. Each point represents an average of 5 models trained using different random seeds for initialization of the CNN parameters and batching / stochastic gradient descent. The GP Continuous model is not shown for the AAV landscape due to memory constraints for training these models. Figure S2 shows the corresponding results for the OHE representation.

As expected, the splits with the least required domain extrapolation tend to have more accurate models (lower RMSE; Fig. 2). However, the relationship between miscalibration area and extrapolation is less clear; some models are highly calibrated on the most difficult (highest domain shift) splits, while others are poorly calibrated even on random splits. There is no single method that performs consistently well across splits and landscapes, but some trends can be observed. For example, ensembling is often one of the highest accuracy CNN models, but also one of the most poorly calibrated. Additionally, GP and BRR models are often better calibrated than CNN models. For the AAV and GB1 landscapes (Fig. 2a, c), model miscalibration area usually increases slightly while RMSE increases more substantially with increasing domain shift.

In addition to accuracy and calibration, we assessed each method in terms of the coverage and width of its uncertainty estimates. A good uncertainty method results in high coverage (a large percentage of points where the true value falls within the 95% confidence region established by the uncertainty) while still maintaining a small average width. The latter is necessary because predicting a very large and uniform value of uncertainty for every point would result in good coverage, so coverage alone is not sufficient. Figure 3 illustrates that many methods perform relatively well in either coverage or width (corresponding to the the top and left limits of the plot, respectively), but few methods perform well in both. Similarly to Figure 2, there is some observable trend that more challenging splits are further from the optimal part (upper left) of the plot; this trend is more clear for the GB1 splits (Fig. 3b) than for the AAV splits. Most models trained on the AAV landscape (Fig. 3a) have a similar average width/range ratio for all splits, but for the GB1 landscape (Fig. 3c), this ratio typically increases as the domain shift increases. The locations of the sets of points for each model type shared some similarities across landscapes. CNN SVI often has low coverage and low width, CNN MVE often has moderate coverage and moderate width, and CNN Evidential and BRR often have high coverage and high width. The results for all prediction and uncertainty metrics are shown in Tables S1-S22.

We next assessed how target predictions and uncertainty estimates depended on the degree of domain shift. Across datasets and splits, we compared the ranking performance of each method in terms of predictions relative to true values and uncertainty estimates relative to true errors (ESM in Figure 4 and OHE in Figure S3). The splits are ordered according to domain shift within their respective landscapes (lowest to highest shift from left to right). We observe that the rank correlation of the predictions to the true labels generally decreases moving from less to more domain shift within a landscape, consistent with expectation, with the exception of AAV/Random vs. Designed models performing better than AAV/7 vs. Rest models (Fig. 4a). Most methods exhibit similar performance in 𝜌 within the same task. For many tasks, GP and BRR models perform as well or better than CNN models. Performance on 𝜌*_unc_* is generally much worse than that on 𝜌, with some results showing negative correlation (Fig. 4b). MVE and evidential uncertainty methods are most performant in 𝜌*_unc_* for most cases of low to moderate domain shift. Most methods have 𝜌*_unc_* near zero for the most challenging splits. Despite the relatively good performance of MVE on tasks with low to moderate domain shift, it performs poorly in cases of high domain shift, which is consistent with its intended use as an estimator of aleatoric (data-dependent) uncertainty.

**Figure 4:**
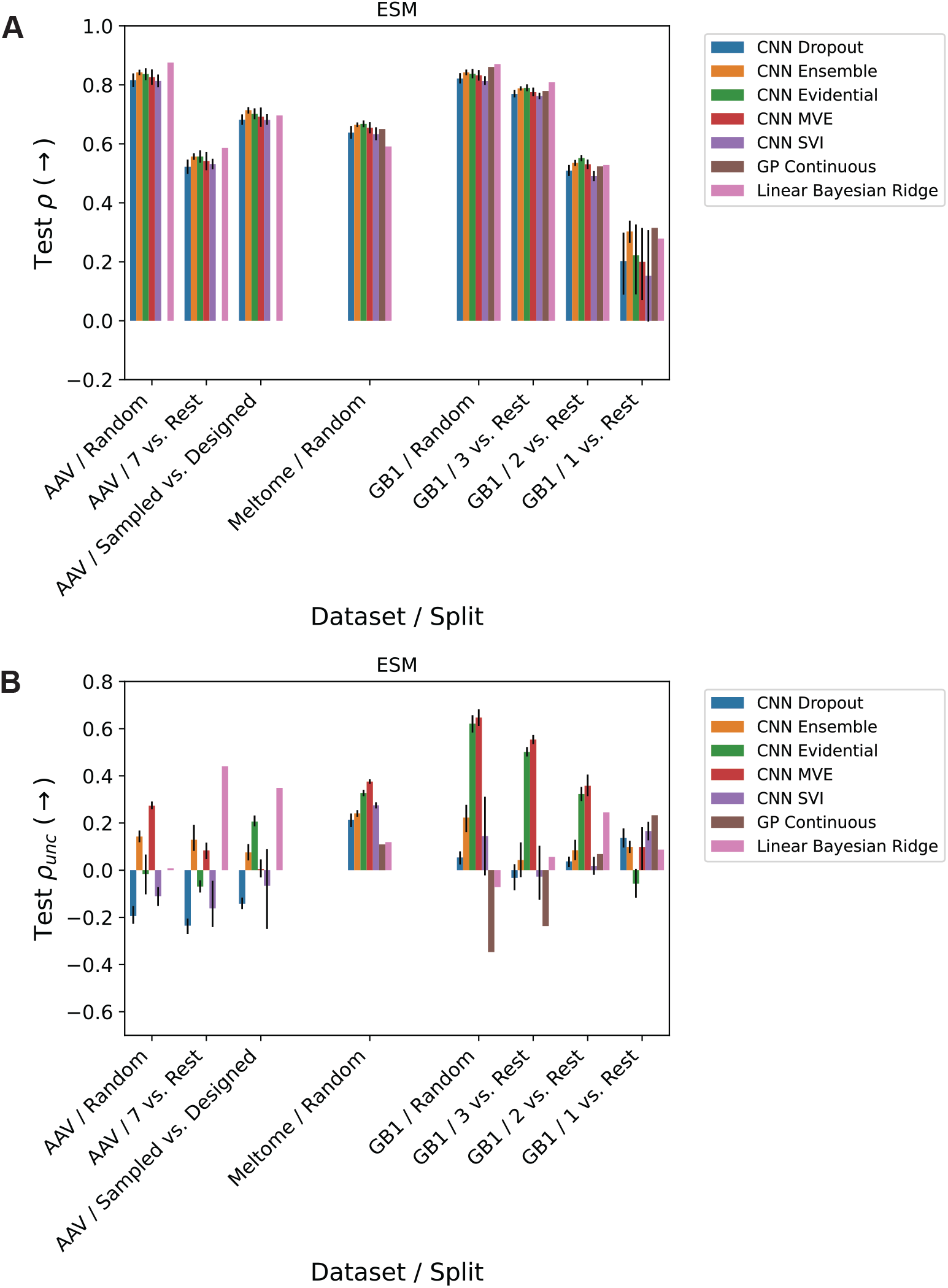
Spearman rank correlations of (A) predictions (𝜌) and (B) uncertainties (𝜌*_unc_*) vs. extrapolation. Within each landscape (AAV, Meltome, and GB1), splits are qualitatively ordered by the amount of domain shift between train and test sets, with the lowest domain shift on the left and the highest domain shift on the right. Error bars on the CNN results represent the 95% confidence interval calculated from 5 different random seed for initialization of the CNN parameters and batching / stochastic gradient descent. Figure S3 shows the corresponding results for the OHE representation.

We find that the models trained on ESM embeddings outperform those trained on one- hot encodings in 21 out of 51 cases for rank correlation of test set predictions, and 29 out of 51 cases for rank correlation of test set uncertainties. The relative performance of the two representations on prediction and uncertainty rank correlation is shown in Figure S4. In terms of predictions, ESM embeddings often yield substantially better performance for tasks with high domain shift (e.g. GB1/1 vs. Rest and Meltome/Random), while OHE performs slightly better on tasks with lower domain shift (e.g. AAV/Random and GB1/3 vs. Rest). The relative uncertainty rank correlation performance, on the other hand, does not have a clear relationship to domain shift.

### 2.2 Active Learning

In protein engineering, the purpose of uncertainty estimation is typically to intelligently prioritize sample acquisition for experimentation. One such use case of uncertainty is in active learning, where uncertainty estimates are used to inform sampling with the goal of improving model predictions overall (i.e., to achieve an accurate model with less training data; Fig. 5a). Having assessed the calibration and accuracy of the panel of UQ methods above, we next evaluated whether uncertainty-based active learning could make the learning process more sample-efficient. Across all datasets and splits using the pretrained ESM embeddings, data acquisition was simulated as iterative selection from the data library according to a given sampling strategy (acquisition function; see Methods for details). The results are summarized in Figure 5 for Spearman rank correlation (⇢) on three methods and one split per landscape, and additional results are shown in the Figures S5-S57 for other metrics, uncertainty methods, and splits. Across most models, the performance difference between the start of active learning (10% of training data) and end of active learning (100% of training data) is relatively small, and many models begin to plateau in performance before reaching 100% of training data.

**Figure 5:**
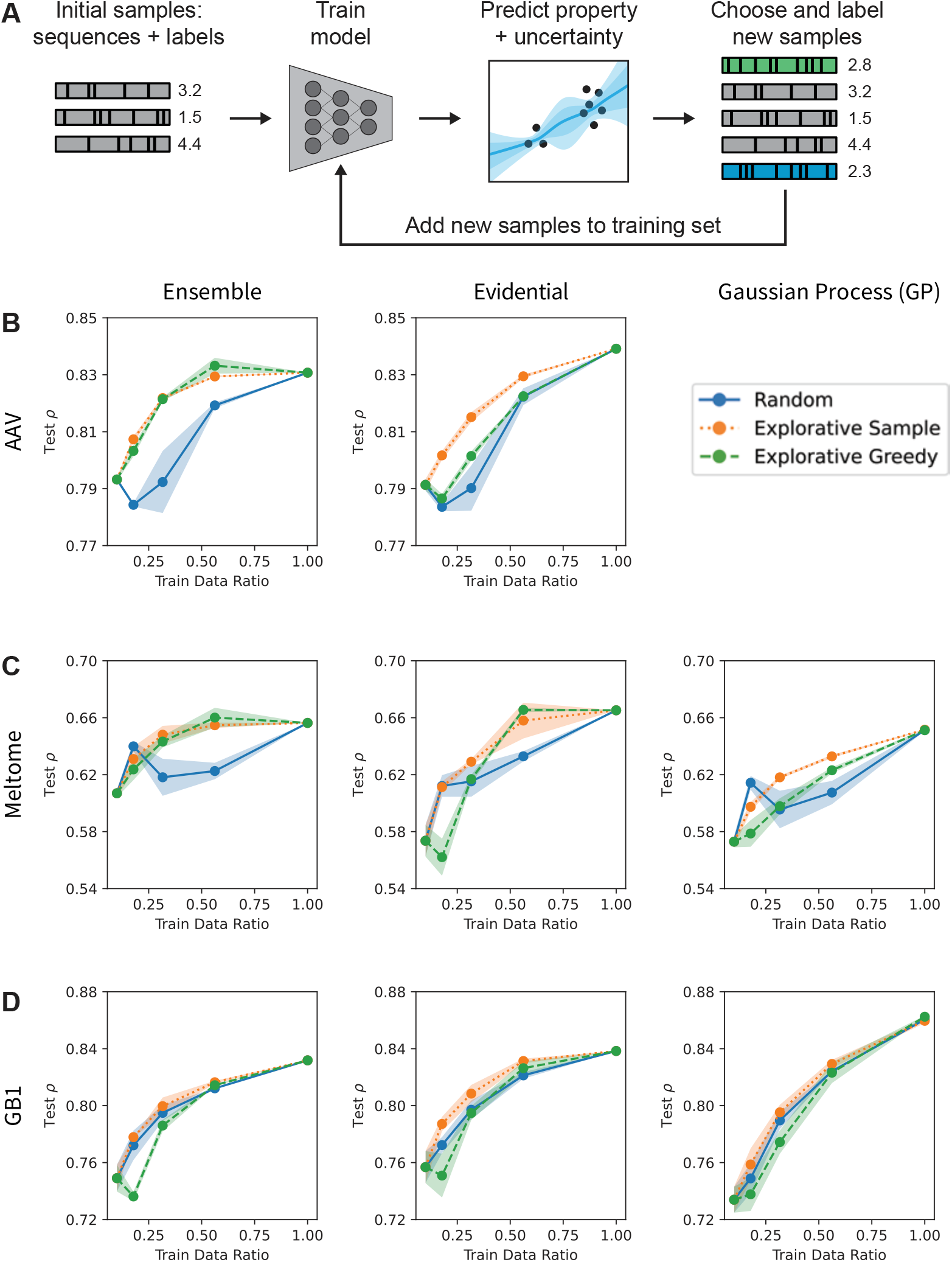
(A) Schematic of active learning approach. A model is trained on an initial dataset, and is then retrained in each iteration by adding more points to the training set based on some selection criteria. (B-D) Uncertainty-guided active learning in protein sequence-function prediction. Spearman rank correlation of predictions (⇢) for the CNN ensemble, CNN evidential, and GP methods evaluated on the AAV/Random (B), Meltome/Random (C), and GB1/Random (D) splits. The “random” strategy acquired sequences with all unseen points having equal probabilities, the “explorative sample” strategy acquired sequences with random sampling weighted by uncertainty, and the “explorative greedy” strategy acquired the previously unseen sequences with the highest uncertainty.

The “explorative greedy” and “explorative sample” acquisition functions (which sample based on uncertainty alone or sample randomly weighted by uncertainty, respectively) sometimes significantly outperform random sampling, but this is not true across all methods and landscapes (Fig. 5b-d). In some cases, the performance of the uncertainty-based sampling strategies also varies depending on the fraction of the total training data available to the model. For example, for the Meltome/Random split and CNN evidential model (Fig. 5c), explorative greedy sampling results in a *decrease* in model performance after the first round of active learning while the explorative sample strategy increases performance. By the fourth round of active learning for this task, the two explorative strategies significantly outperform random sampling. This indicates that in the early stages of active learning when a model’s uncertainty estimates are poorly calibrated, it may be advantageous to sample with at least some randomness included in an uncertainty-based acquisition function. Overall, the results indicate that uncertainty-informed active learning can outperform random sampling and thus lead to more accurate machine learning models with fewer training points needing to be measured (Fig. 5b-d).

## 3 Conclusions

Calibrated uncertainty estimations for ML predictions of biomolecular properties are necessary for effective model improvement using active learning or property optimization using Bayesian methods. In this work, we benchmarked a panel of uncertainty quantification (UQ) methods on protein datasets, including on train-test splits that are representative of realworld data collection practices. After evaluating each method based on accuracy, calibration, coverage, width, rank correlation, and performance in active learning, we find that there is no method that performs consistently well across all metrics or all landscapes and splits.

We also examined how models trained using one-hot-encoding representations of sequences compare to those trained on more informative and generalizable representations such as embeddings from a pretrained ESM language model. This comparison illustrated that while the pretrained embeddings do improve model accuracy and uncertainty correlation/calibration in some cases, particularly on splits with higher domain shifts, this is not universally true and in some cases makes performance worse.

While the UQ evaluation metrics used in this work provide valuable information, they are ultimately only a proxy for expected performance in Bayesian optimization and active learning. We found that UQ evaluation metrics are not well-correlated with gains in accuracy from one active learning iteration to another on these datasets. This suggests that future work in UQ should include retrospective Bayesian optimization and/or active learning studies rather than relying on UQ evaluation metrics alone. Our retrospective active learning studies using holdouts of the training sets demonstrate that many of the uncertainty methods outperform random sampling baselines. In some of our experiments, we observe that the uncertainty-based sampling strategies perform worse than random sampling during the earliest stages of active learning, then perform better as a model’s accuracy and quality of uncertainty estimates improve in later stages.

Future work in this area could expand on methods (e.g. Bayesian neural networks (*24*) and conformal prediction (*25*)), metrics (e.g. sharpness (*5*), dispersion (*26*), and tightness (*27*)), and representations (e.g. ESM-2 (*28*) or using an attention layer rather than mean aggregation on our ESM-1b embeddings). While this work considered uncertainty predictions as directly output by the models, further study is needed to understand the effects of posthoc calibration methods (e.g. scalar recalibration (*26*) or CRUDE (*29*)). Future work should consider additional active learning strategies beyond “explorative greedy” and “explorative sample”, such as Thompson sampling (*30*), other exploitative strategies, strategies that consider batch diversity in the acquisition function (*31*), and methods that consider the desired domain shift. Ultimately, this work contributes to a more thorough understanding of how to best apply UQ to sequence-function models and provides a foundation for future work to enable more effective protein engineering.

## 4 Methods

### 4.1 Regression Tasks

All tasks studied in this work are regression problems, in which we attempt to fit a model to a dataset with 𝒟 data points (𝑥_𝑖_, 𝑦_𝑖_). 𝑥_𝑖_ is a protein sequence representation (either a one-hot encoding or an embedding vector from an ESM language model), and 𝑦_𝑖_ ɛ ℝis a scalar-valued target property from the protein landscapes described in Section 4.2.

### 4.2 Datasets

The landscapes and splits in this work are taken from the FLIP benchmark (*3*). GB1 is a landscape commonly used for investigating epistasis (interactions between mutations) using the binding domain of protein G, an immunoglobulin binding protein in Streptococcal bacteria. These splits are designed primarily to test generalization from few-to many-mutation sequences. The AAV landscape is based on data collected for the Adeno-associated virus capsid protein, which help the virus integrate a DNA payload into a target cell. The mutations in this landscape are restricted to a subset of positions within a much longer sequence. The Meltome landscape includes data from proteins across 13 different species for a non-protein-specific property (thermostability), so it includes both local and global variations. The total number of data points in the GB1, AAV, and Meltome sets are 8,733, 284,009, and 27,951, respectively. In the AAV set, 82,583 are sampled (mutations) and 201,426 are designed. For AAV, only the 82,583 sampled sequences are used for the Random and 7 vs. Rest tasks, while all 284,009 are used for the Sampled vs. Designed task.

The names of several of the tasks were changed slightly from the original FLIP nomenclature for clarity: GB1/Random was originally called GB1/Sampled, AAV/Random was originally called AAV/Sampled, AAV/7 vs. Rest was originally called AAV/7 vs. Many, AAV/Sampled vs. Designed was originally called AAV/Mut-Des, and Meltome/Random was originally called Meltome/Mixed.

### 4.3 ESM Embeddings

We used the pretrained, 650M-parameter ESM-1b model (esm1b t33 650M UR50S) from (*15*) to generate embeddings of the protein sequences in this study and to compare these embeddings to one-hot encoding representations. Sequence embeddings from the final representation layer (layer 33) were mean pooled per amino acid over the length of each protein sequence, which resulted in a fixed embedding size of 1280 for each sequence.

### 4.4 Base CNN Model Architectures

The base architecture of all CNN models in this work was taken from the CNNs in the FLIP benchmark (*3*). For the one-hot encoding inputs, this was comprised of a convolution with 1024 channels and kernel width 5, a ReLU non-linear activation function, a linear mapping to 2048 dimensions, a max pool over the sequence, and a linear mapping to 1 dimension. For ESM embedding inputs, the architecture was the same except with 1280 input channels rather than 1024, and a linear mapping to 2560 dimensions rather than 2048.

### 4.5 CNN Model Training Procedures

To train our CNN models, we used a batch size of 256 (GB1, AAV) or 30 (Meltome). Adam (*32*) was used for optimization with the following learning rates: 0.001 for the convolution weights, 0.00005 for the first linear mapping, and 0.000005 for the second linear mapping. Weight decay was set to 0.05 for both the first and second linear mappings. CNNs were trained with early stopping using a patience of 20 epochs. Each model was trained on an NVIDIA Volta V100 GPU. Code, data, and instructions needed to reproduce results can be found at https://github.com/microsoft/protein-uq.

### 4.6 Uncertainty Methods

For all models and landscapes, the sequences were featurized using either one-hot encodings or embeddings from a pretrained language model (see Section 4.3).

We used the scikit-learn (*33*) implementation of Bayesian ridge regression (BRR) with default hyperparameters. BRR for one-hot encodings of the Meltome/Random split was not feasible because the required work array was too large to perform the computation with standard 32-bit LAPACK in scipy.

For Gaussian processes (GPs), we used the GPyTorch (*34*) implementation with the constant mean module, scaled rational quadratic (RQ) kernel covariance module, and Gaussian likelihood. Some GP models (for AAV one-hot encodings and ESM embeddings, and Meltome one-hot encodings) were not feasible to train due to GPU-memory requirements for exact GP models, so these are omitted from the results.

For our uncertainty methods that rely on sampling (dropout, ensemble, and SVI), the final model prediction is defined as the mean of the set of inference samples, and the uncertainty is the standard deviation of these samples. In other words, for a set of predictions 𝜀 = {𝐺_1_(𝑥), 𝐺_2_(𝑥), …, 𝐺_𝑛_(𝑥)} (each coming from an individual model 𝐺_𝑖_), the final prediction is defined as

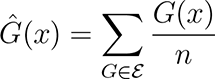

and the uncertainty 𝑈(𝑥) is defined as

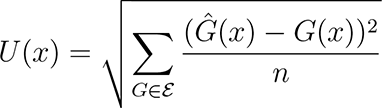

The uncertainty is sometimes defined as the variance 𝑈^2^, but using the standard deviation puts the uncertainty in the same units as the predictions.

For dropout uncertainty (*19*), a single model 𝐺 was trained normally. At inference time, we applied 𝑛 = 10 random dropout masks with dropout probability 𝑝 to obtain the set of predictions 𝜀 for each input 𝑥_𝑖_. We tested dropout rates of 𝑝 ɛ {0.1, 0.2, 0.3, 0.4, 0.5} and reported the model with the lowest miscalibration area.

Similarly for last-layer stochastic variational inference (SVI) (*23*), we obtained 𝜀 using 𝑛 = 10 samples from a set of models where each 𝐺_𝑖_ has the weight and bias terms of its last layer themselves sampled from a distribution 𝑞(𝜃) that has been trained to approximate the true posterior 𝑝(𝜃|𝒟).

Traditional model ensembling calculated 𝜀 using 𝑛 = 5 models trained using different random seeds for initialization of the CNN parameters and batching / stochastic gradient descent. The computational cost of this approach is 5 times that of a standard CNN model since the cost scales linearly with the size of the ensemble.

In mean-variance estimation (MVE) models, we adapt the base CNN architecture to produce 2 outputs (𝜃 = {𝜇,𝜎^2^}) for each data point (𝑥_𝑖_, 𝑦_𝑖_) in the last layer rather than 1, and we train using the negative log-likelihood loss:

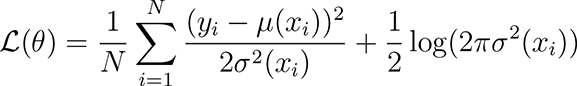

In practice, the variance (𝜎^2^) is clamped to a minimum value of 10*^—^*^6^ to prevent division by 0.

Evidential deep learning modifies the loss function of the traditional CNN to jointly maximize the model’s fit to data while also minimizing its evidence on errors (increasing uncertainty on unreliable predictions) (*21*):

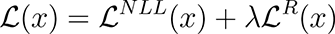

where 𝐿^𝑁𝐿𝐿^(𝑥) is the negative log-likelihood, 𝐿^𝑅^(𝑥) is the evidence regularizer, and λ controls the trade-off between these two terms. In this study, we use λ = 1 for all evidential models. In these models, the last layer of the model produces 4 outputs **m** = {𝛾, 𝜐, 𝛼, 𝛽 } that parameterize the Normal-Inverse-Gamma distribution. This distribution assumes that targets 𝑦_𝑖_ are drawn i.i.d. from a Gaussian distribution with unknown mean and variance (𝜃 = {𝜇,𝜎^2^}), where the mean is drawn from a Gaussian and the variance is drawn from an Inverse-Gamma distribution. The output of the evidential model can be divided into the prediction and the epistemic (model) and aleatoric (data) uncertainty components following the analysis of Amini et al. (*21*):

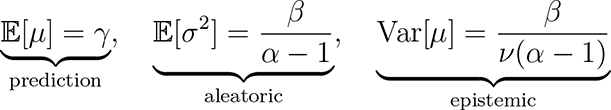

We report the sum of the aleatoric and epistemic uncertainties as the total uncertainty.

### 4.7 Evaluation Metrics

To give a comprehensive report of model accuracy, we computed the following metrics on the test sets: root mean square error (RMSE), mean absolute error (MAE), coefficient of determination (𝑅^2^), and Spearman rank correlation (𝜌). RMSE is more sensitive to outliers than MAE, so while both are informative independently, the combination of the two gives additional information about the distribution of errors. 𝑅^2^ and 𝜌 are both unitless and are thus more easily interpreted and compared across datasets.

We evaluated the quality of the uncertainty estimates using four metrics. First, 𝜌*_unc_* is the Spearman rank correlation between uncertainty and absolute prediction error. Following Kompa et al. (*35*), we measured the coverage as the percentage of true values that fall within the 95% confidence interval (±2σ) of each prediction. Kompa et al. (*35*) define the width as the size of the 95% confidence region (4σ), but we normalized this width relative to the range (𝑅) of the training set as 4σ/𝑅 to make these values unitless and thus more interpretable across datasets. Finally, the miscalibration area (also called the area under the calibration error curve or AUCE) quantifies the absolute difference between the calibration plot and perfect calibration in a single number (*36*).

### 4.8 Active Learning

Each active learning run began with a random sample of 10% of the full training data. We evaluated several alternatives for adding to this initial dataset using different sampling strategies (acquisition functions): explorative greedy, explorative sample, and random. “Explorative greedy” sampled the sequences with the highest uncertainty; “explorative sample” sampled the data according to the probability of sampling a sequence equal to the ratio of its uncertainty to the sum of all uncertainties in the dataset (i.e. random sampling weighted by uncertainty); and “random” sampled the data uniformly from all unobserved sequences. We employed these sampling strategies 5 times in each active learning run, with the 5 training set sizes equally spaced on a log scale. We repeated this process using 3 folds (different random seeds for sampling initial dataset and “explorative sample” probabilities) and calculated the mean and standard deviation across these folds.

## Acknowledgement

K.P.G. was supported by a Microsoft Research micro-internship and by the National Science Foundation Graduate Research Fellowship Program under Grant No. 1745302. The authors thank the MIT Lincoln Laboratory Supercloud cluster (*37*) at the Massachusetts Green High Performance Computing Center (MGHPCC) for providing high-performance computing resources to train our machine learning models.

## Abbreviations

ML, UQ, CNN, FLIP, GP, BO, BRR, MVE, SVI, AAV, GB1, RMSE, MAE, RQ

## 5 Author Contributions

A.P.A. and K.K.Y. conceived the project. K. P. G. wrote the computer code, analyzed the data, and wrote the first manuscript draft. A.P.A. and K.K.Y. supervised the research and edited the manuscript.

## Supporting Information Available

Code availability, OHE results, OHE vs. ESM results comparison, additional prediction and uncertainty evaluation metrics, and additional active learning results.

## Supporting Information Benchmarking Uncertainty Quantification for Protein Engineering

### 1 Code Availability

The code for the models, uncertainty methods, and evaluation metrics in this work is available at https://github.com/microsoft/protein-uq.

### 2 OHE Results

**Figure S1:**
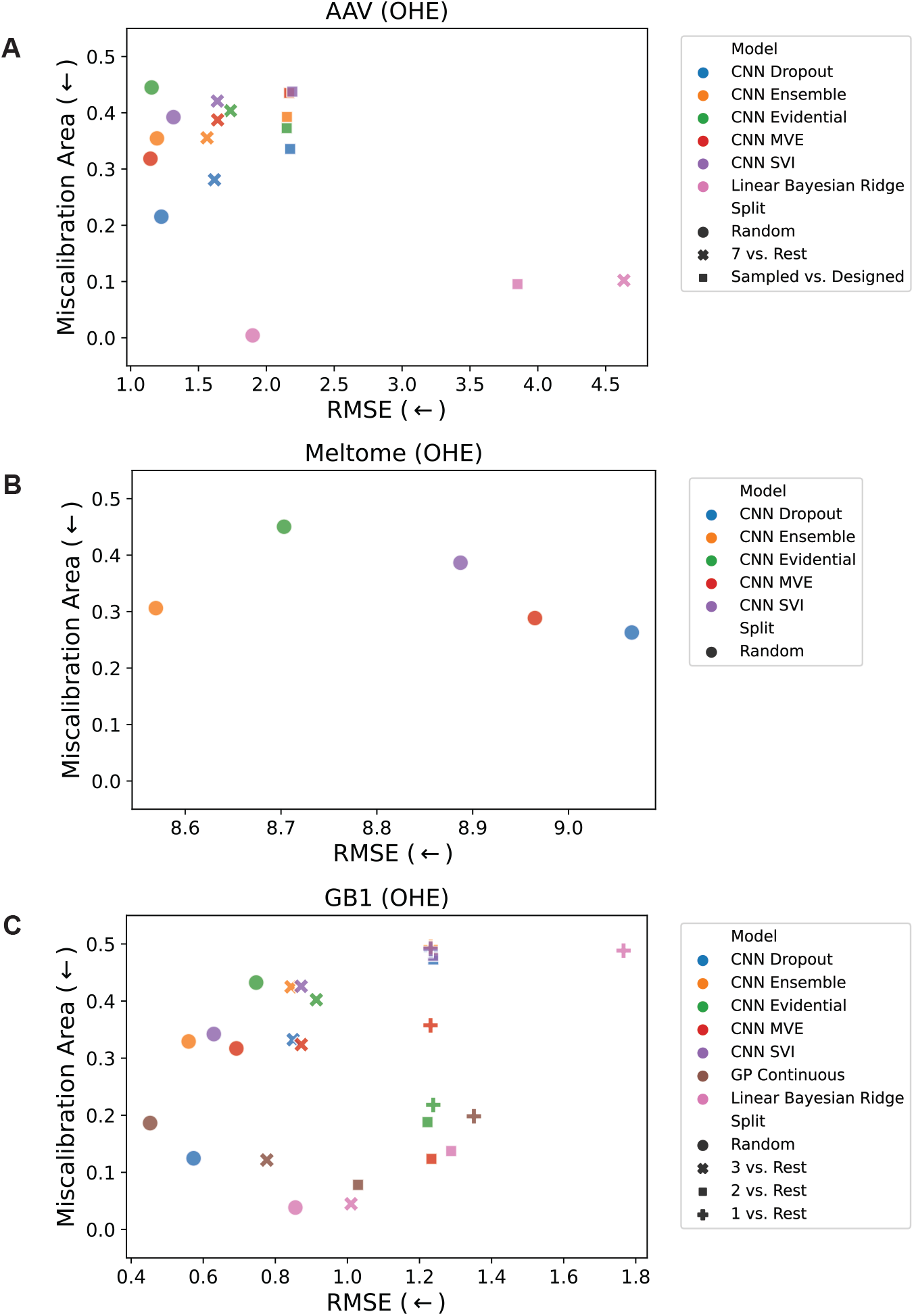
Miscalibration area vs. root mean square error (RMSE) for the (a) AAV, (b) Meltome, and (c) GB1 landscapes. Miscalibration area (also called the area under the calibration error curve or AUCE) quantifies the absolute difference between the calibration plot and perfect calibration. It is desirable to have a model that is both accurate and well-calibrated, so the best performing points are those closest to the lower left corner of the plots. The GP Continuous model is not shown for the AAV landscape due to memory constraints for training these models. The GP Continuous and Linear Bayesian Ridge models are not shown for the Meltome landscape due to memory constraints and limitations of 32-bit LAPACK, respectively.

**Figure S2:**
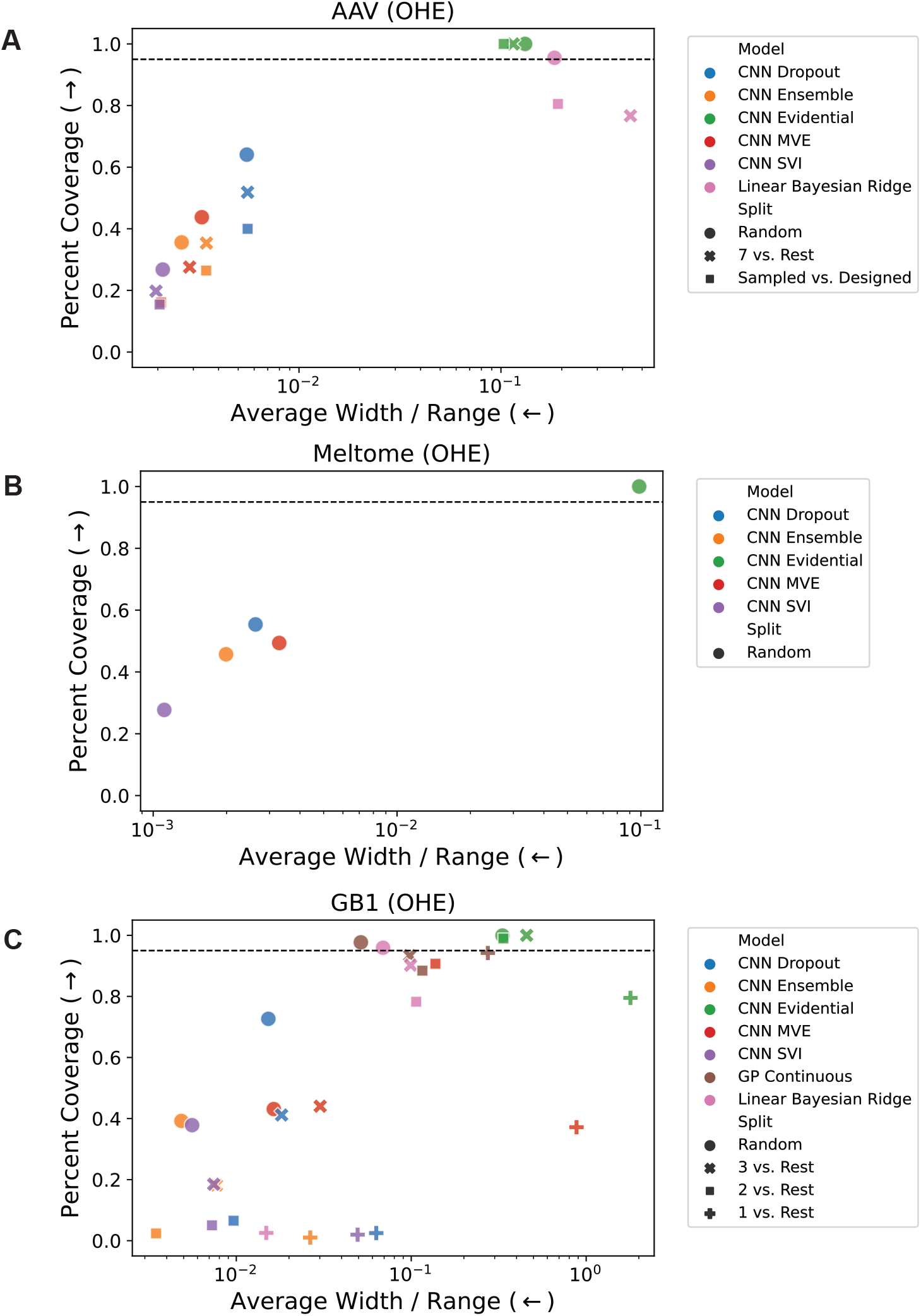
Coverage vs. average width / range for the (a) AAV, (b) Meltome, and (c) GB1 landscapes. Coverage is the percentage of true values that fall within the 95% confidence interval (±2𝜎) of each prediction, and the width is the size of the 95% confidence region relative to the range of the training set (4𝜎/𝑅 where 𝑅 is the range of the training set). A good model exhibits high coverage and low width, which corresponds to the upper left of each plot. The horizontal dashed line indicates 95% coverage. The GP Continuous model is not shown for the AAV landscape due to memory constraints for training these models. The GP Continuous and Linear Bayesian Ridge models are not shown for the Meltome landscape due to memory constraints and limitations of 32-bit LAPACK, respectively.

**Figure S3:**
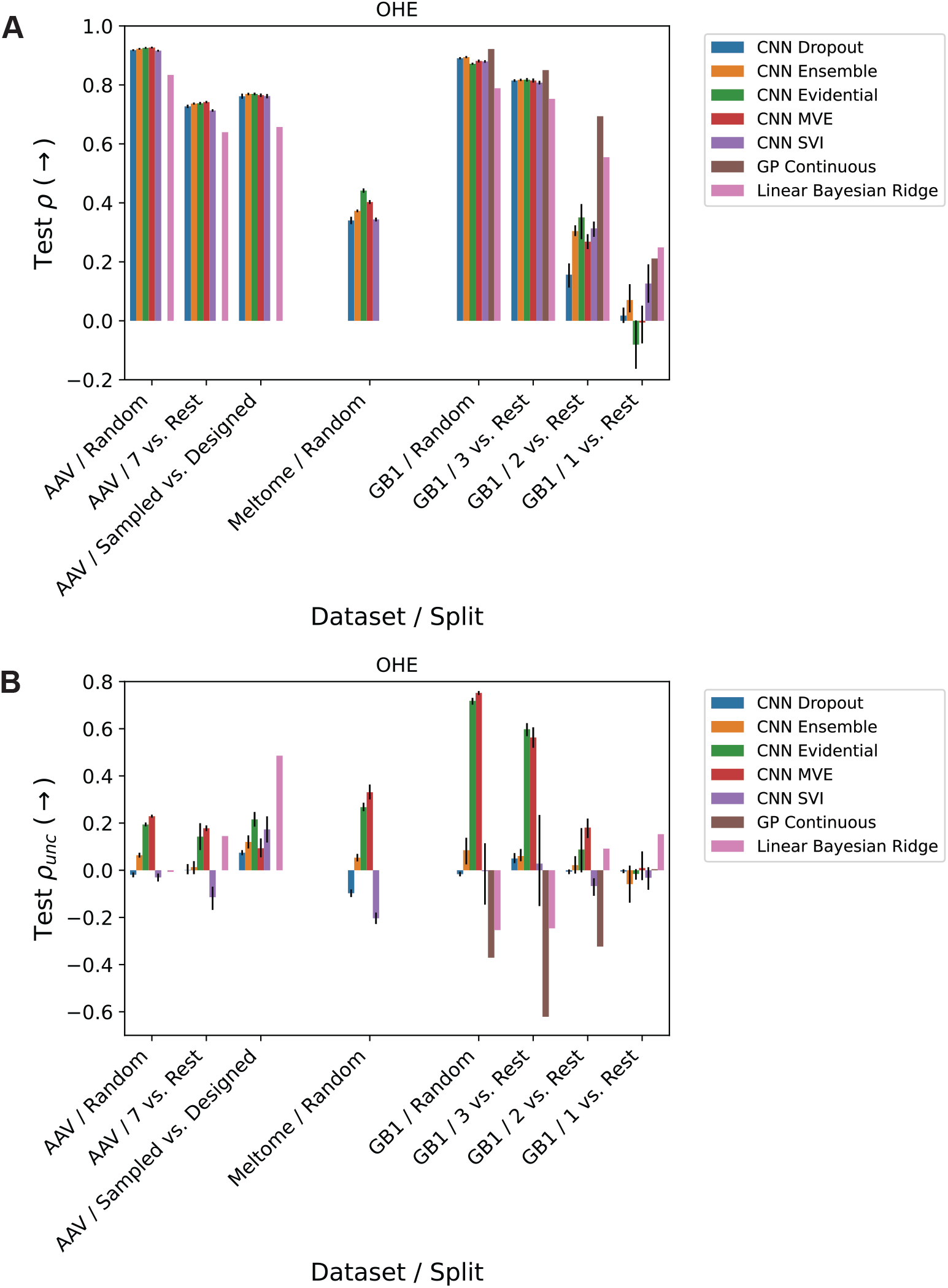
Spearman rank correlations of (a) predictions (𝜌) and (b) uncertainties (𝜌*_unc_*) vs. extrapolation. Within each landscape (AAV, Meltome, and GB1), splits are qualitatively ordered by the amount of domain shift between train and test sets, with the lowest domain shift on the left and the highest domain shift on the right. Error bars on the CNN results represent the 95% confidence interval calculated from 5 different random initializations of the CNN parameters.

### 3 OHE vs. ESM Comparison

**Figure S4:**
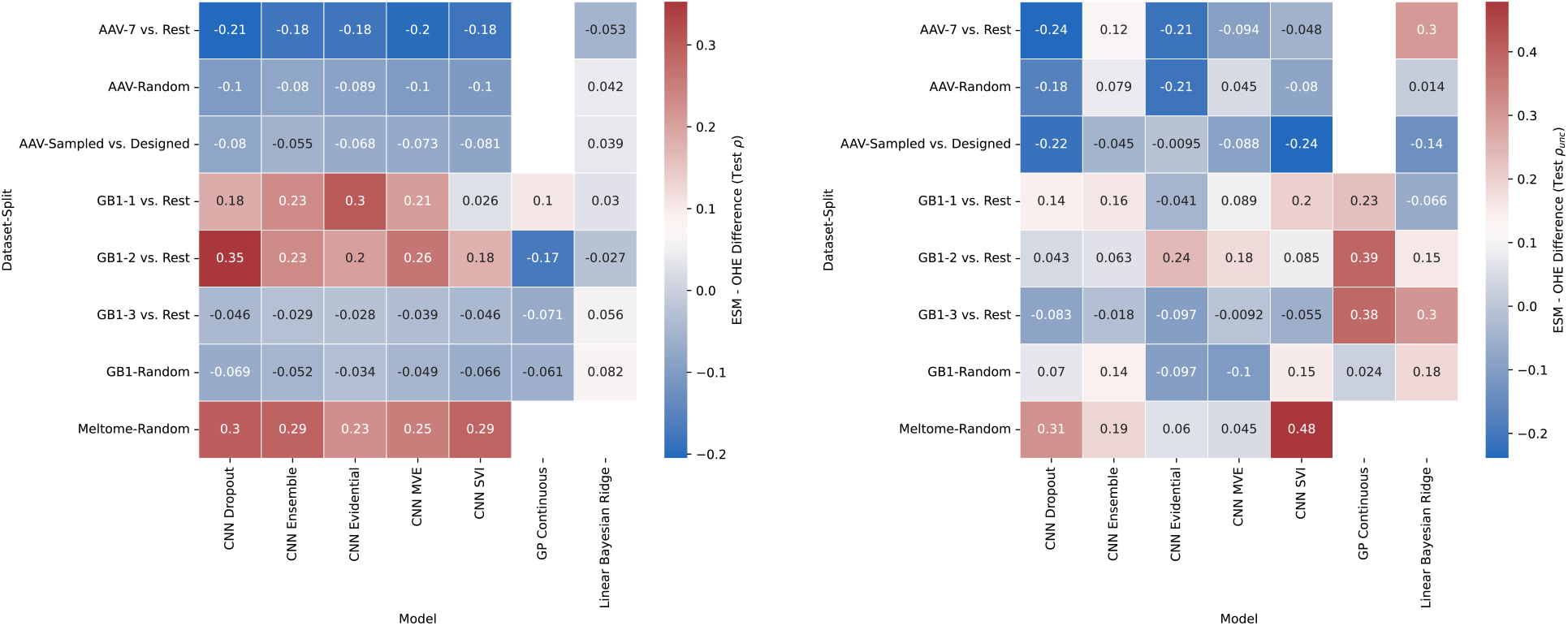
Comparison of prediction (𝜌) and uncertainty (𝜌*_unc_*) performance between the OHE and ESM representations across all models and tasks. Red cells indicate that the ESM representation performed better, while blue cells indicate that the OHE representation performed better.

### 4 Prediction and Uncertainty Evaluation Metrics

**Table S1:**
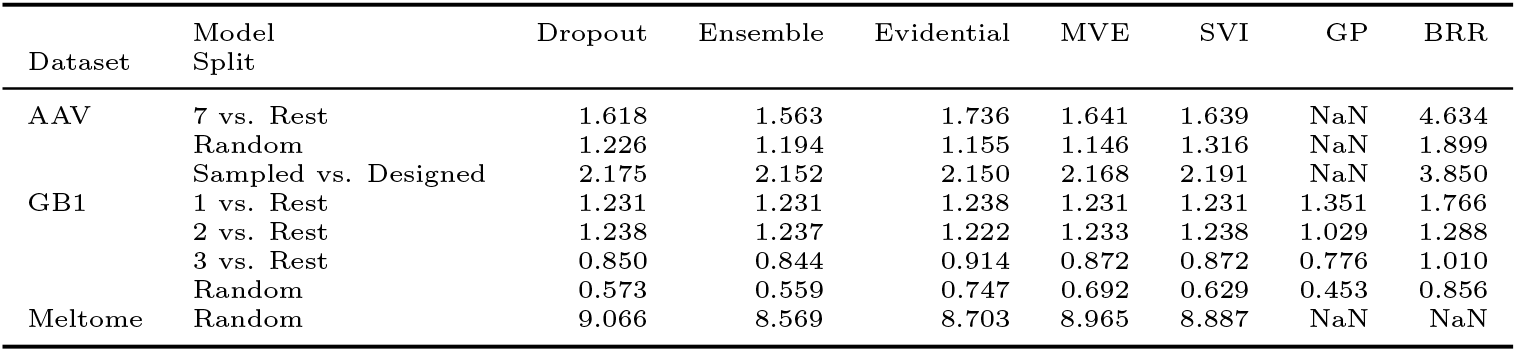
Test set RMSE for models trained on OHE representation (↓)

**Table S2:**
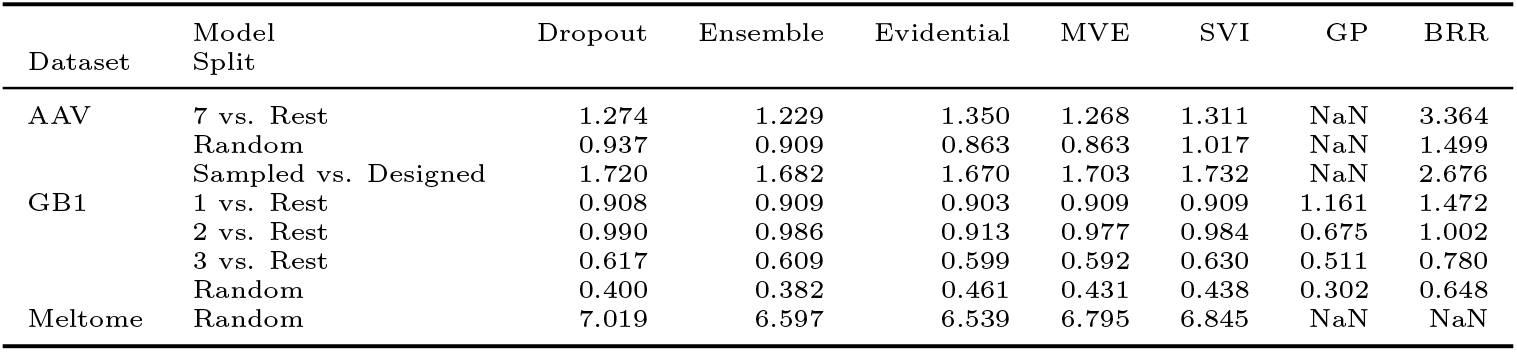
Test set MAE for models trained on OHE representation (↑)

**Table S3:**
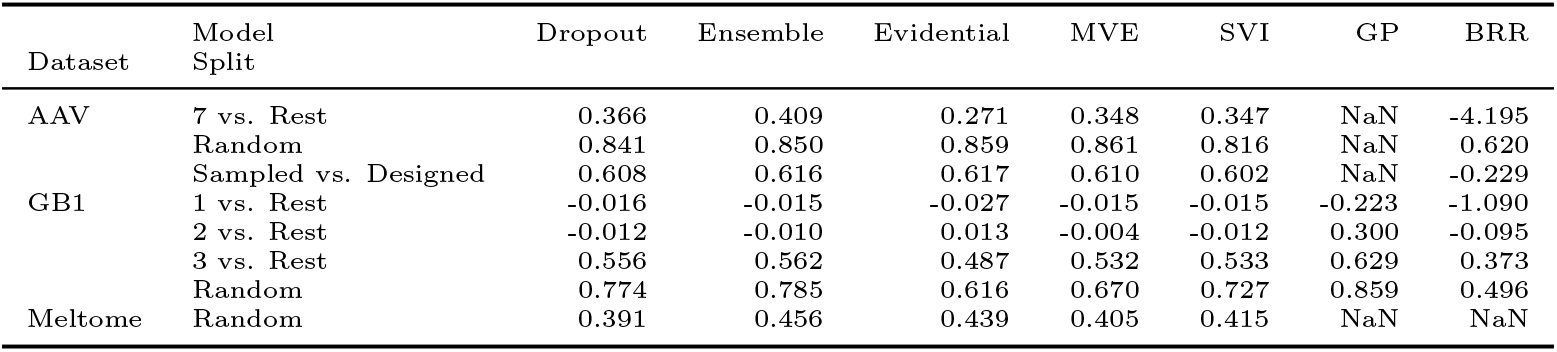
Test set 𝑅^2^ for models trained on OHE representation (↑)

**Table S4:**
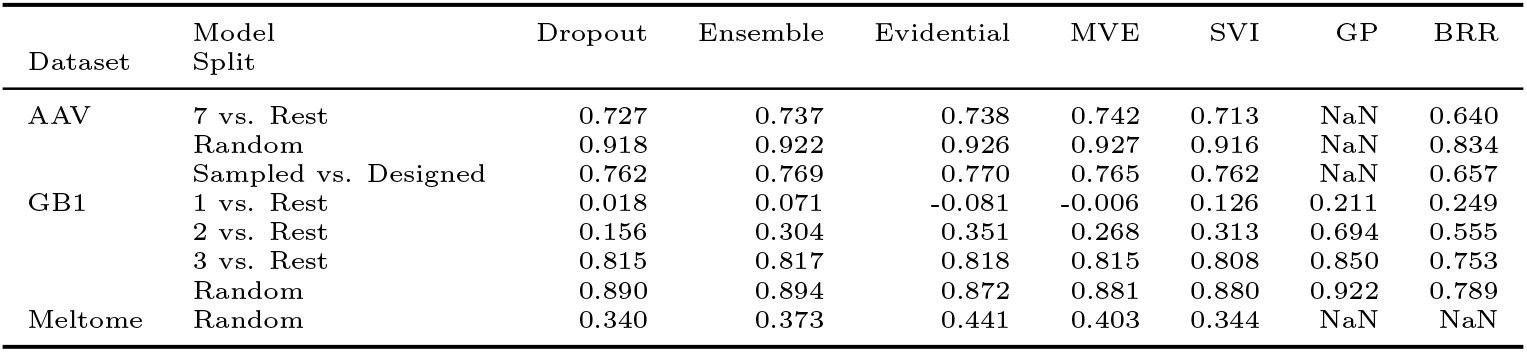
Test set 𝜌 for models trained on OHE representation (↑)

**Table S5:**
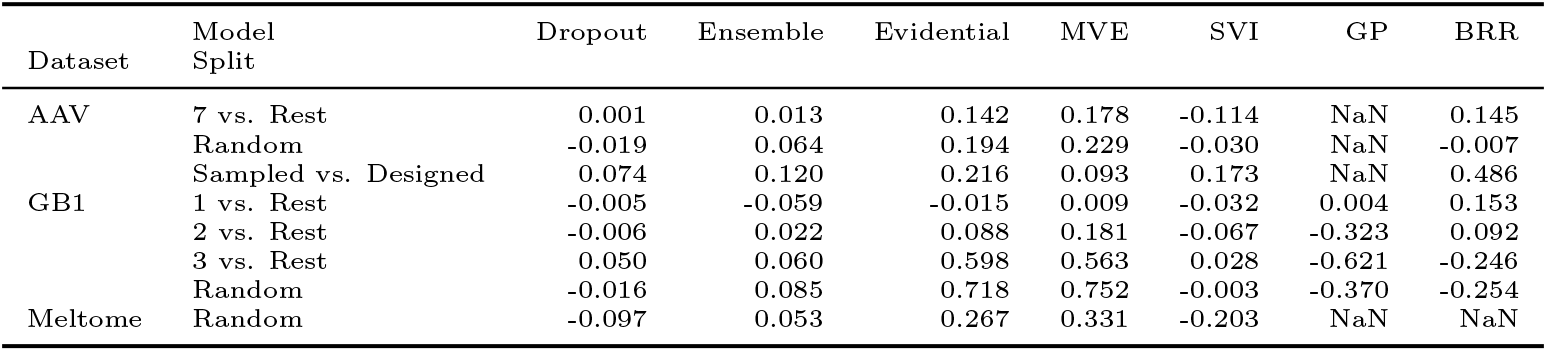
Test set 𝜌*_unc_* for models trained on OHE representation (↑)

**Table S6:**
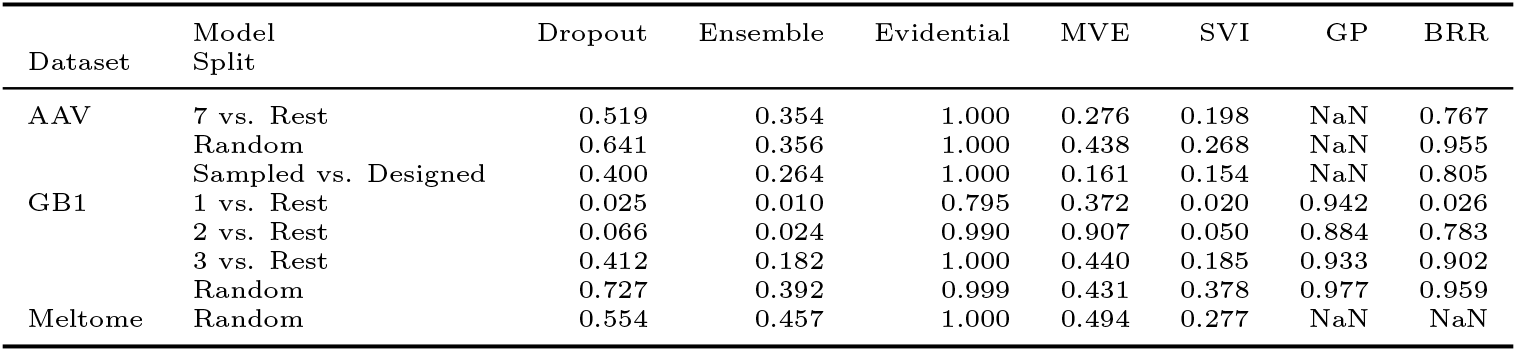
Test set % coverage for models trained on OHE representation (↑)

**Table S7:**
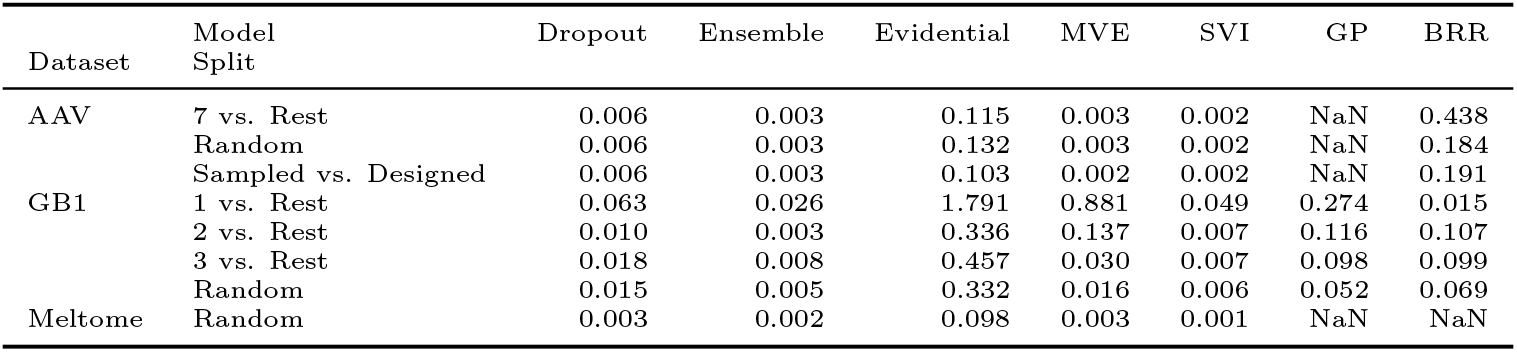
Test set 4𝜎/𝑅 for models trained on OHE representation (↓)

**Table S8:**
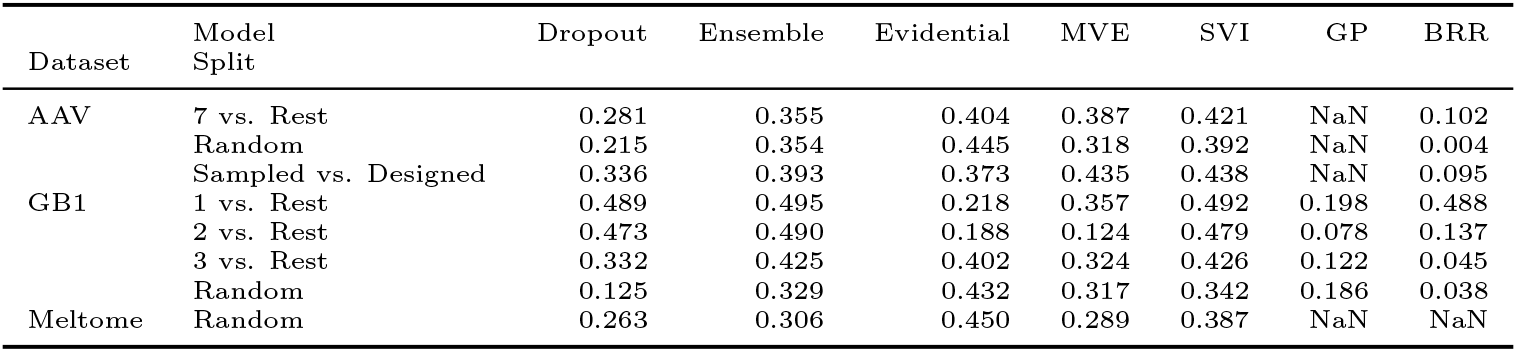
Test set miscalibration area for models trained on OHE representation (↓)

**Table S9:**
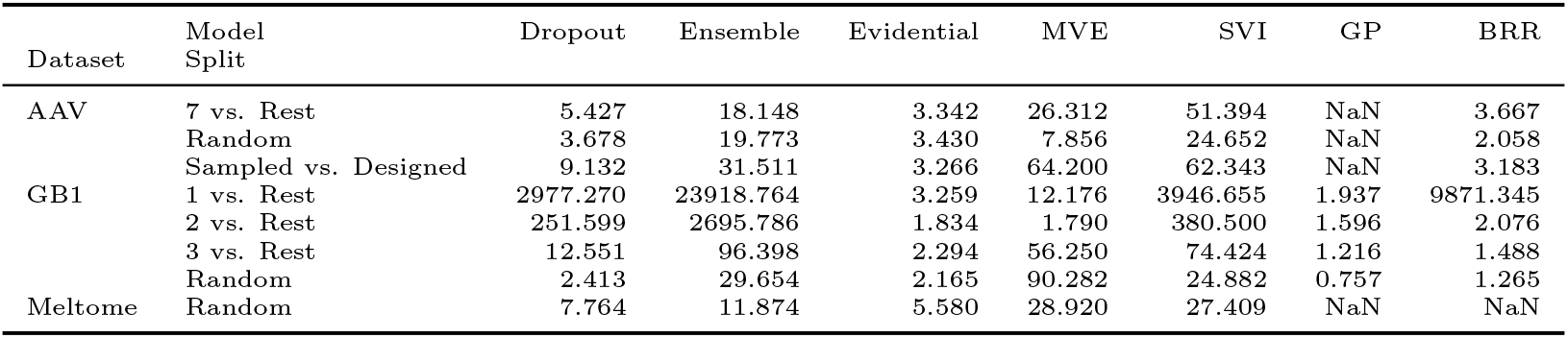
Test set 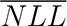 for models trained on OHE representation (↓)

**Table S10:**
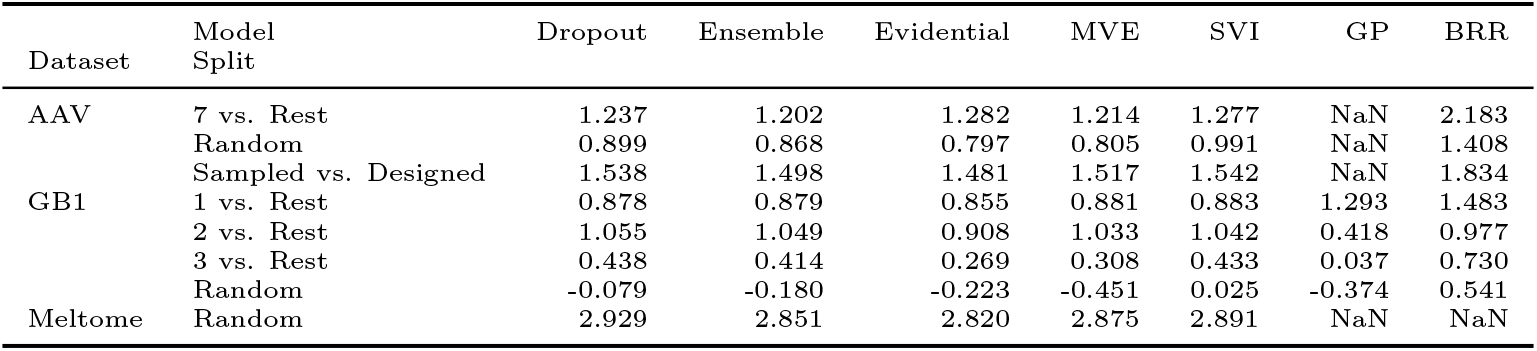
Test set 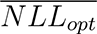 for models trained on OHE representation

**Table S11:**
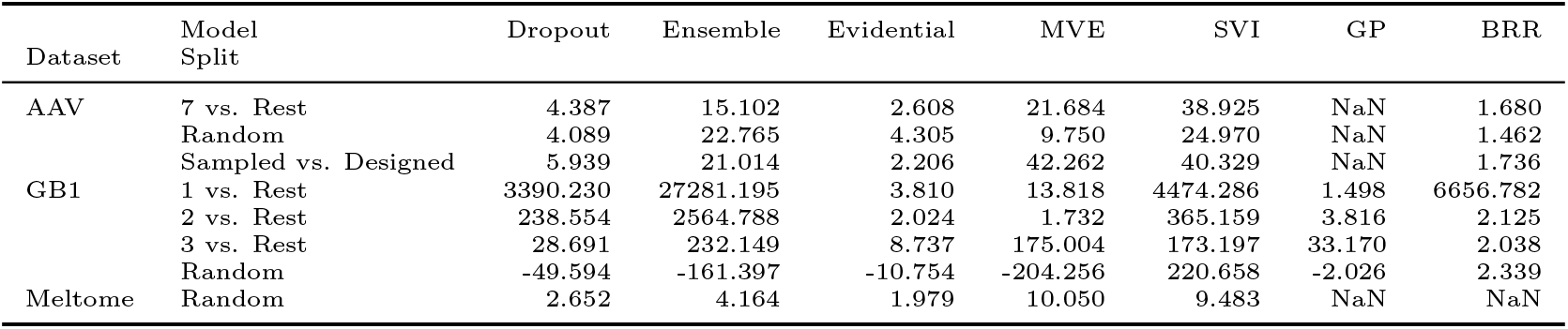
Test set 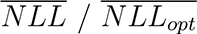 ratio for models trained on OHE representation (↓)

**Table S12:**
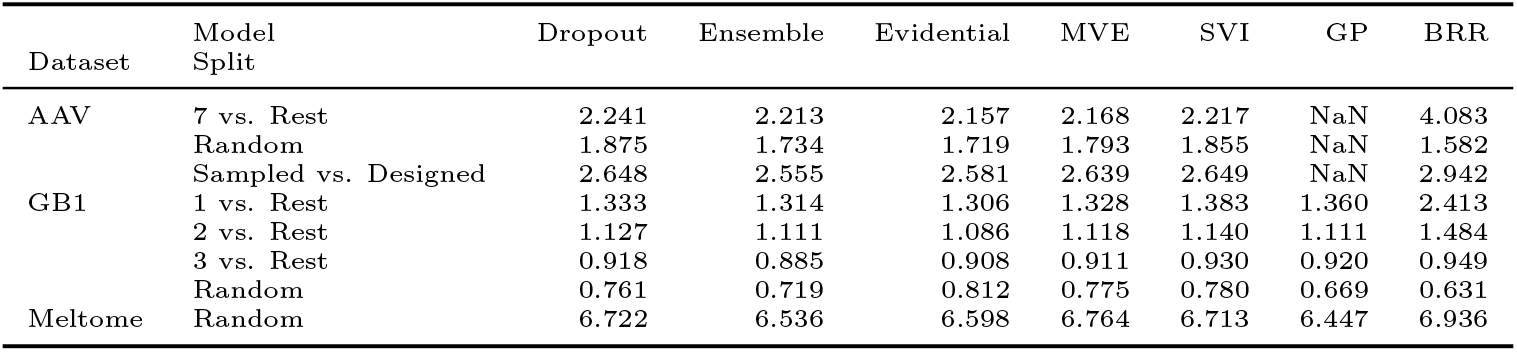
Test set RMSE for models trained on ESM representation (↓)

**Table S13:**
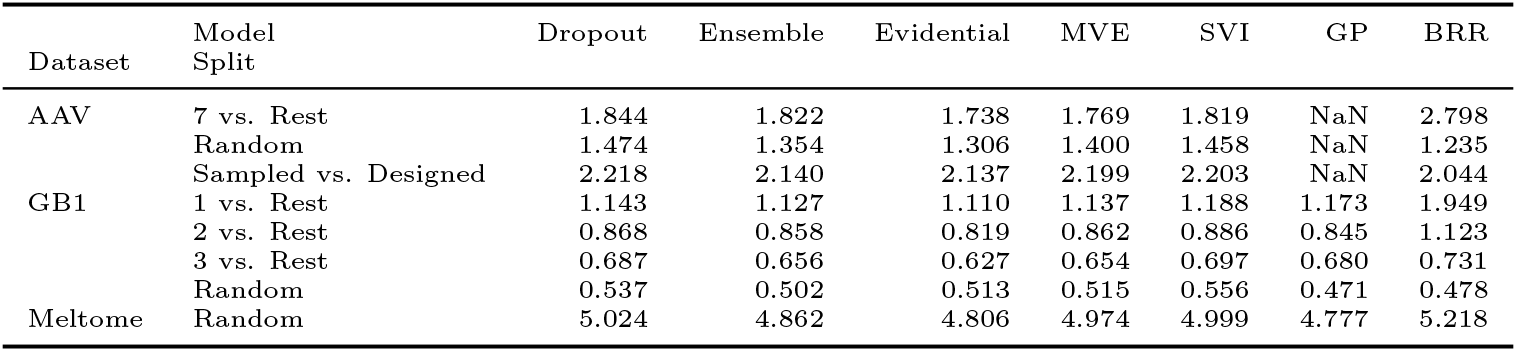
Test set MAE for models trained on ESM representation (↓)

**Table S14:**
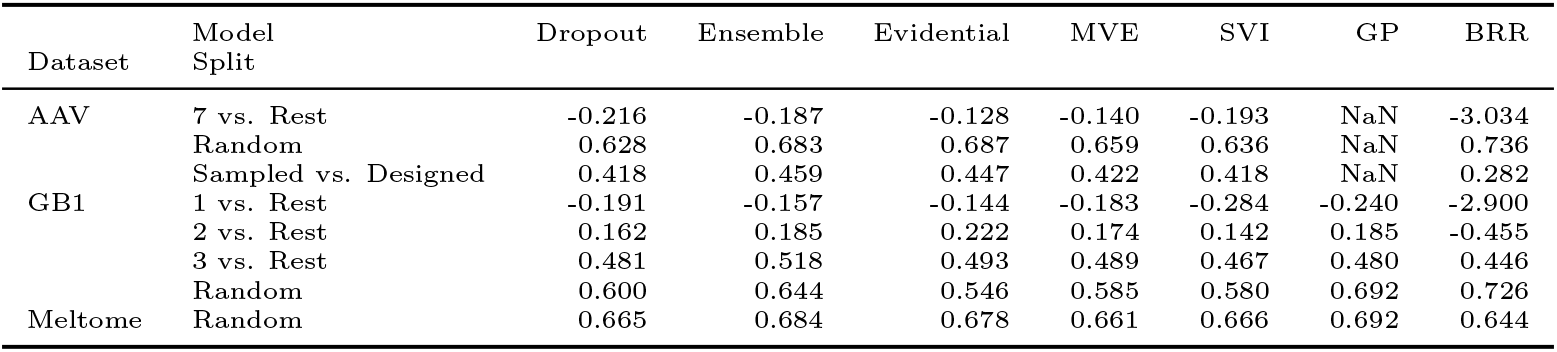
Test set 𝑅^2^ for models trained on ESM representation (↑)

**Table S15:**
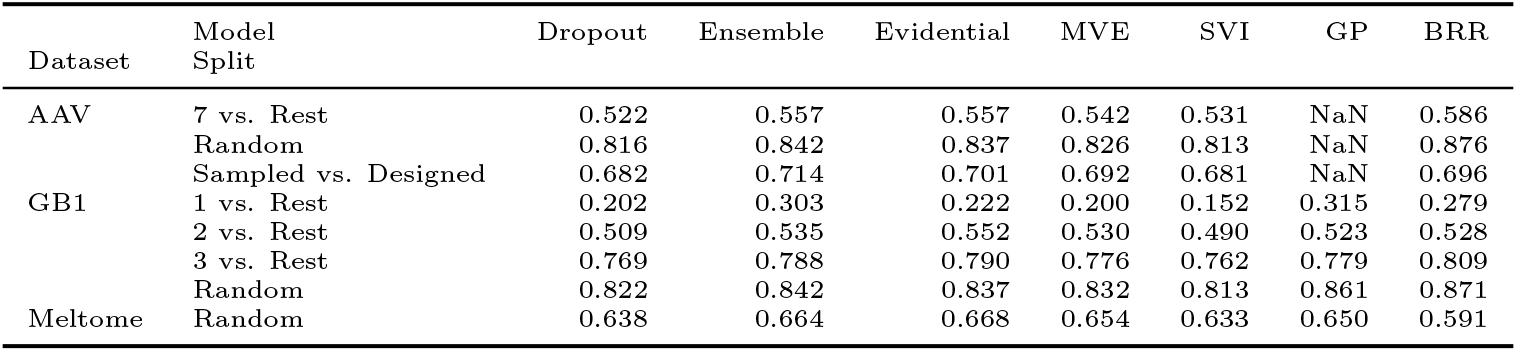
Test set 𝜌 for models trained on ESM representation (↑)

**Table S16:**
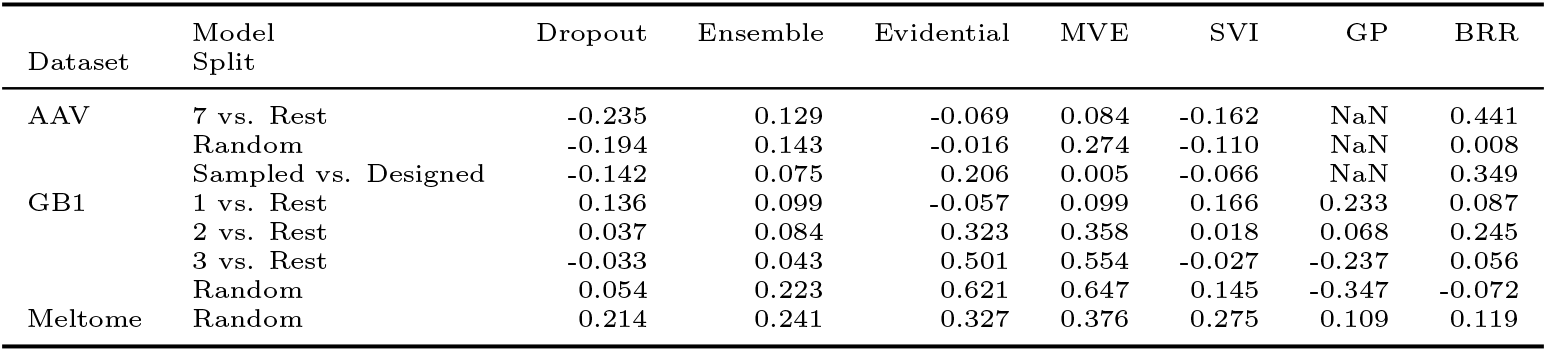
Test set 𝜌_unc_ for models trained on ESM representation (↑)

**Table S17:**
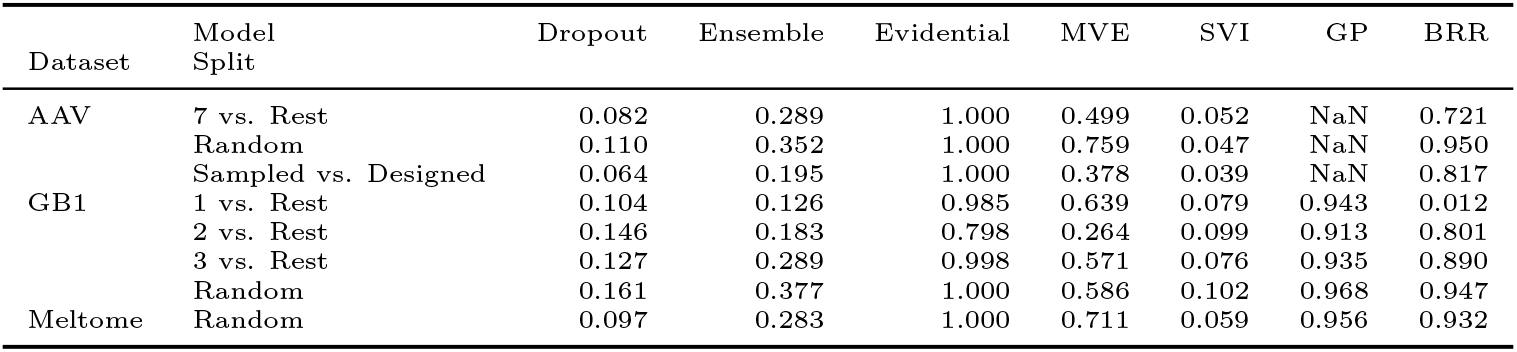
Test set % coverage for models trained on ESM representation (↑)

**Table S18:**
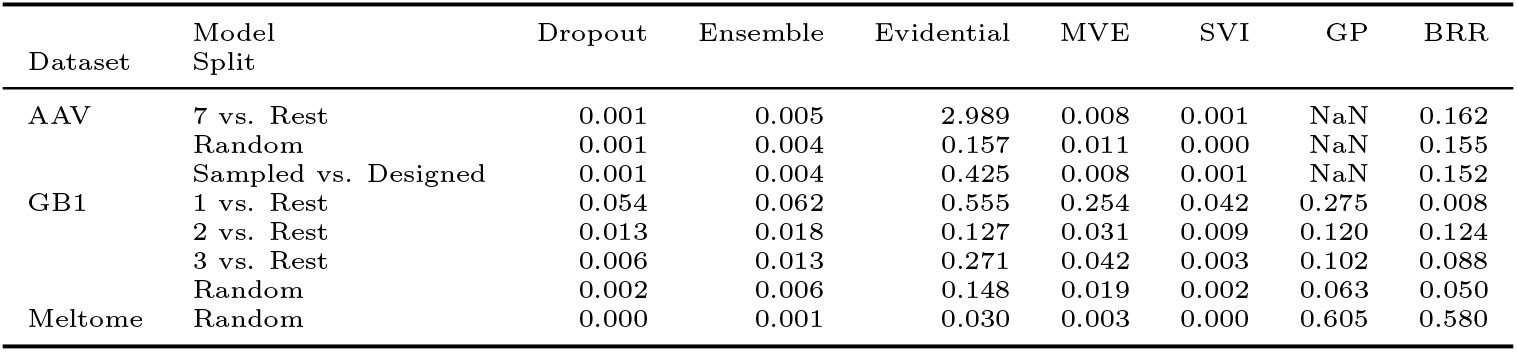
Test set 4𝜎/𝑅 for models trained on ESM representation (↓)

**Table S19:**
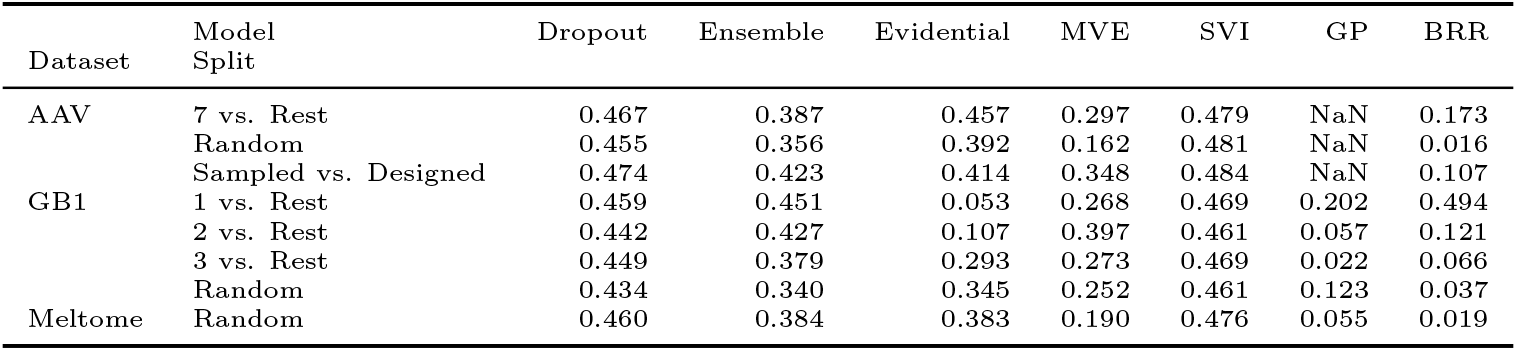
Test set miscalibration area for models trained on ESM representation (↓)

**Table S20:**
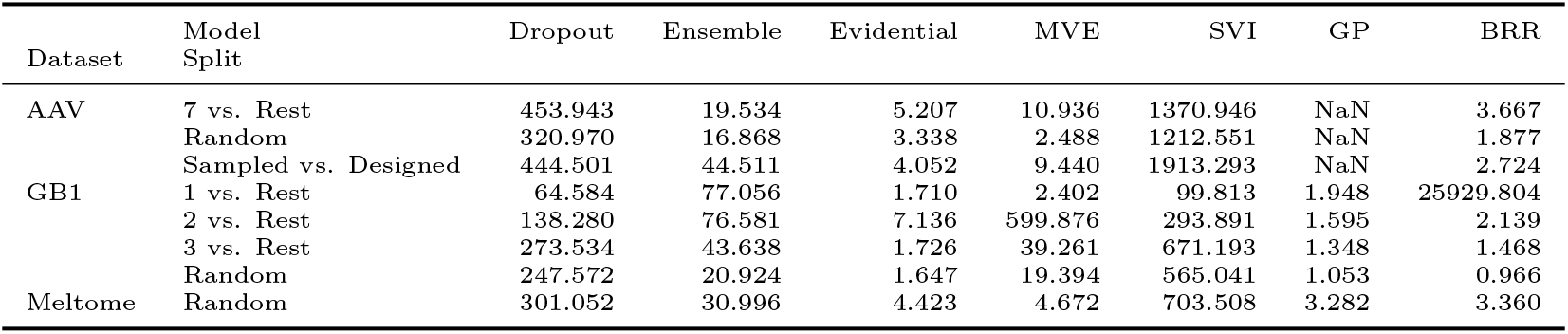
Test set 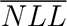 for models trained on ESM representation (↓)

**Table S21:**
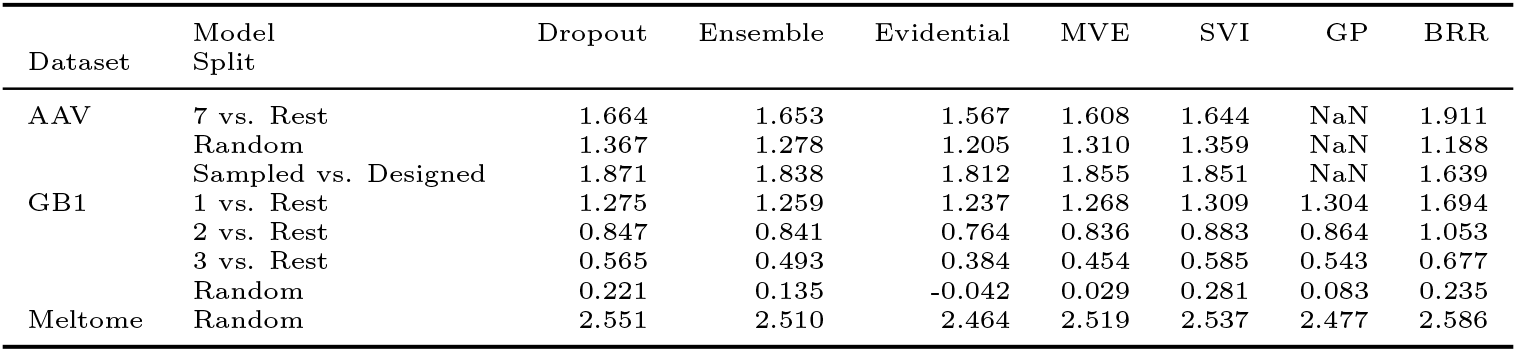
Test set 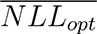 for models trained on ESM representation

**Table S22:** Test set 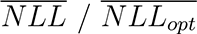 ratio for models trained on ESM representation (↓)

| Dataset | Model Split | Dropout | Ensemble | Evidential | MVE | SVI | GP | BRR |
| --- | --- | --- | --- | --- | --- | --- | --- | --- |
| AAV | 7 vs. Rest | 273.638 | 11.806 | 3.324 | 6.734 | 826.919 | NaN | 1.919 |
|  | Random | 228.987 | 13.229 | 2.786 | 1.915 | 868.280 | NaN | 1.579 |
|  | Sampled vs. Designed | 235.180 | 24.210 | 2.239 | 5.094 | 1013.232 | NaN | 1.662 |
| GB1 | 1 vs. Rest | 50.427 | 61.282 | 1.385 | 1.901 | 77.453 | 1.494 | 15310.669 |
|  | 2 vs. Rest | 160.595 | 91.310 | 8.750 | 697.394 | 328.277 | 1.846 | 2.031 |
|  | 3 vs. Rest | 453.390 | 89.292 | 4.715 | 90.982 | 1070.519 | 2.483 | 2.168 |
|  | Random | 991.498 | 168.540 | -6.076 | 42.706 | 1751.474 | 12.645 | 4.110 |
| Meltome | Random | 117.634 | 12.360 | 1.796 | 1.856 | 276.206 | 1.325 | 1.299 |

### 5 Active Learning

**Figure S5:**
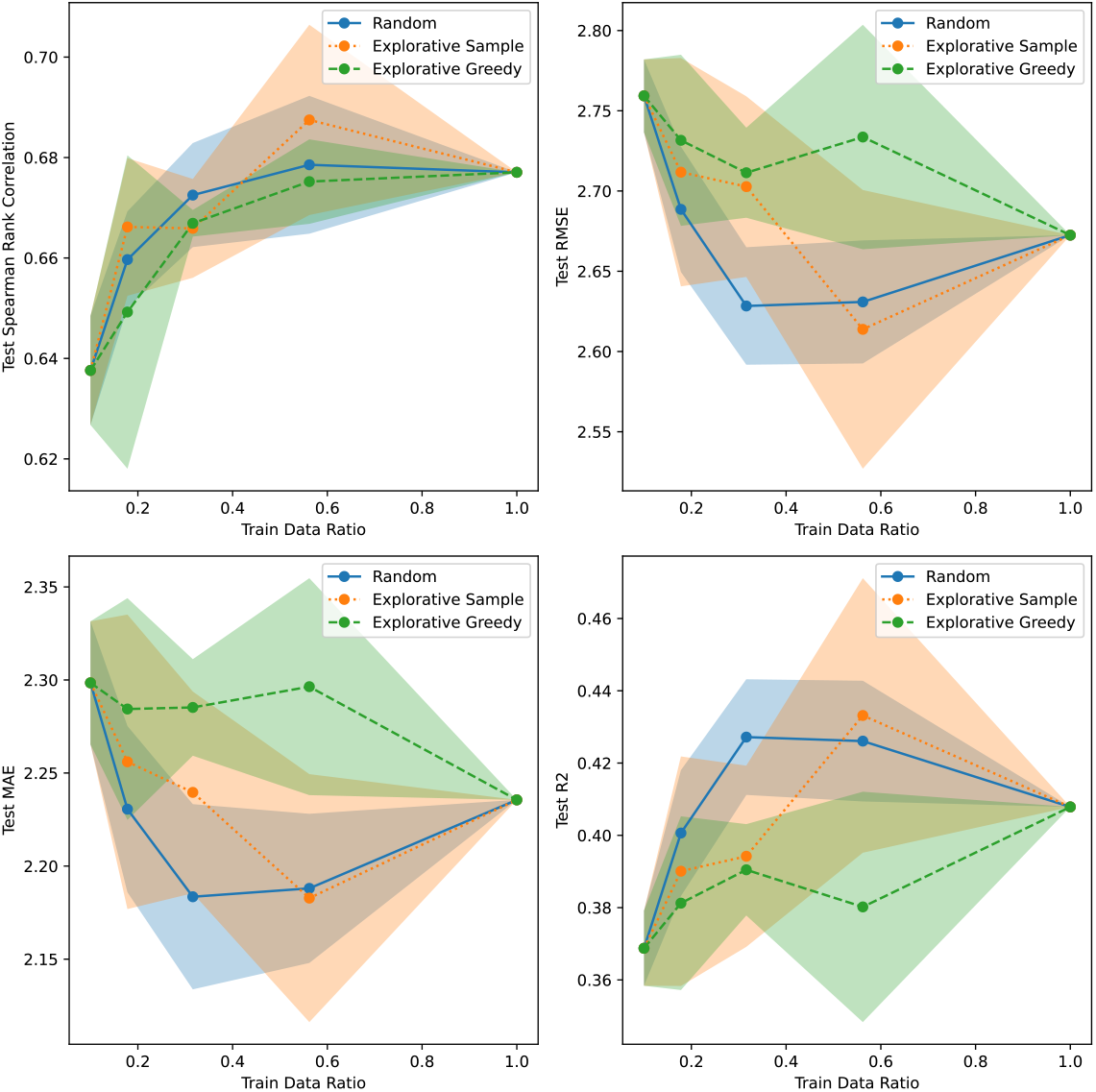
Active learning results for AAV/Sampled vs. Designed using CNN Dropout uncertainty.

**Figure S6:**
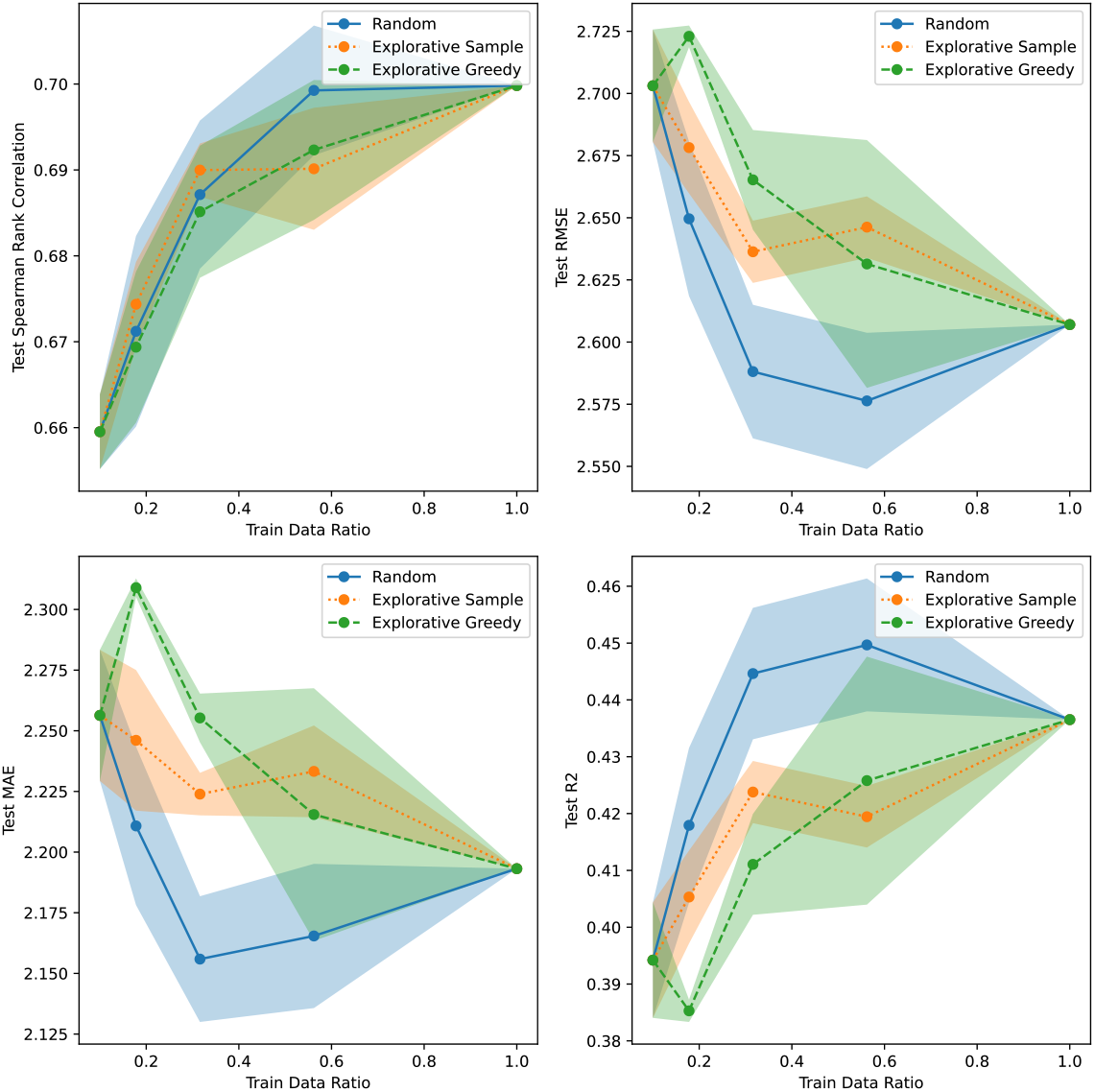
Active learning results for AAV/Sampled vs. Designed using CNN Ensemble uncertainty.

**Figure S7:**
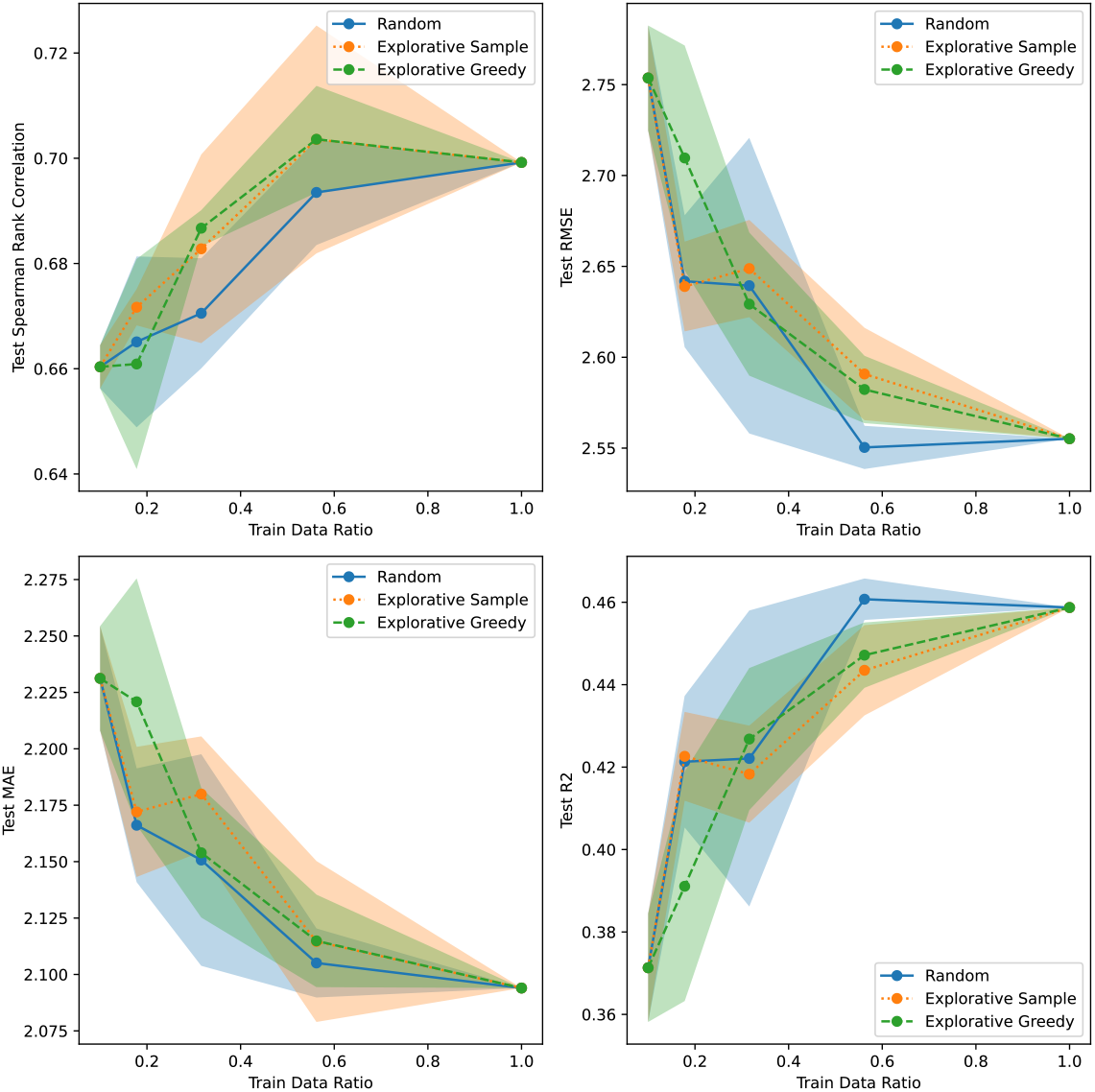
Active learning results for AAV/Sampled vs. Designed using CNN Evidential uncertainty.

**Figure S8:**
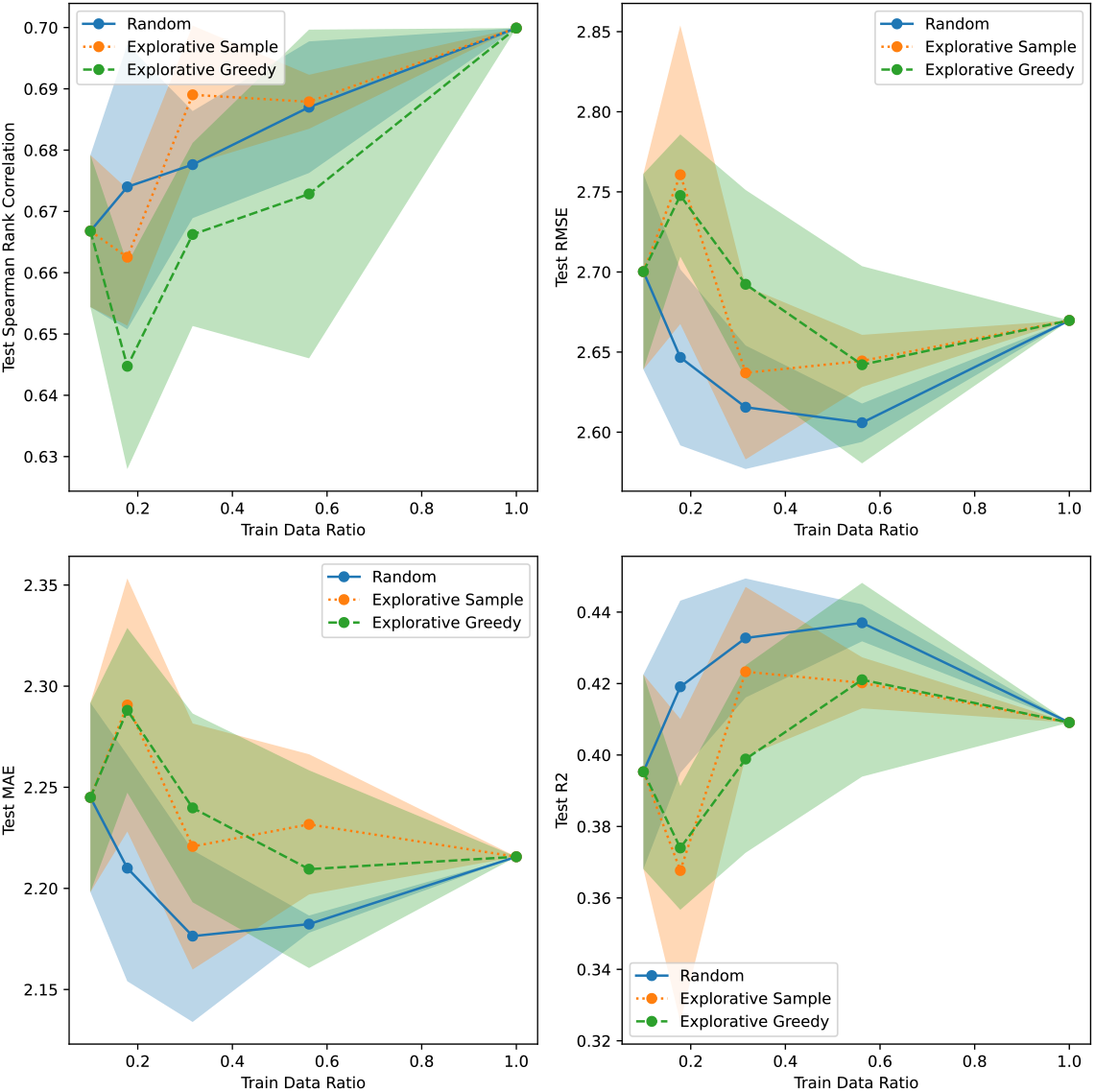
Active learning results for AAV/Sampled vs. Designed using CNN MVE uncertainty.

**Figure S9:**
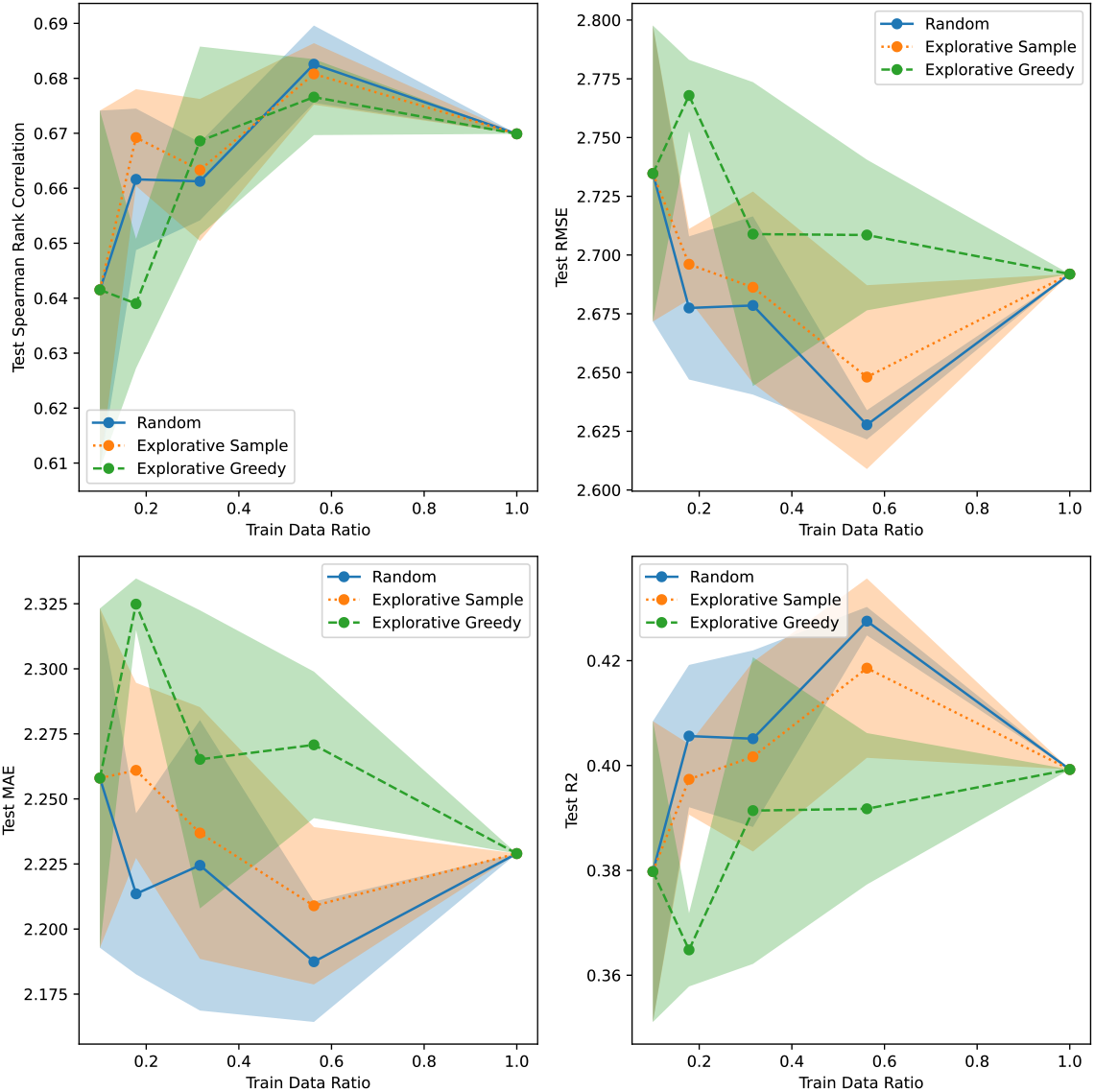
Active learning results for AAV/Sampled vs. Designed using CNN SVI uncertainty.

**Figure S10:**
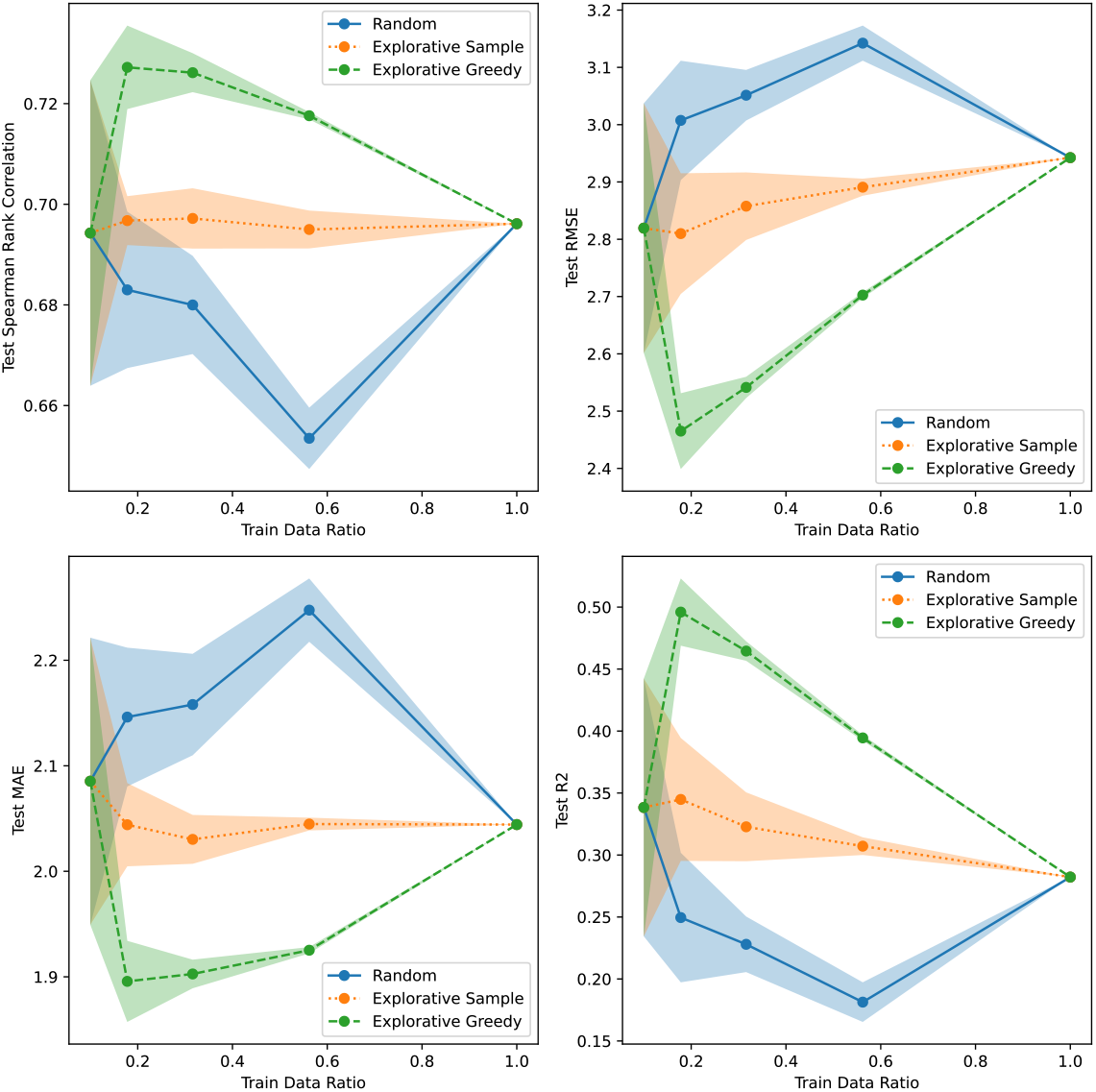
Active learning results for AAV/Sampled vs. Designed using Linear Bayesian Ridge uncertainty.

**Figure S11:**
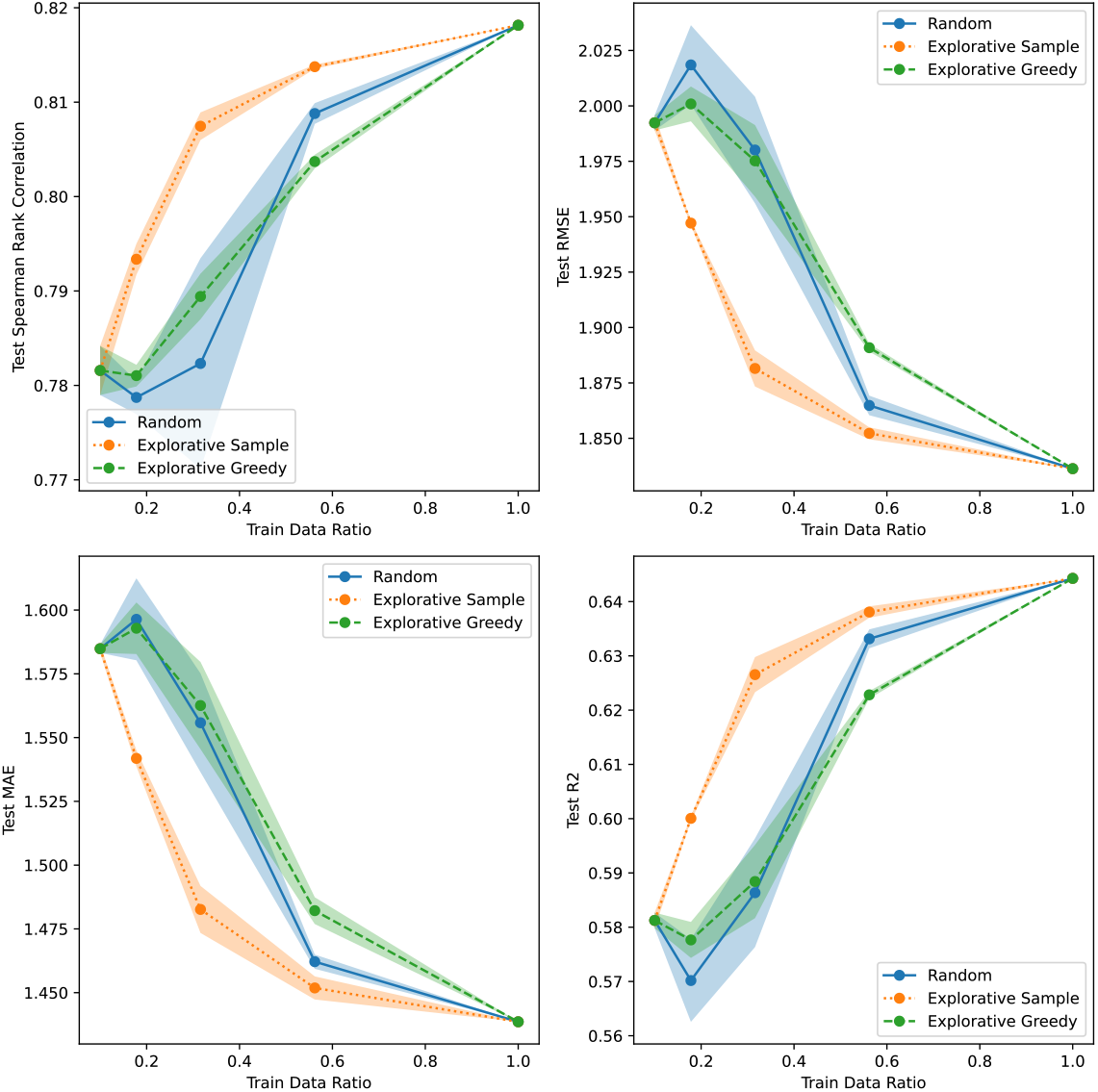
Active learning results for AAV/Random using CNN Dropout uncertainty.

**Figure S12:**
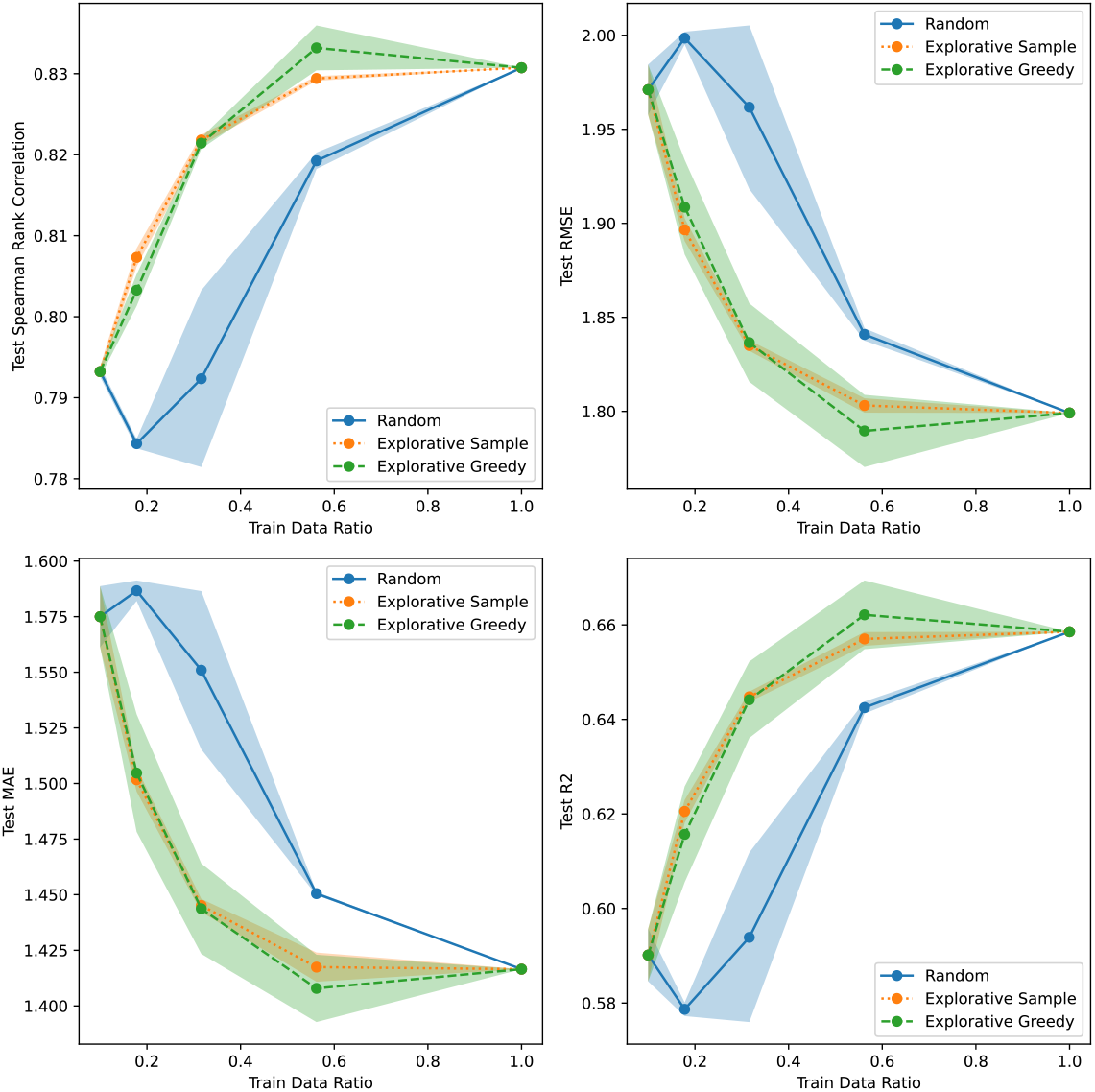
Active learning results for AAV/Random using CNN Ensemble uncertainty.

**Figure S13:**
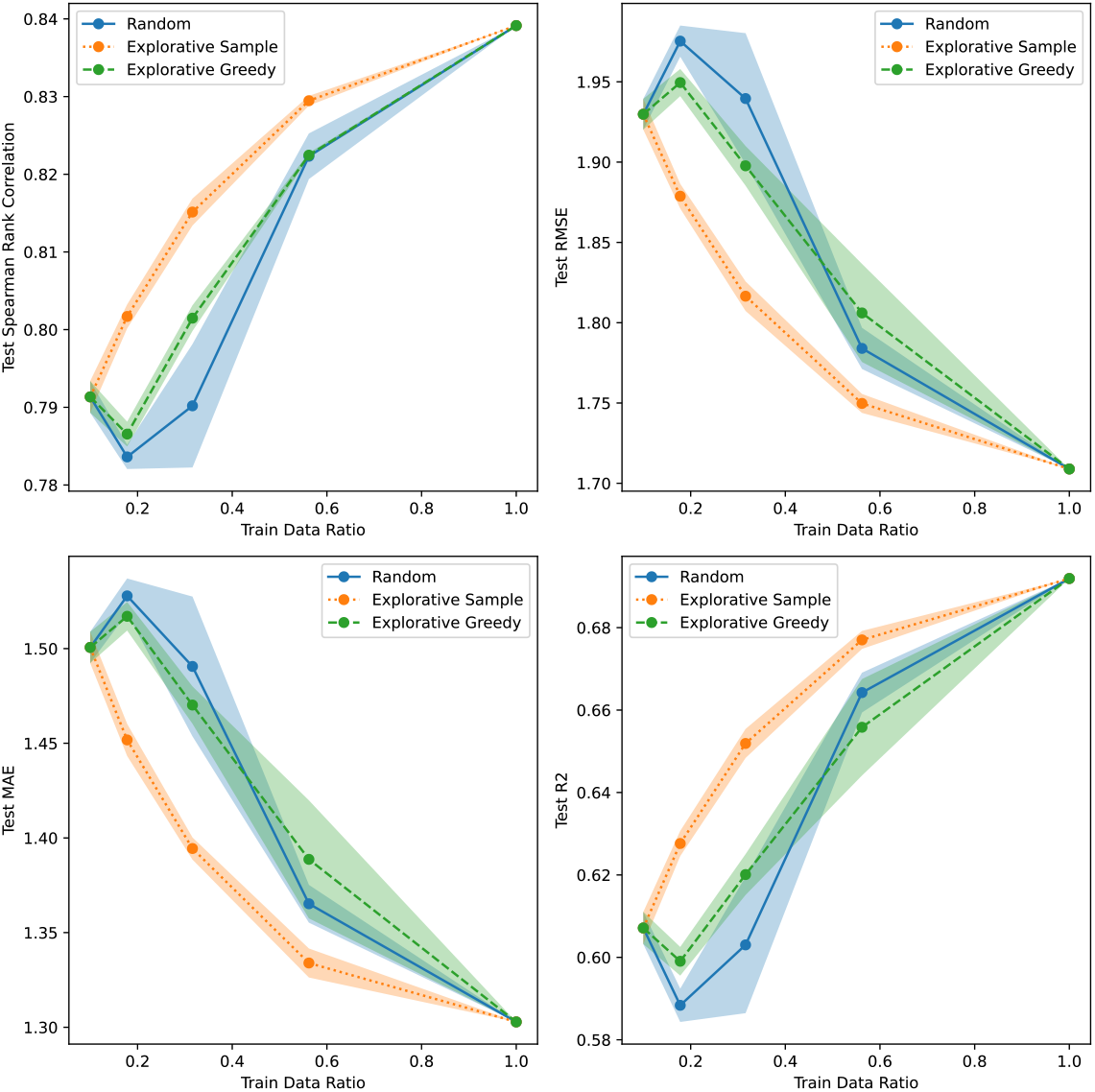
Active learning results for AAV/Random using CNN Evidential uncertainty.

**Figure S14:**
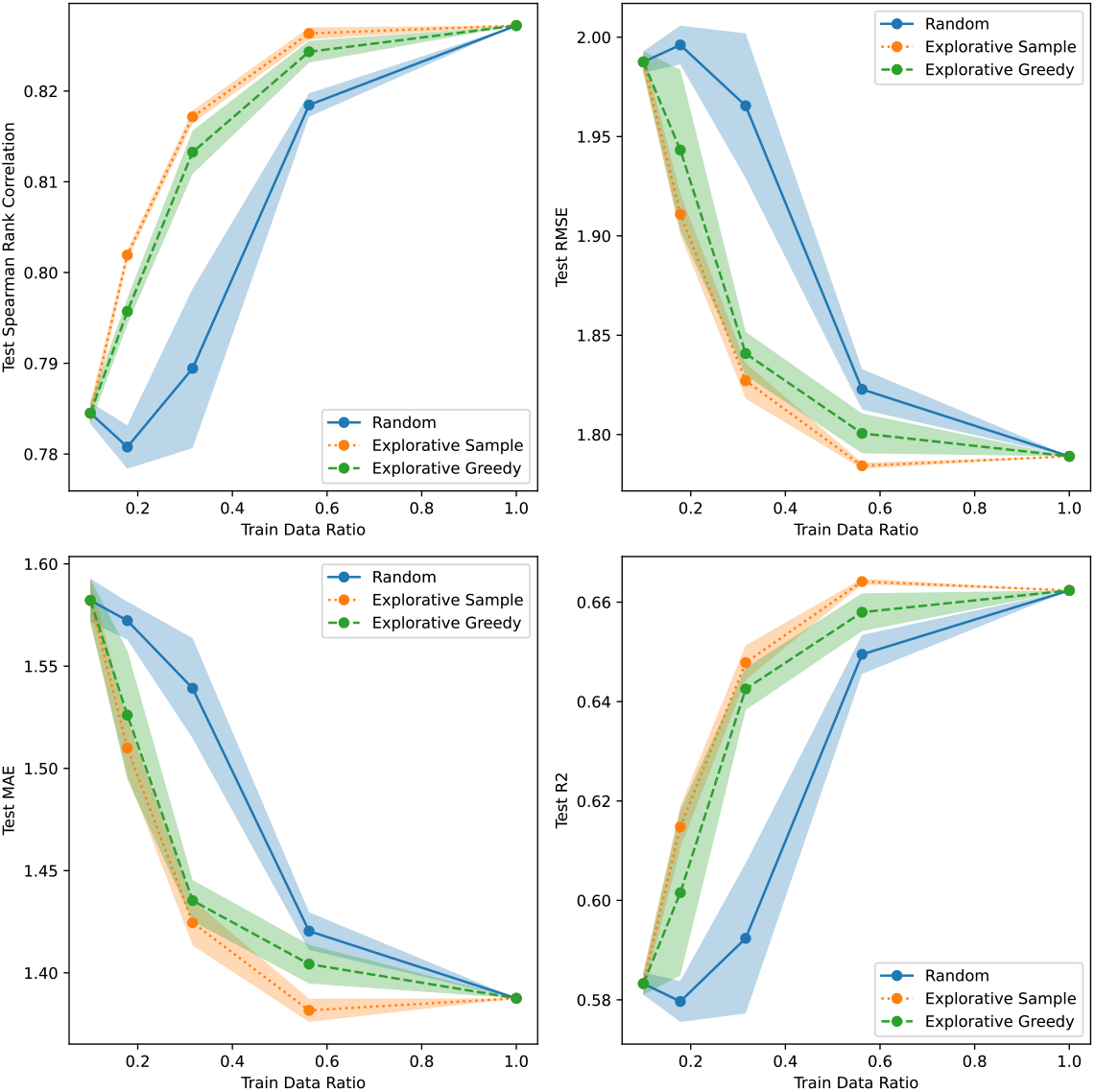
Active learning results for AAV/Random using CNN MVE uncertainty.

**Figure S15:**
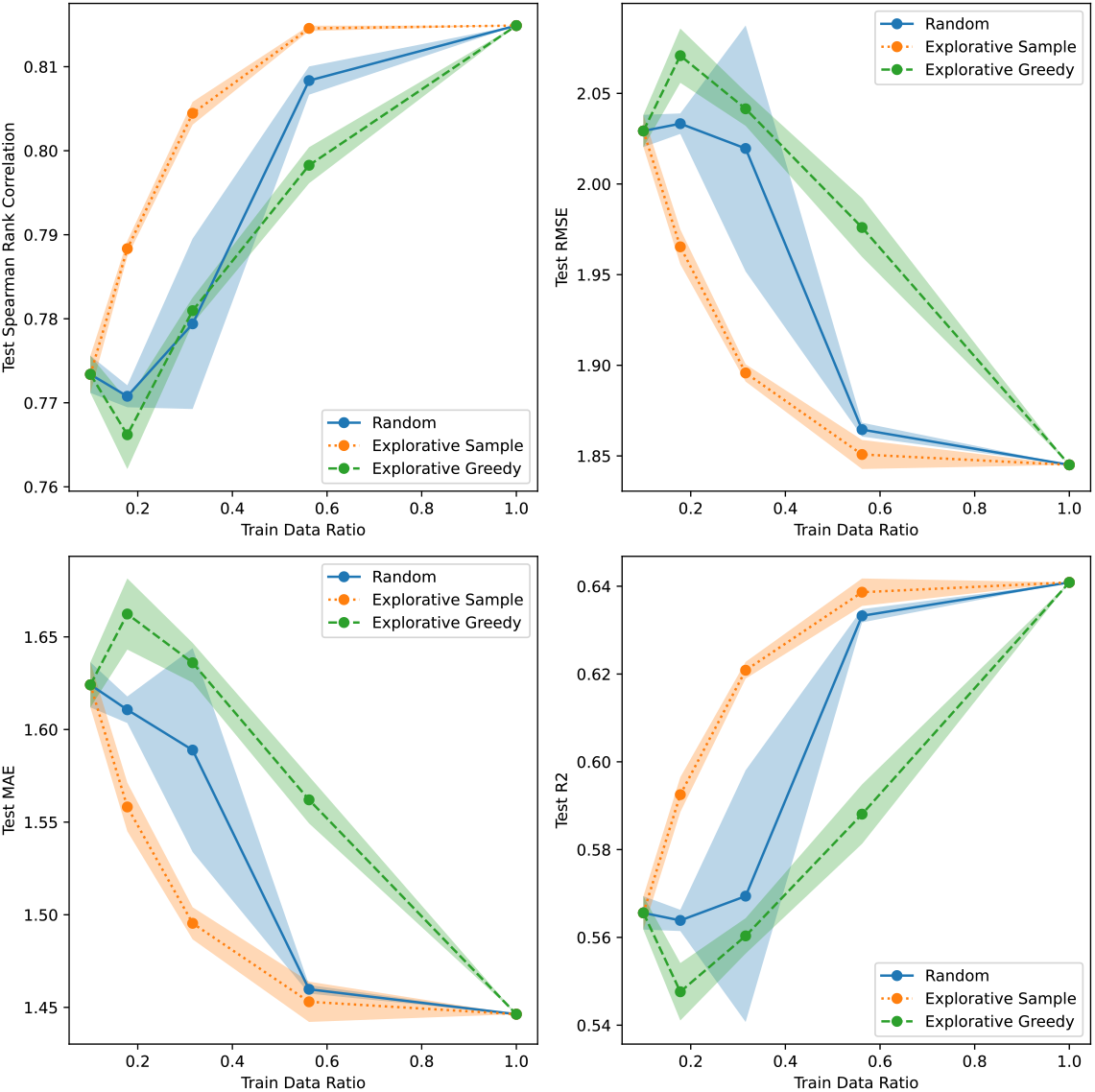
Active learning results for AAV/Random using CNN SVI uncertainty.

**Figure S16:**
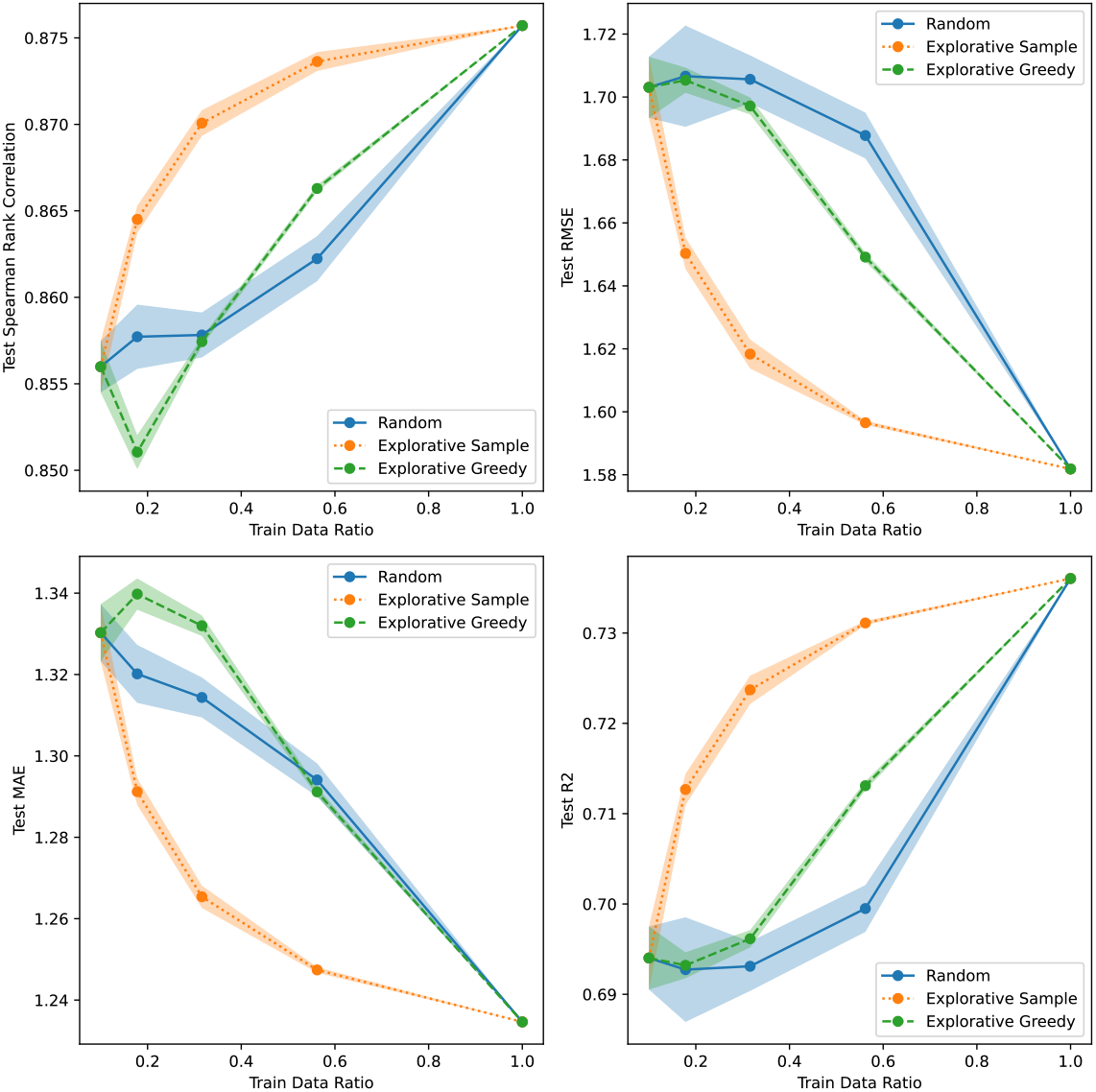
Active learning results for AAV/Random using Linear Bayesian Ridge uncertainty.

**Figure S17:**
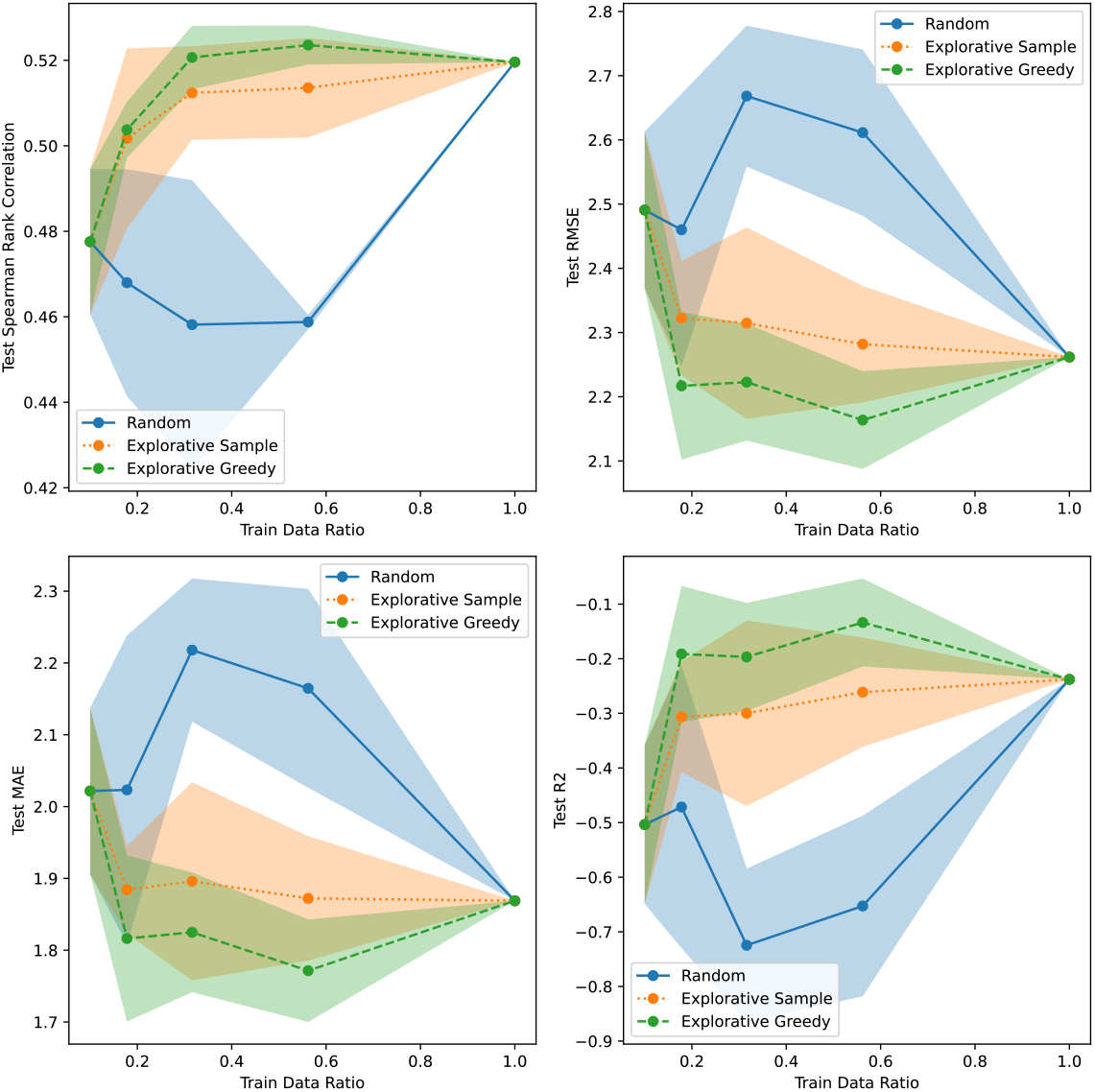
Active learning results for AAV/7 vs. Rest using CNN Dropout uncertainty.

**Figure S18:**
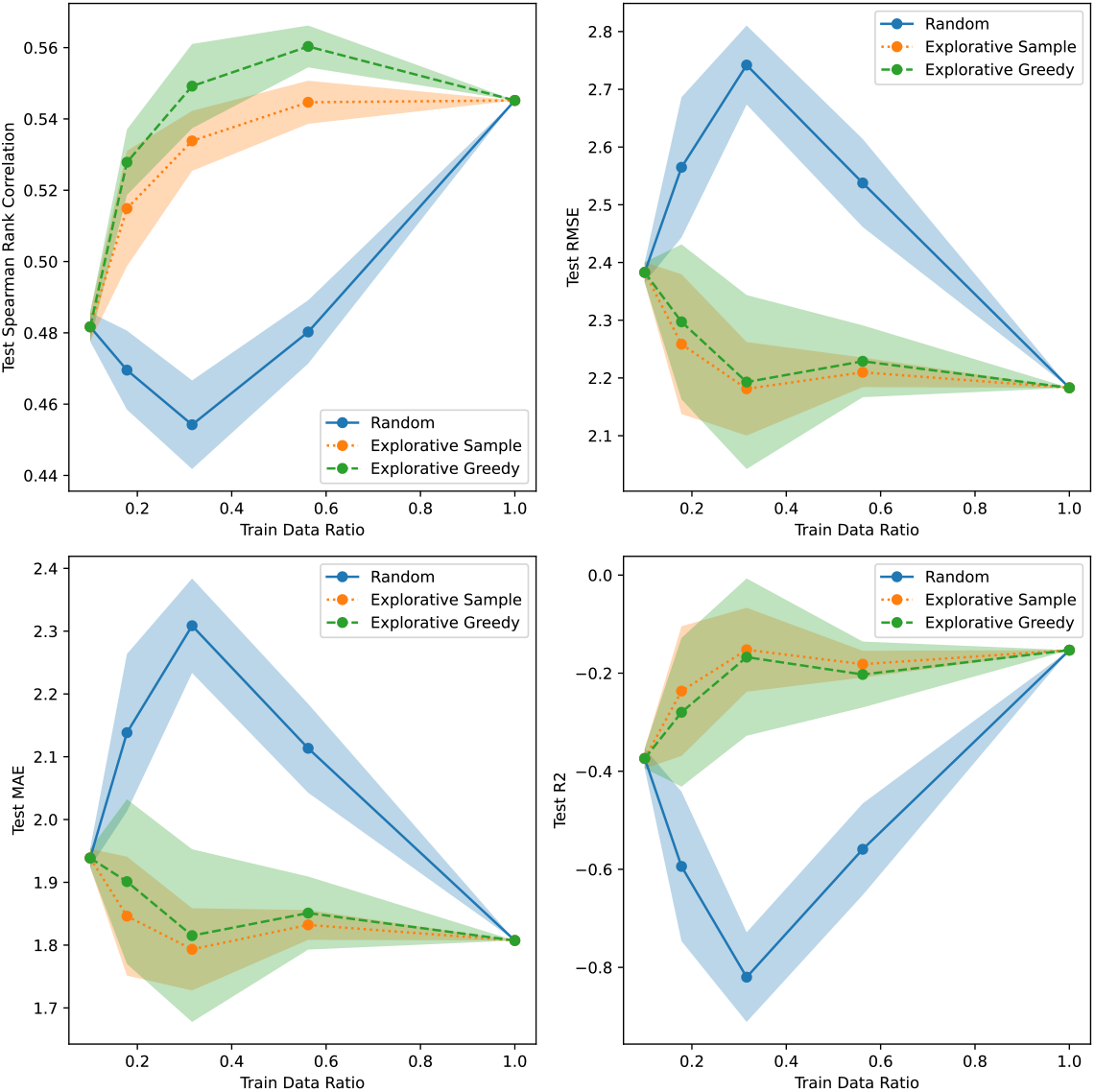
Active learning results for AAV/7 vs. Rest using CNN Ensemble uncertainty.

**Figure S19:**
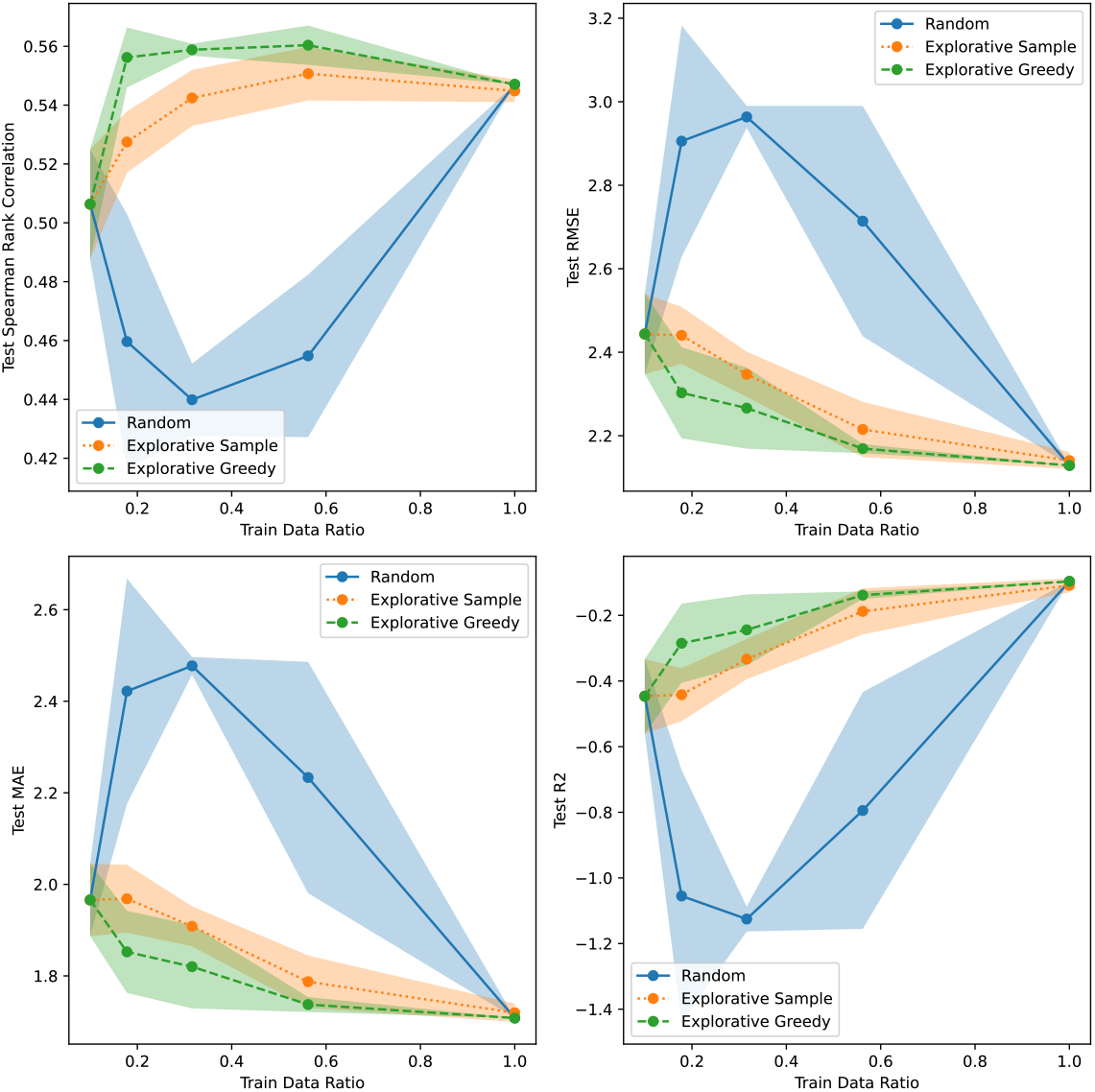
Active learning results for AAV/7 vs. Rest using CNN Evidential uncertainty.

**Figure S20:**
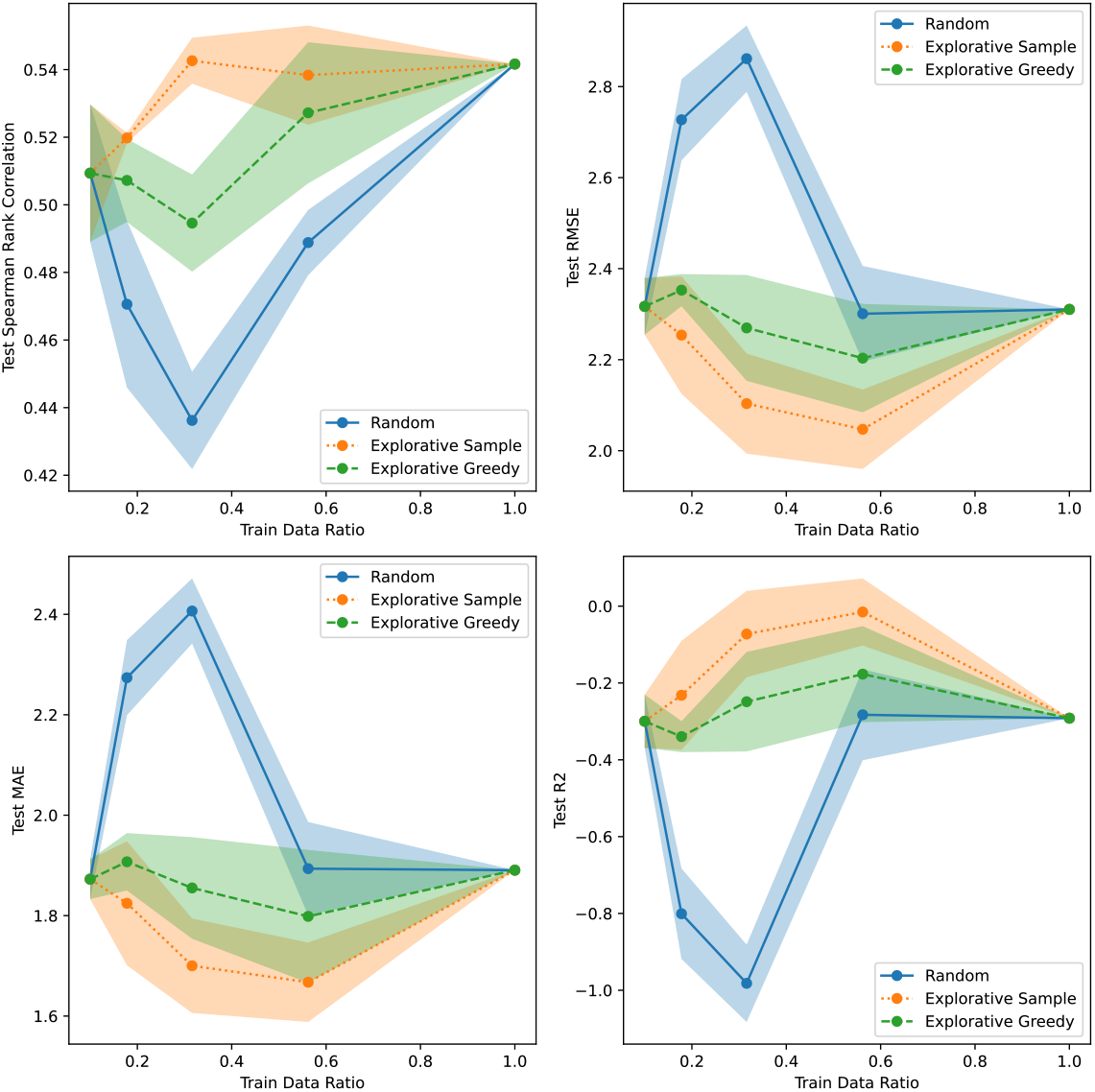
Active learning results for AAV/7 vs. Rest using CNN MVE uncertainty.

**Figure S21:**
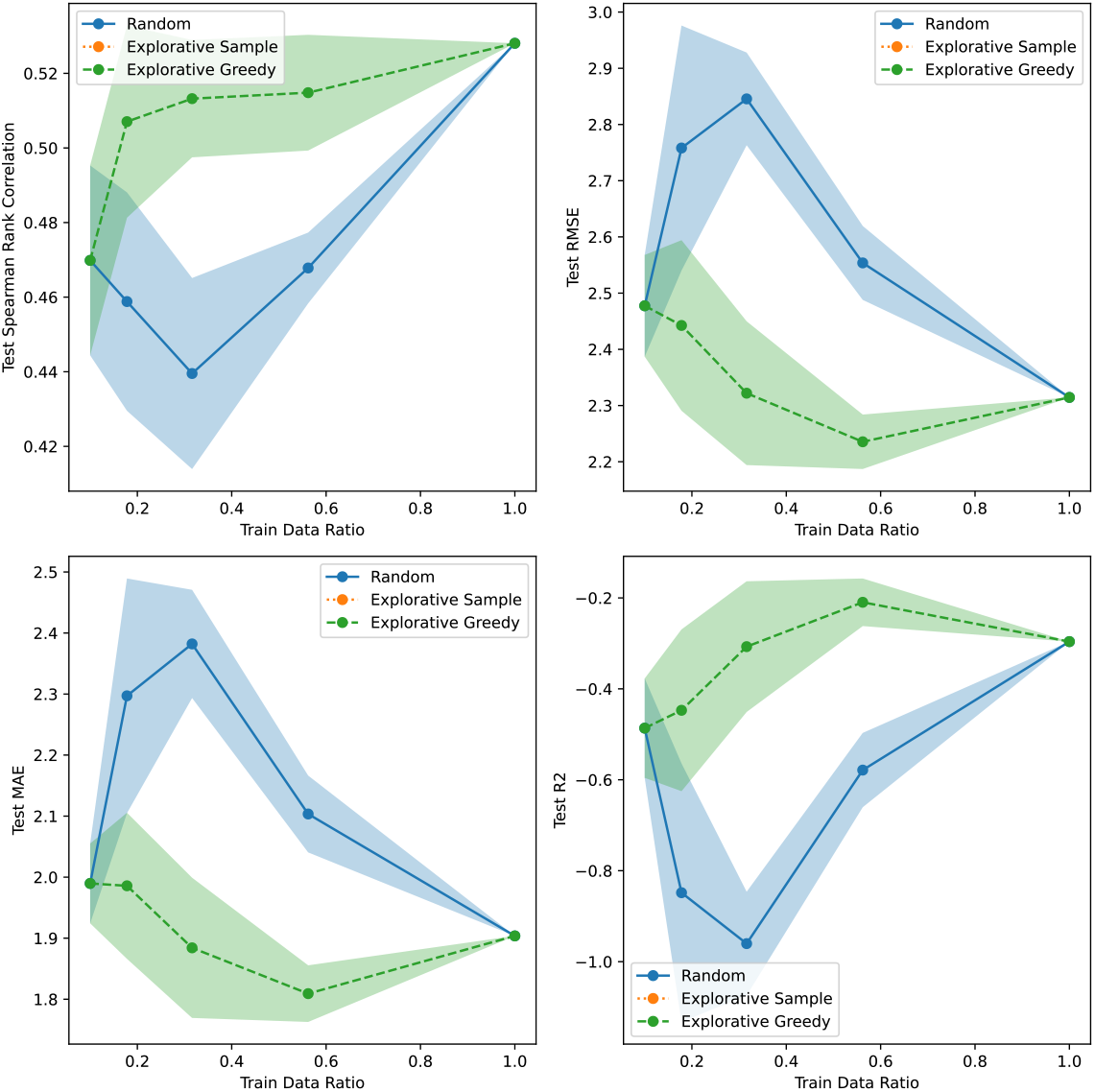
Active learning results for AAV/7 vs. Rest using CNN SVI uncertainty.

**Figure S22:**
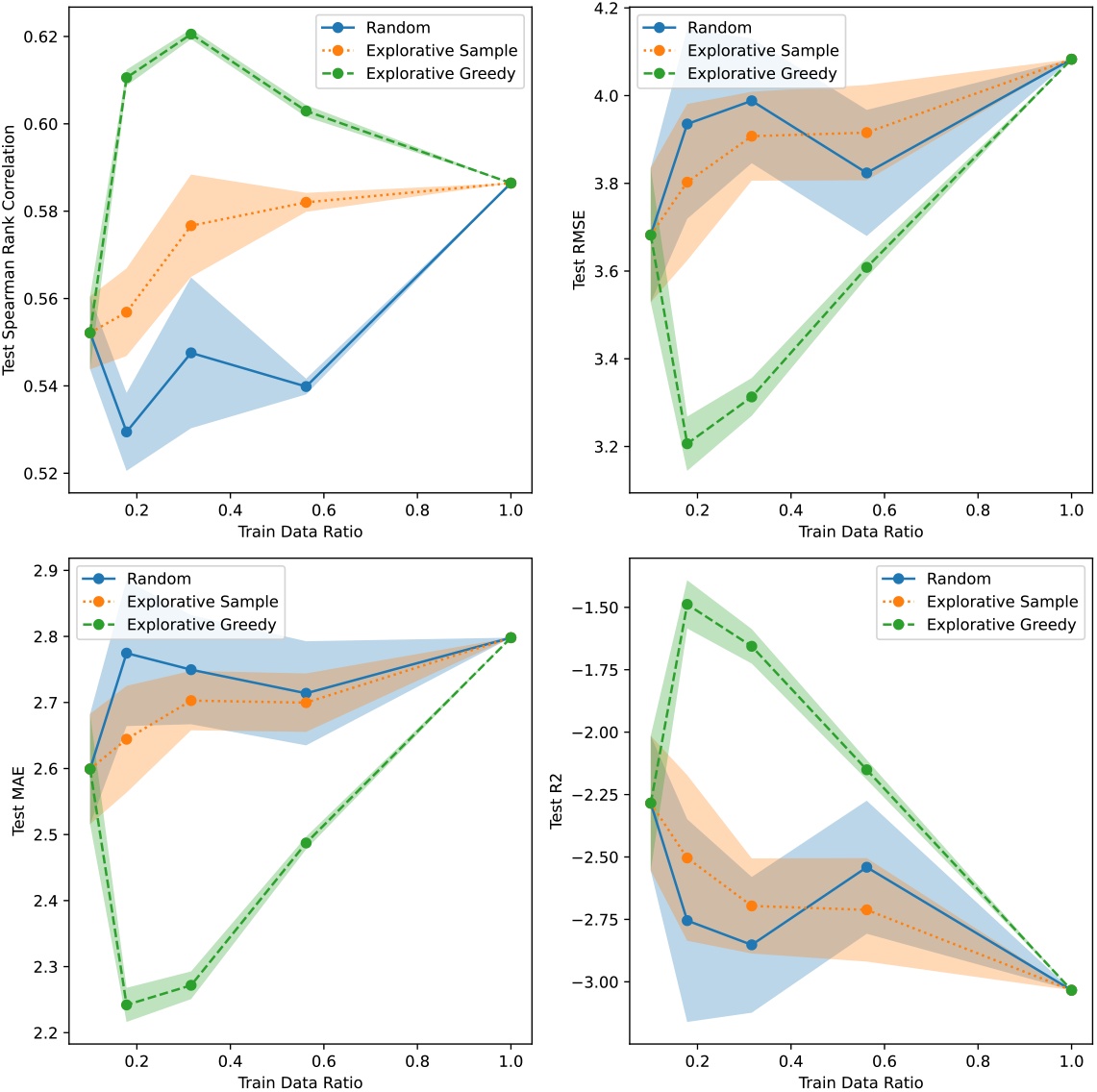
Active learning results for AAV/7 vs. Rest using Linear Bayesian Ridge uncertainty.

**Figure S23:**
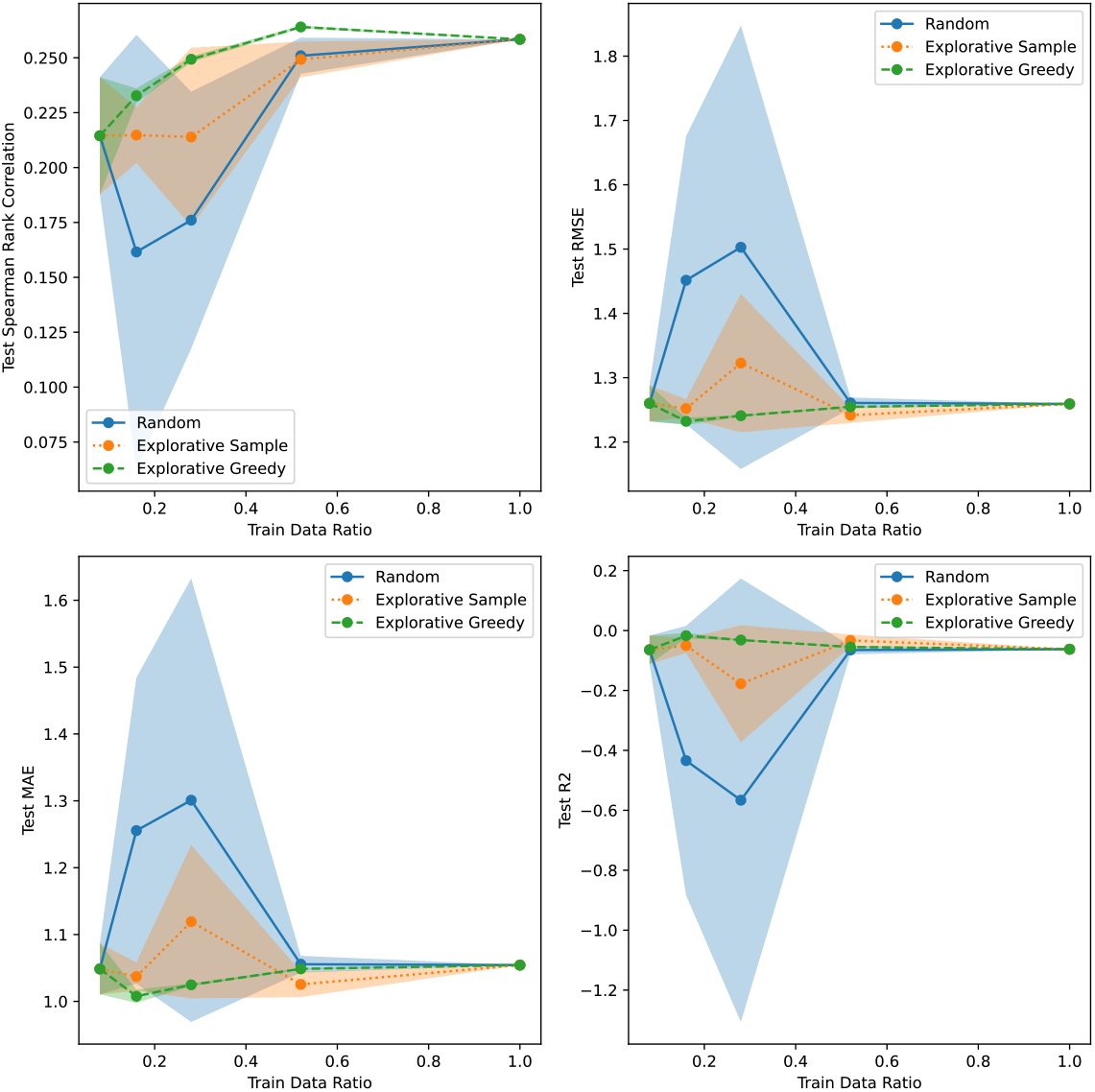
Active learning results for GB1/1 vs. Rest using CNN Dropout uncertainty.

**Figure S24:**
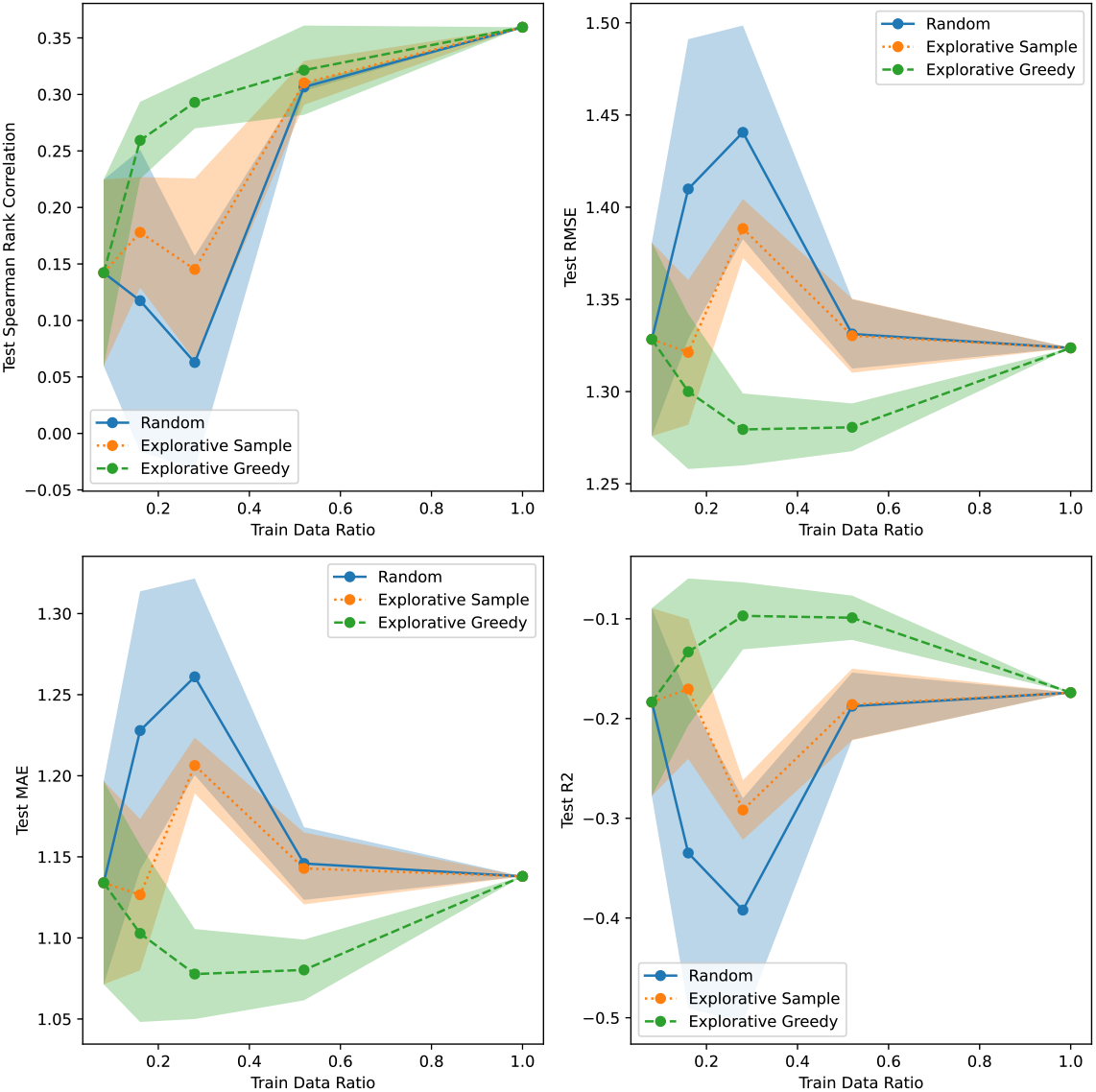
Active learning results for GB1/1 vs. Rest using CNN Ensemble uncertainty.

**Figure S25:**
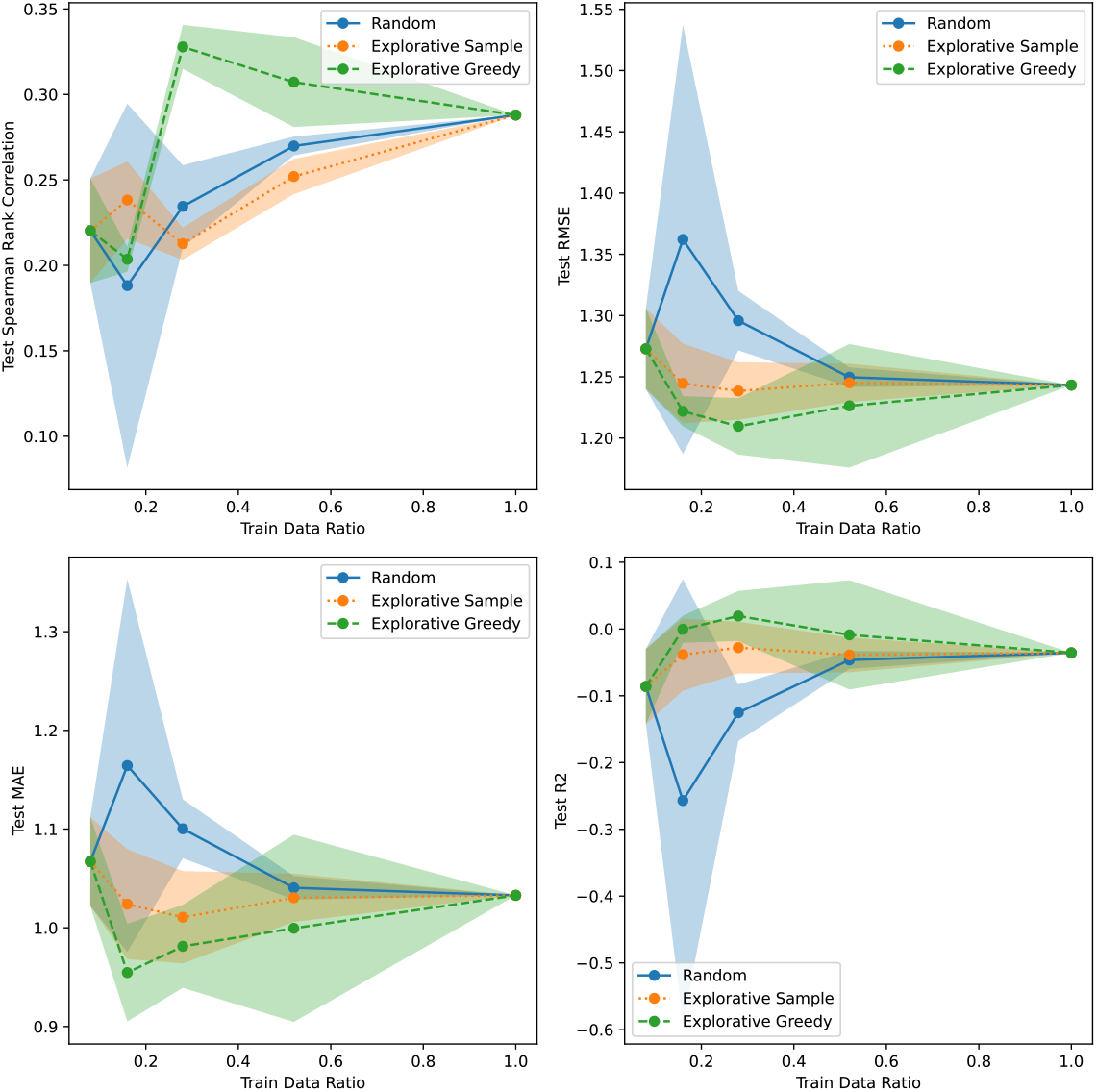
Active learning results for GB1/1 vs. Rest using CNN Evidential uncertainty.

**Figure S26:**
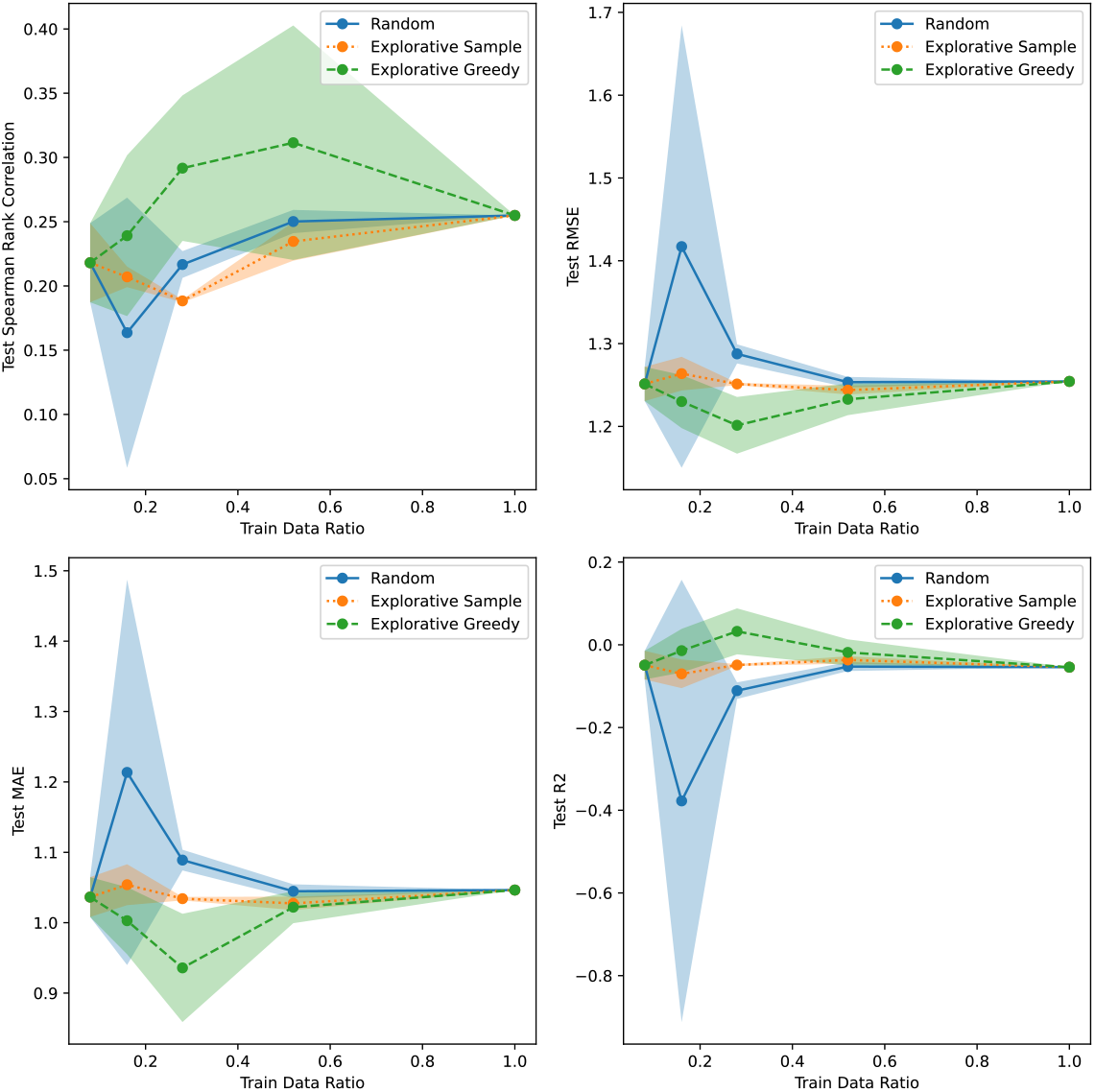
Active learning results for GB1/1 vs. Rest using CNN MVE uncertainty.

**Figure S27:**
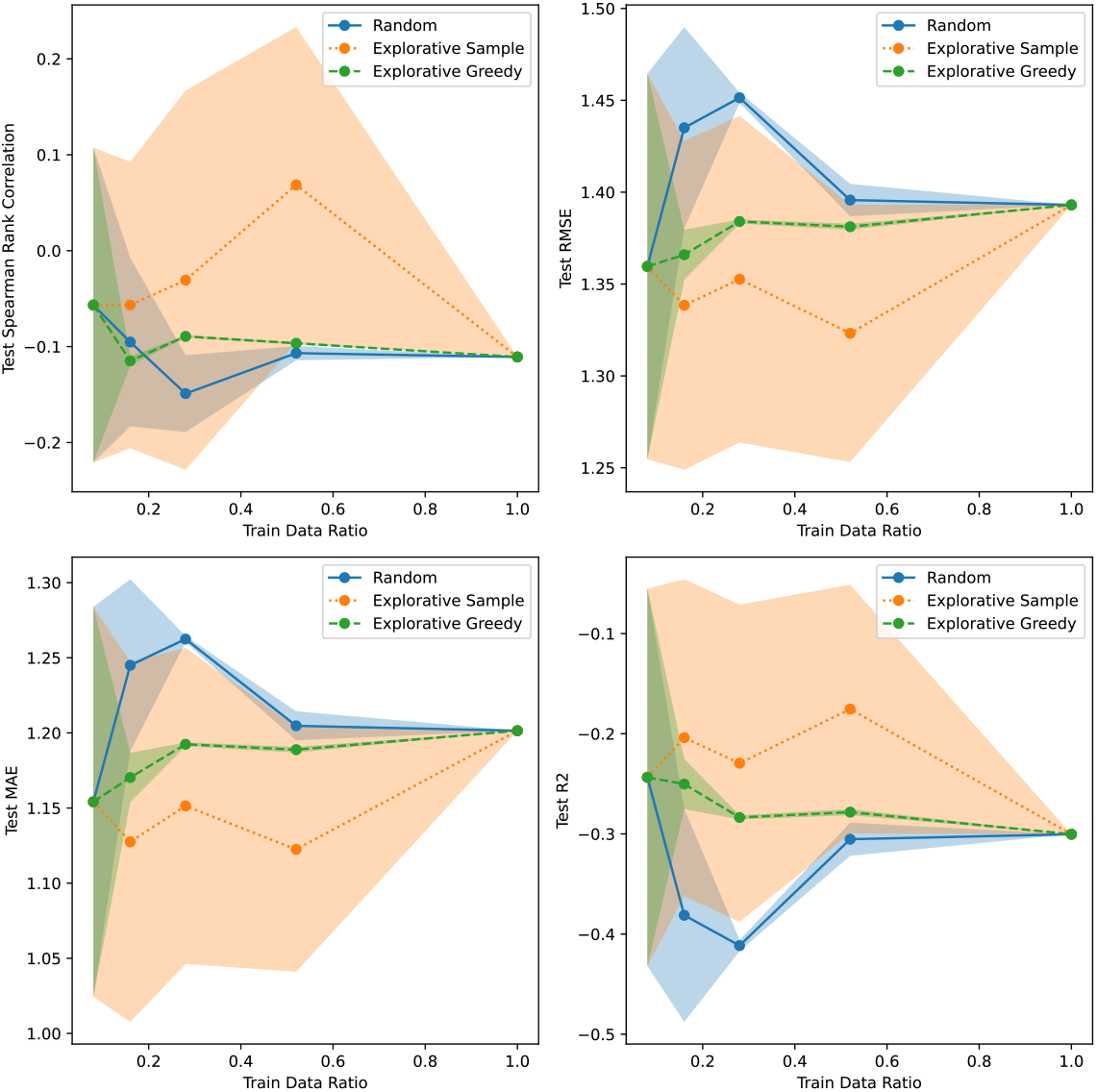
Active learning results for GB1/1 vs. Rest using CNN SVI uncertainty.

**Figure S28:**
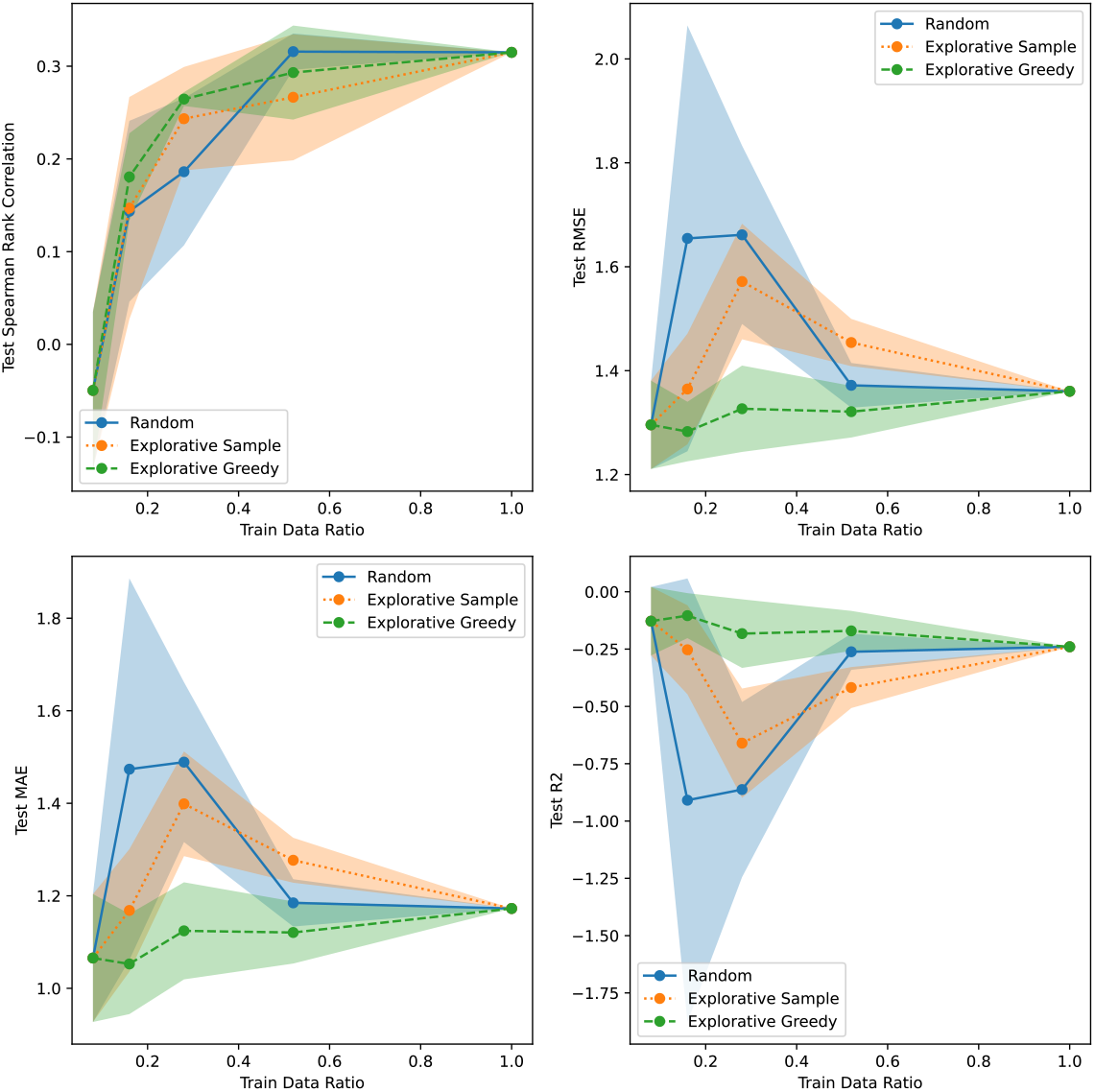
Active learning results for GB1/1 vs. Rest using GP Continuous uncertainty.

**Figure S29:**
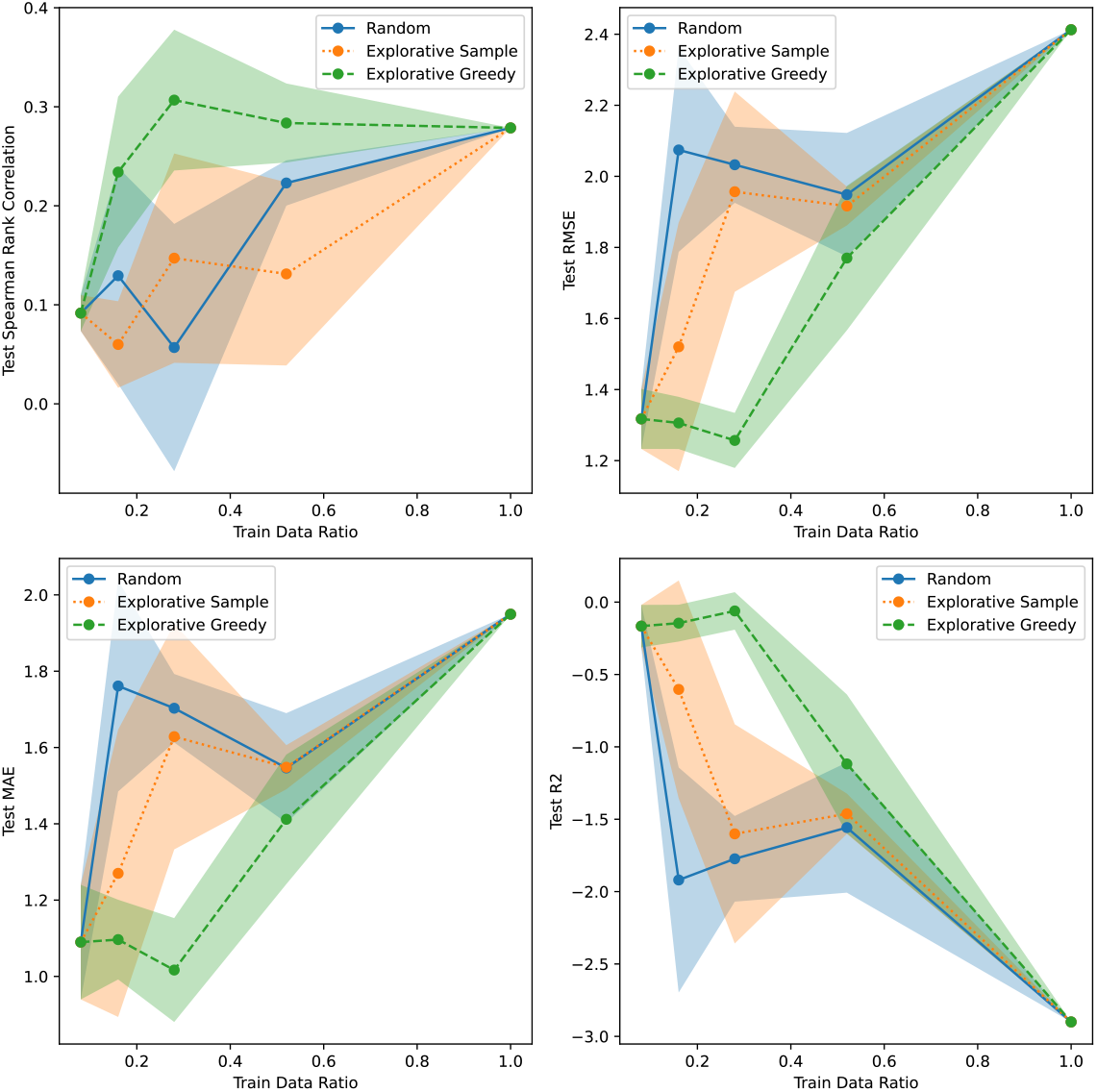
Active learning results for GB1/1 vs. Rest using Linear Bayesian Ridge uncertainty.

**Figure S30:**
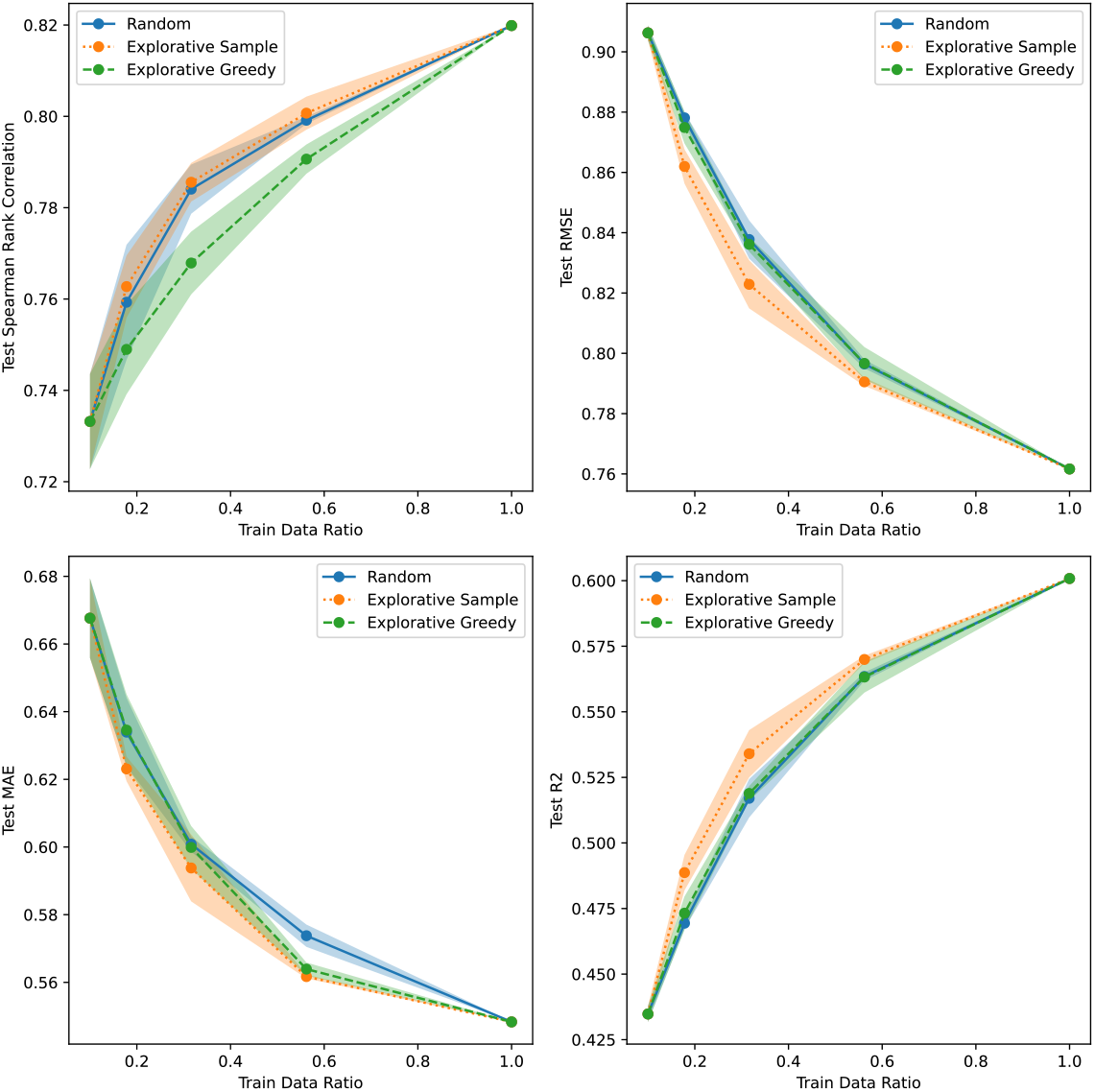
Active learning results for GB1/Random using CNN Dropout uncertainty.

**Figure S31:**
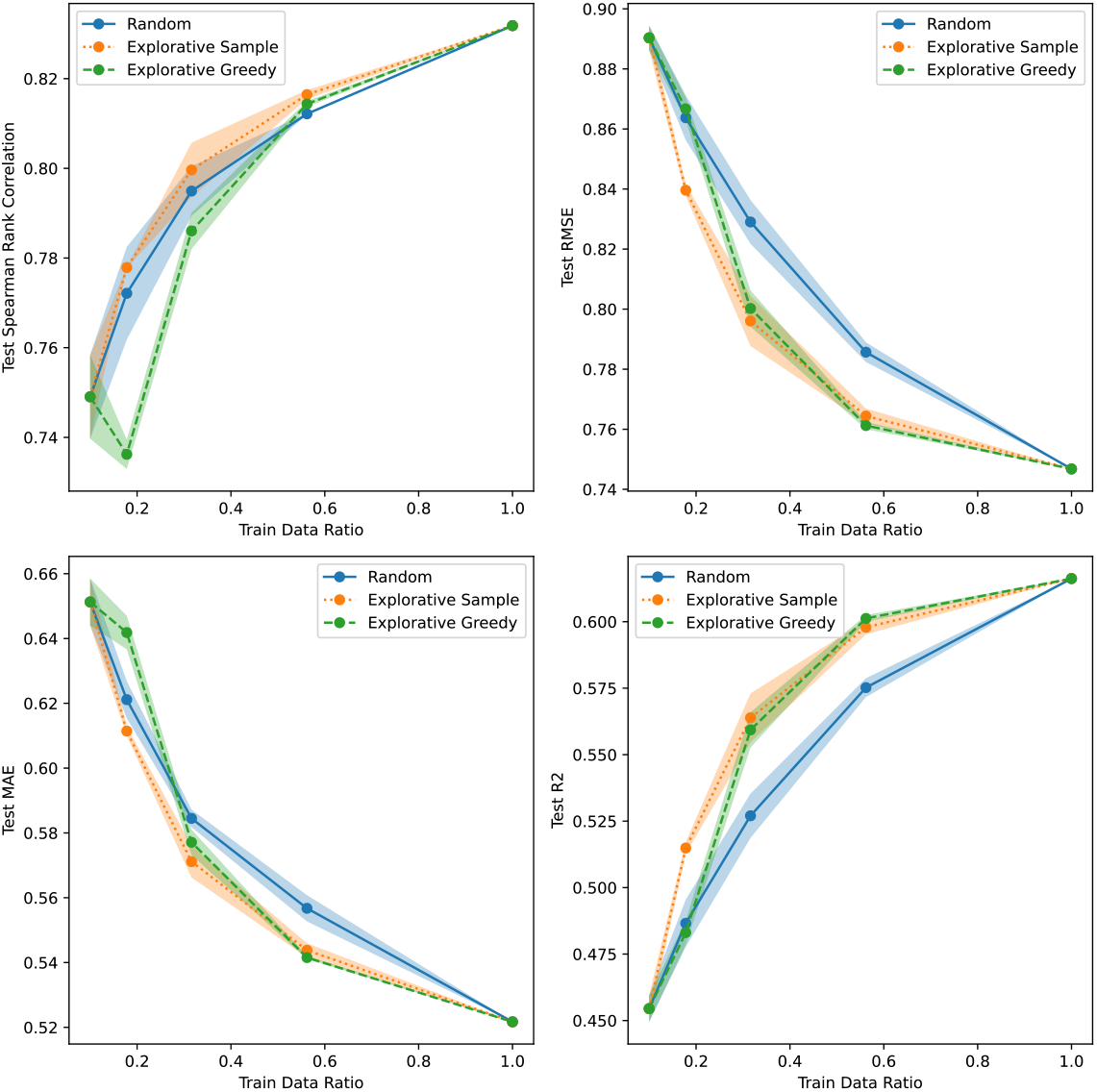
Active learning results for GB1/Random using CNN Ensemble uncertainty.

**Figure S32:**
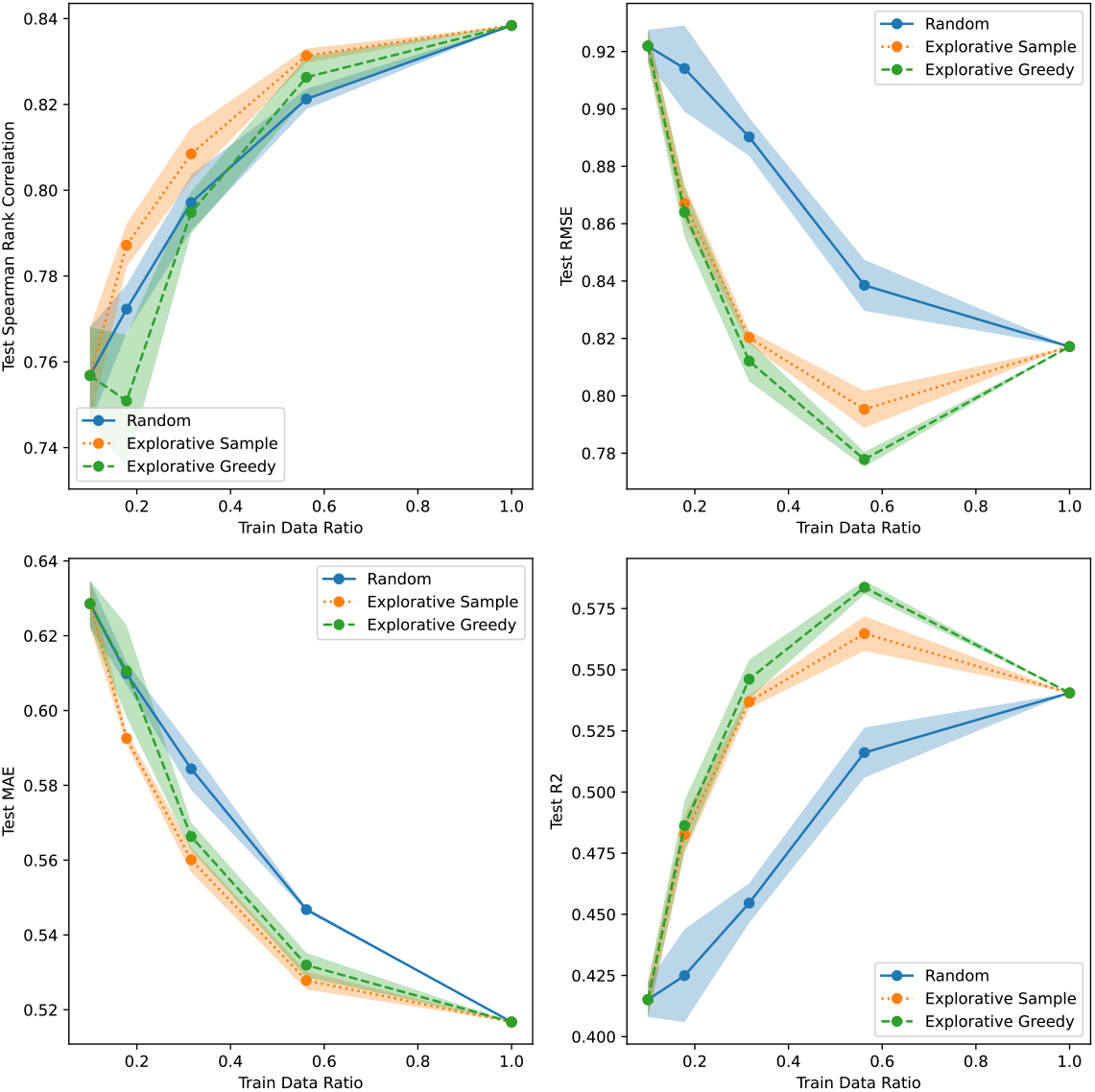
Active learning results for GB1/Random using CNN Evidential uncertainty.

**Figure S33:**
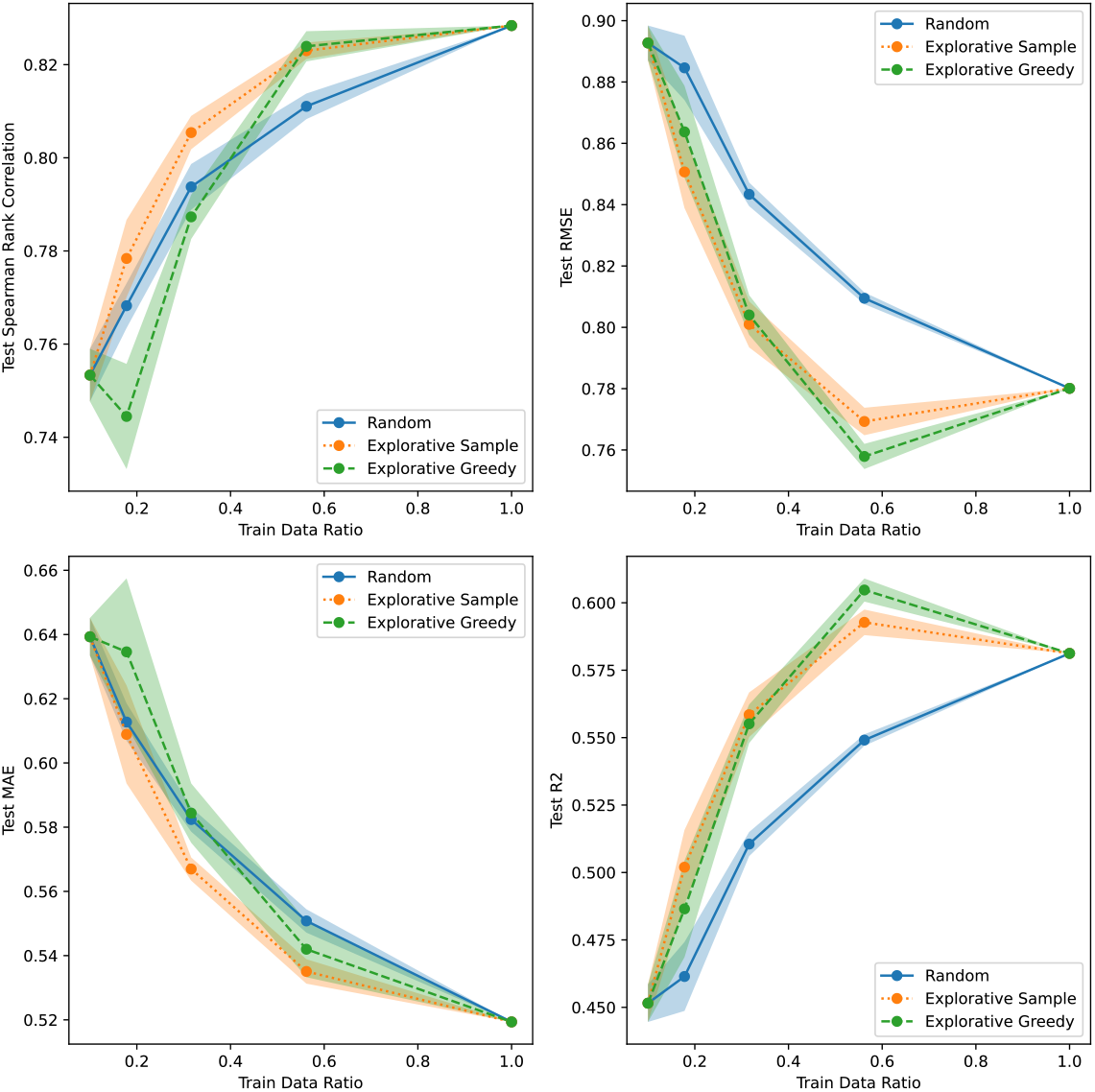
Active learning results for GB1/Random using CNN MVE uncertainty.

**Figure S34:**
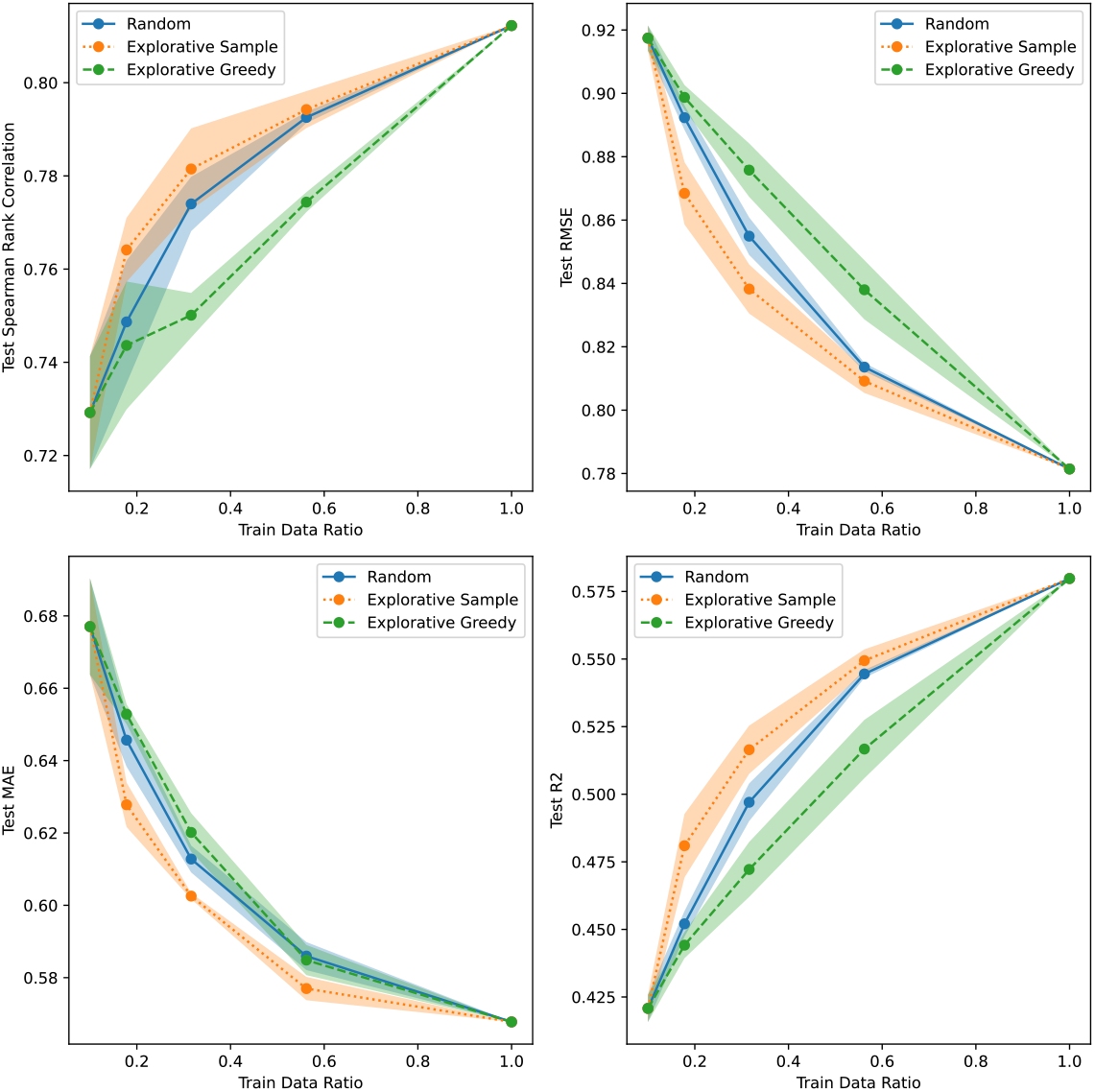
Active learning results for GB1/Random using CNN SVI uncertainty.

**Figure S35:**
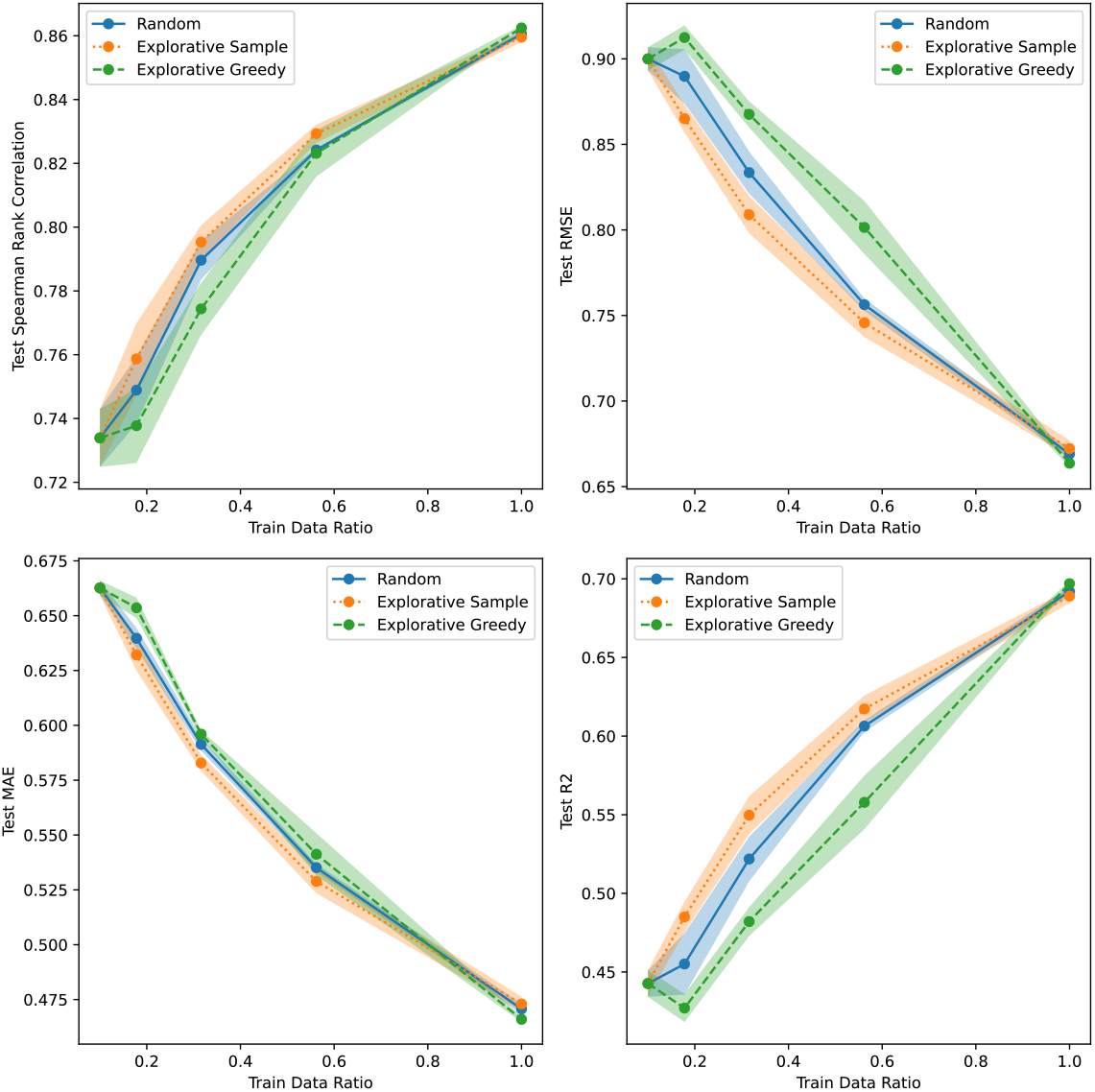
Active learning results for GB1/Random using GP Continuous uncertainty.

**Figure S36:**
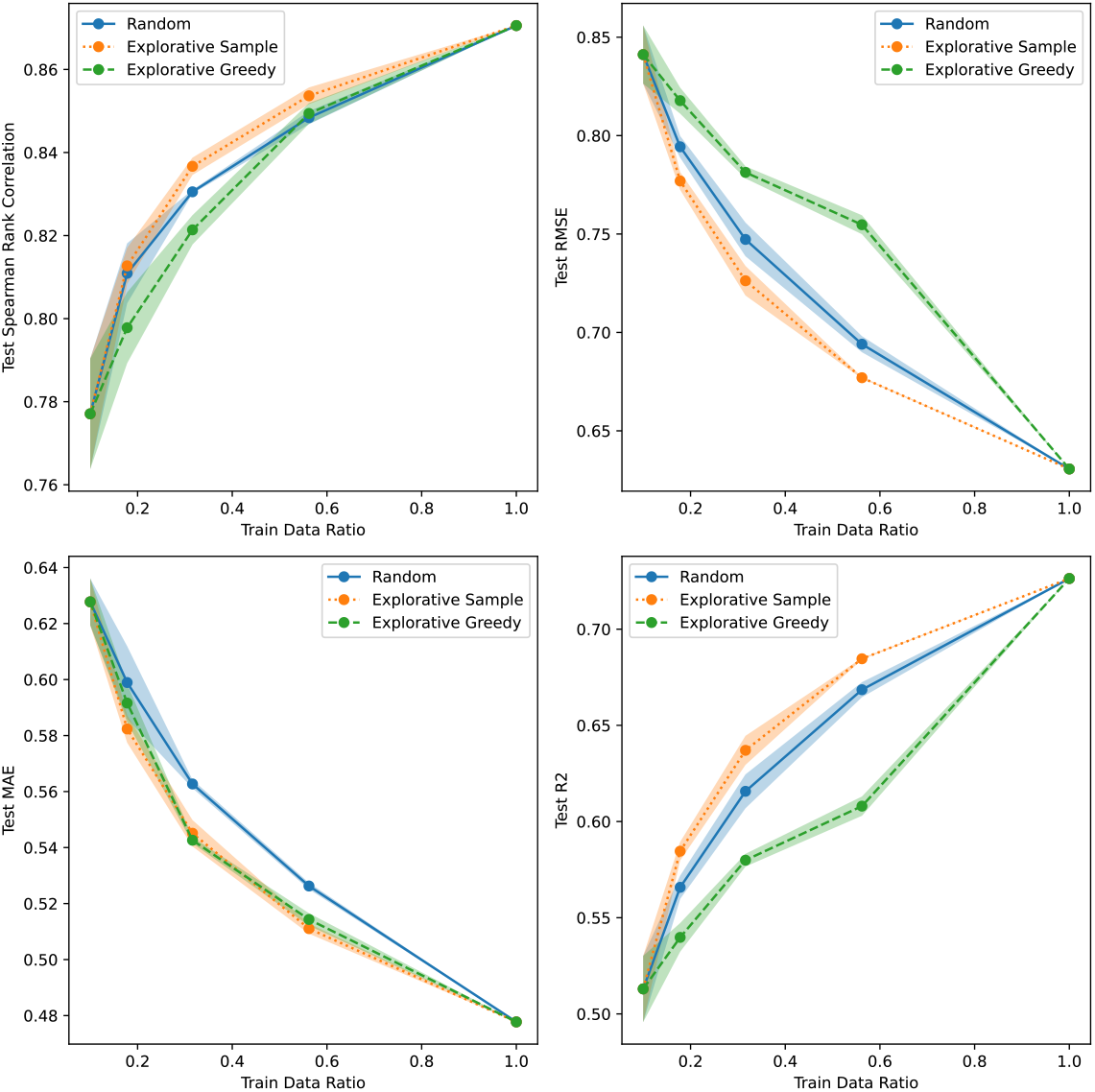
Active learning results for GB1/Random using Linear Bayesian Ridge uncertainty.

**Figure S37:**
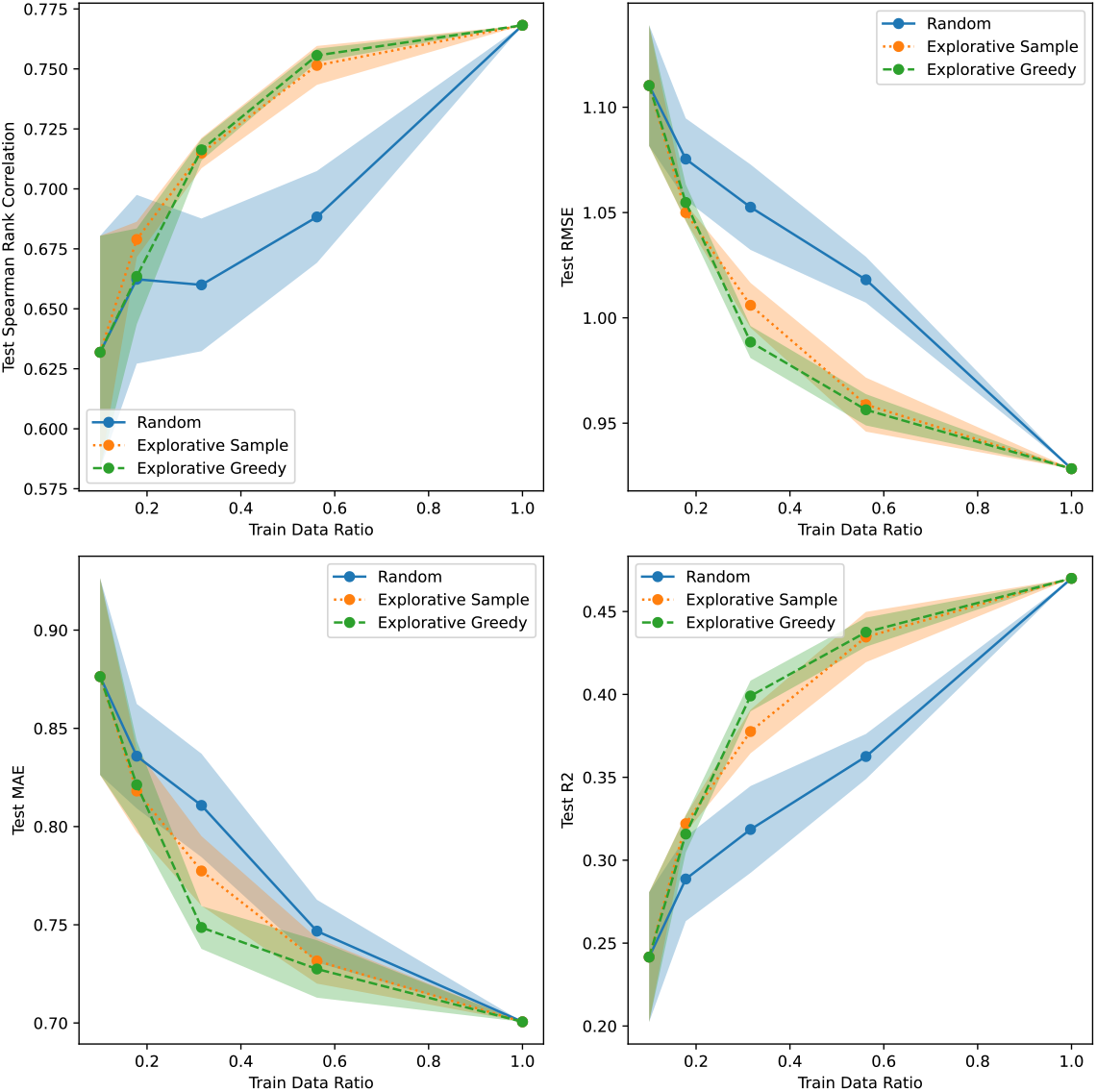
Active learning results for GB1/3 vs. Rest using CNN Dropout uncertainty.

**Figure S38:**
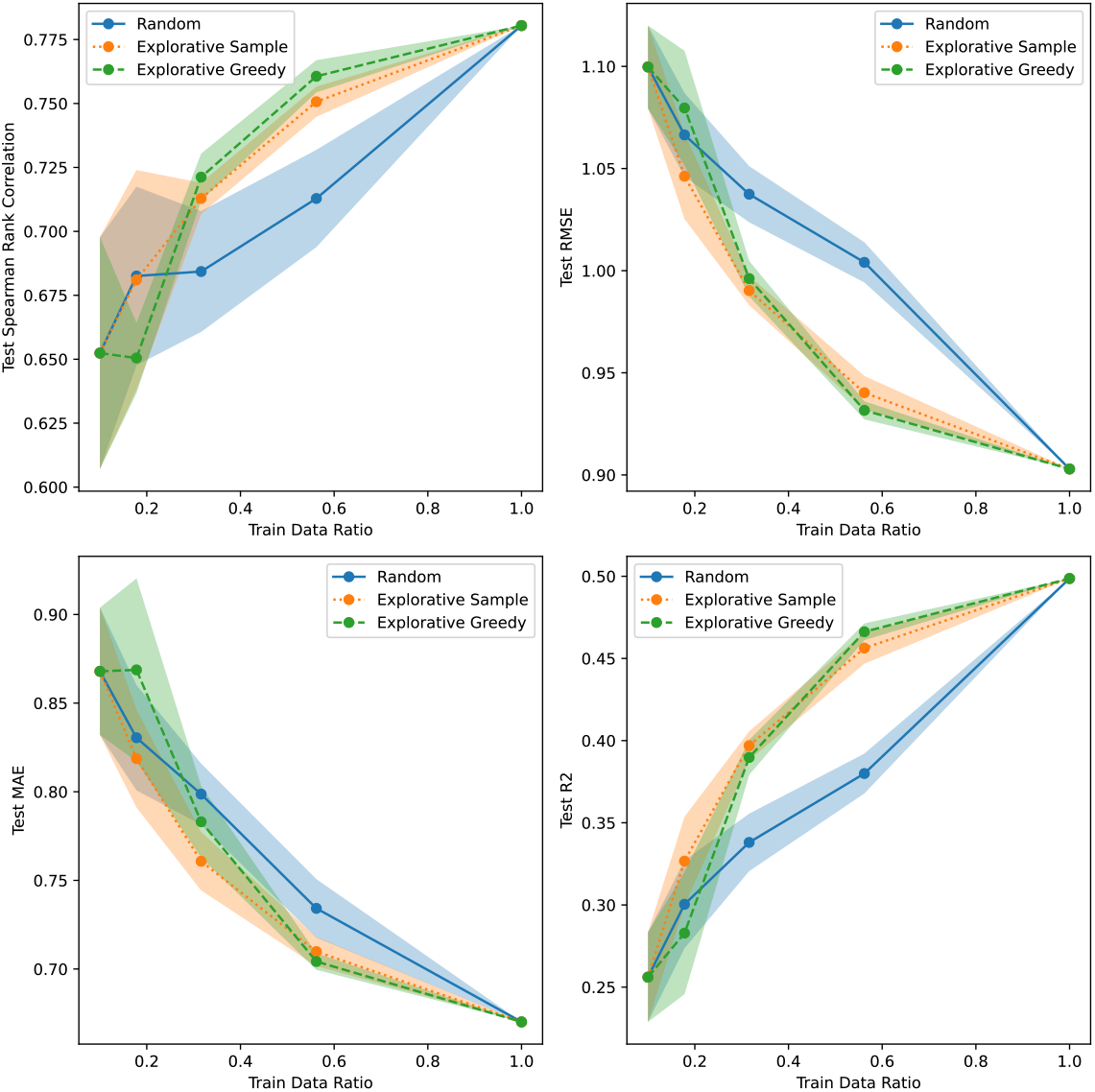
Active learning results for GB1/3 vs. Rest using CNN Ensemble uncertainty.

**Figure S39:**
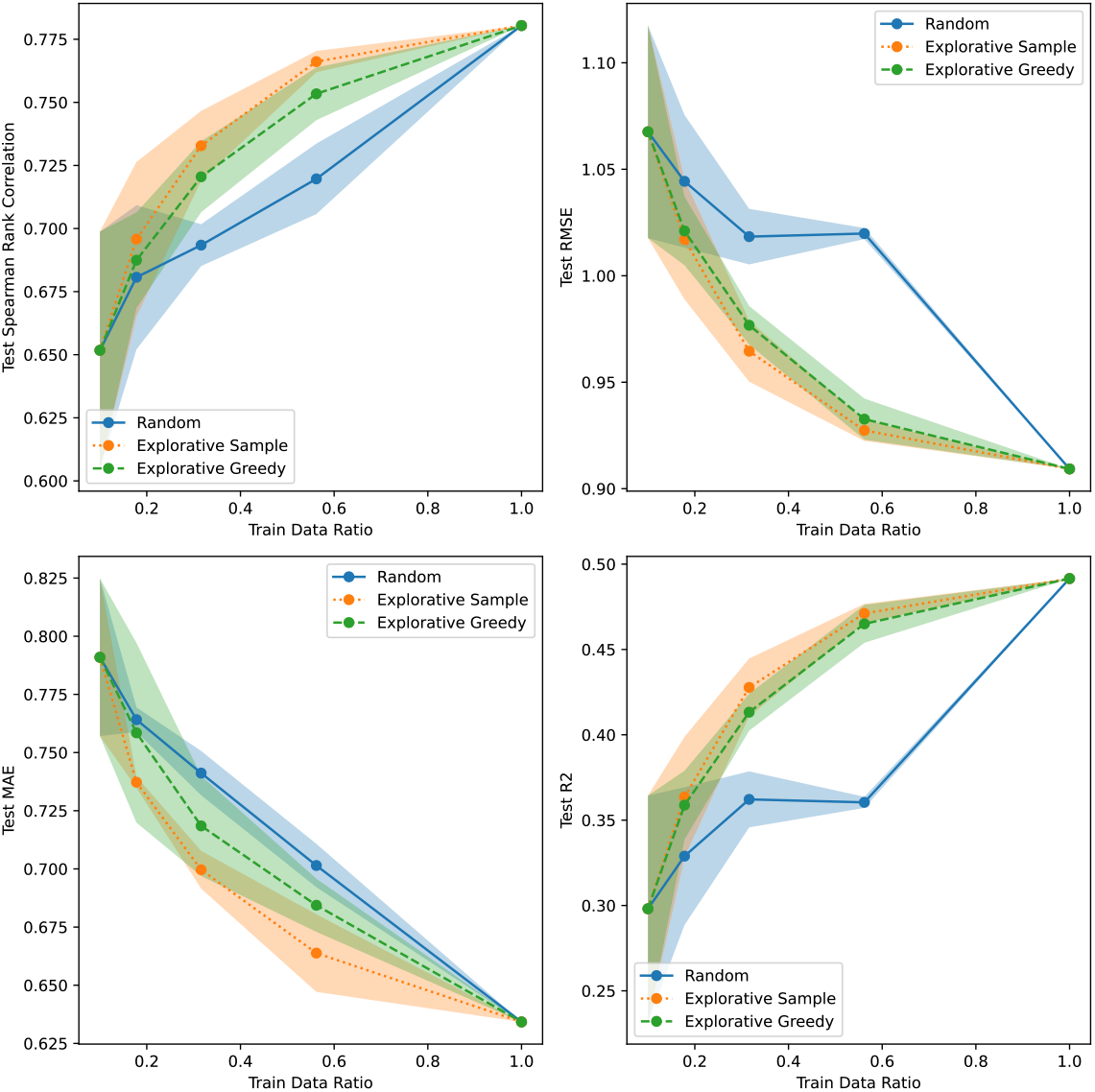
Active learning results for GB1/3 vs. Rest using CNN Evidential uncertainty.

**Figure S40:**
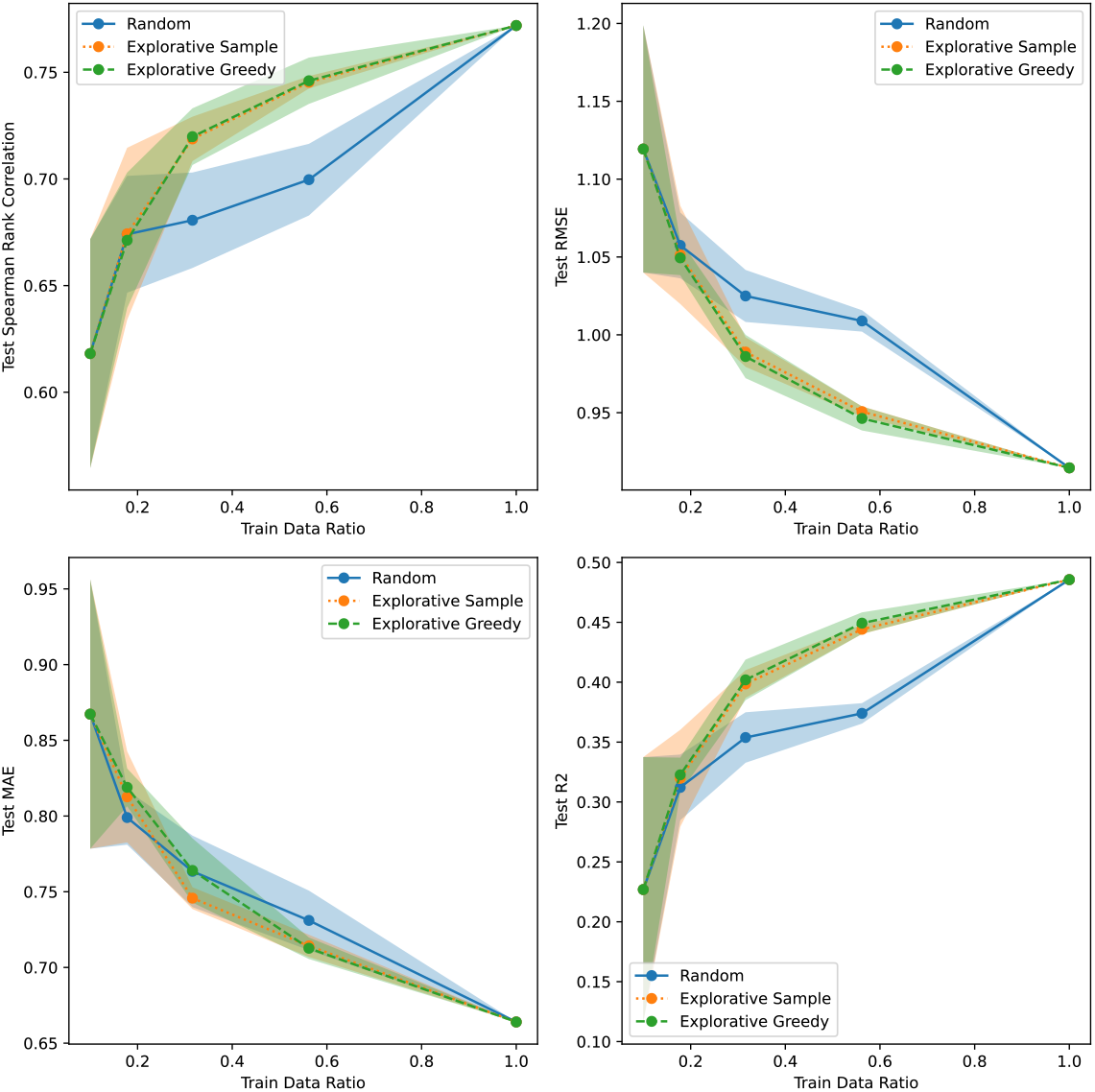
Active learning results for GB1/3 vs. Rest using CNN MVE uncertainty.

**Figure S41:**
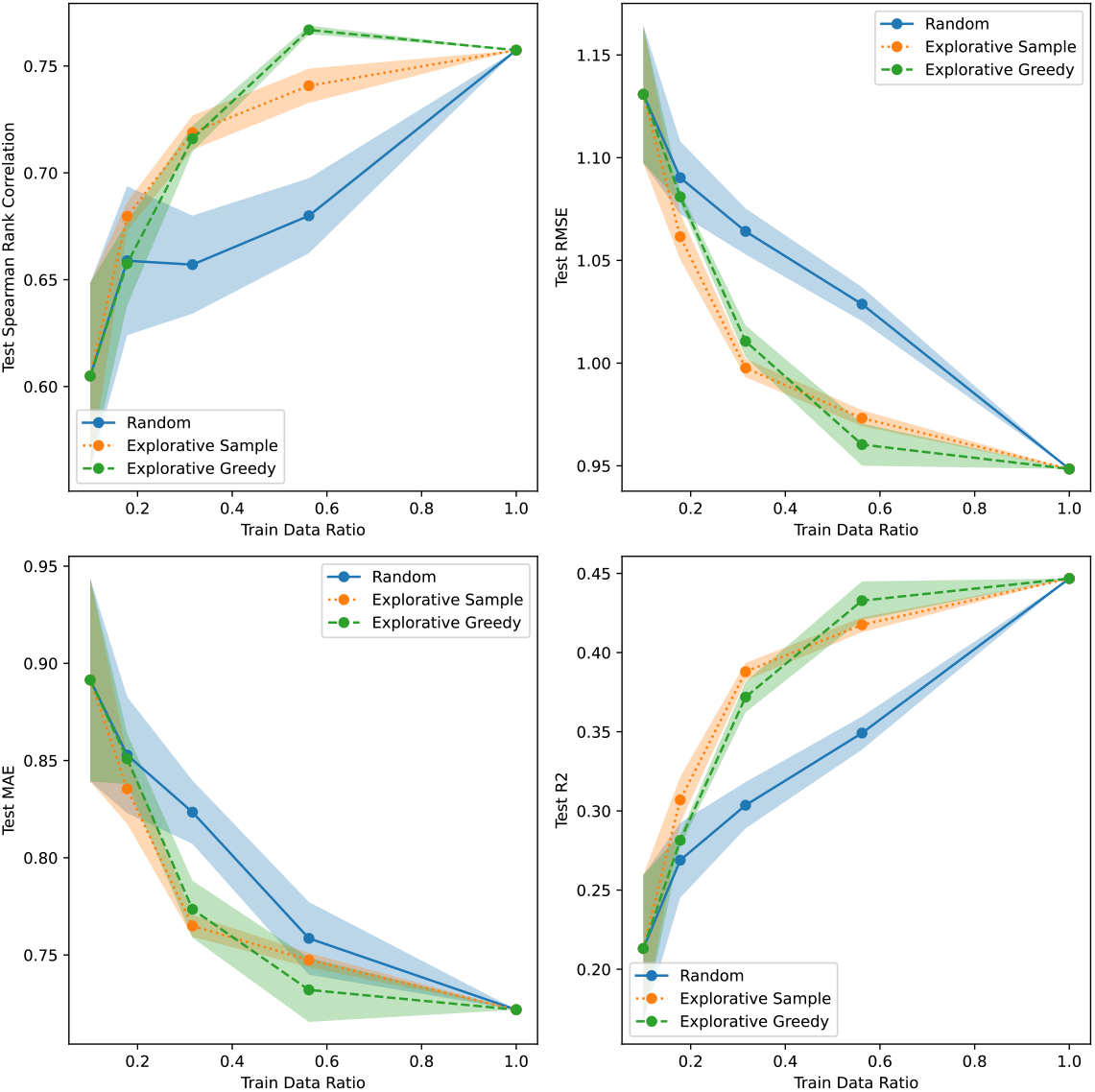
Active learning results for GB1/3 vs. Rest using CNN SVI uncertainty.

**Figure S42:**
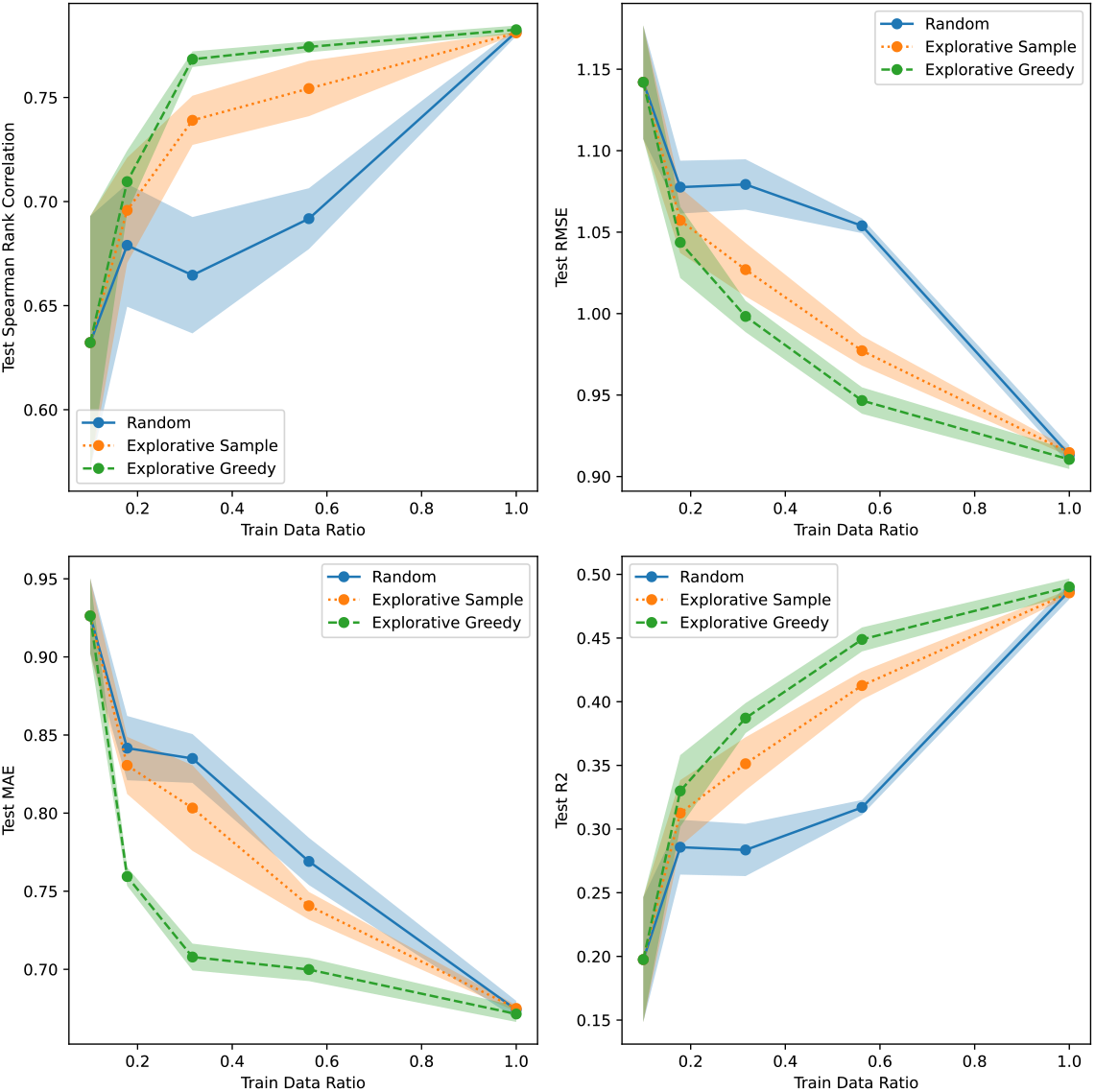
Active learning results for GB1/3 vs. Rest using GP Continuous uncertainty.

**Figure S43:**
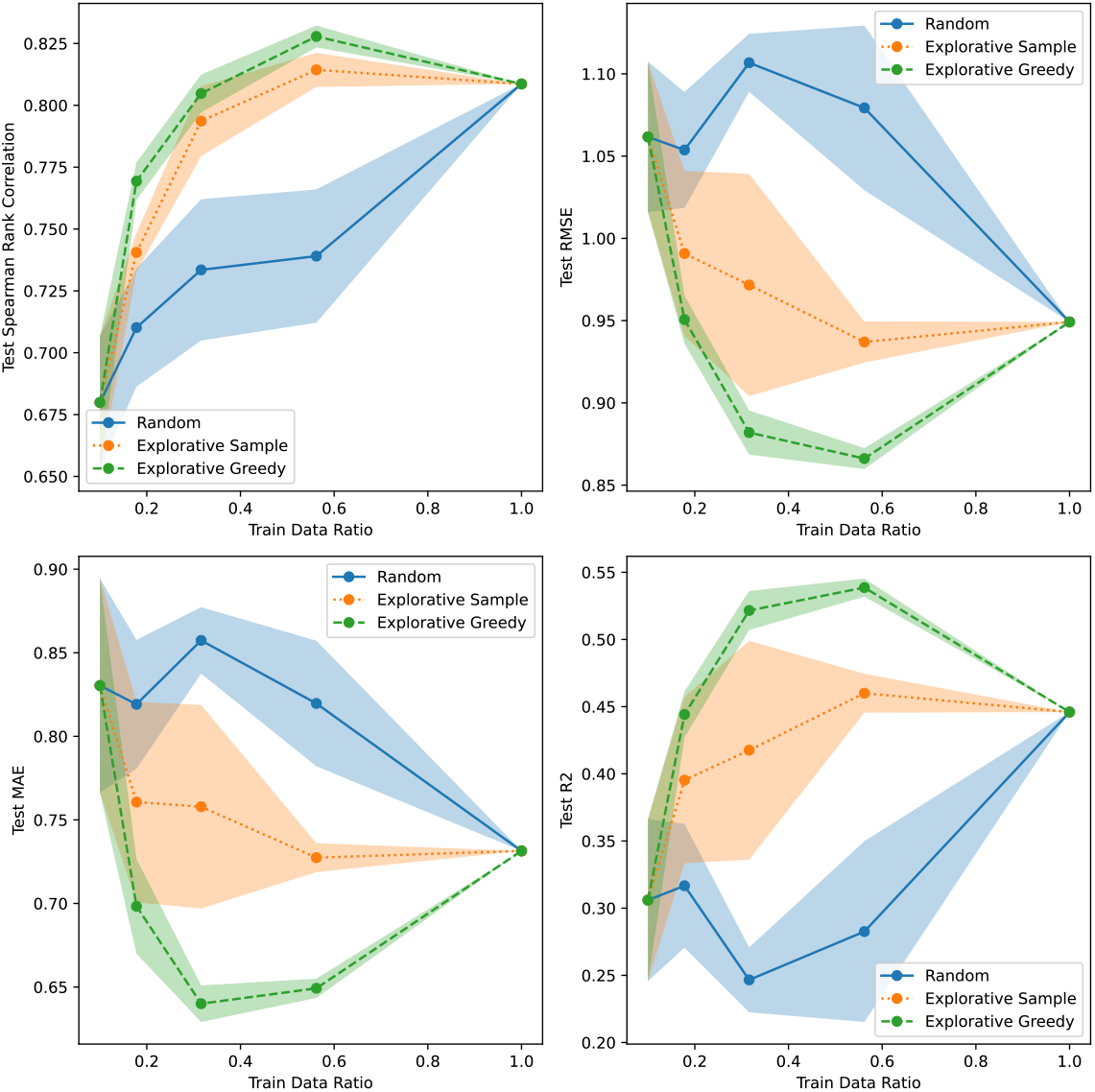
Active learning results for GB1/3 vs. Rest using Linear Bayesian Ridge uncertainty.

**Figure S44:**
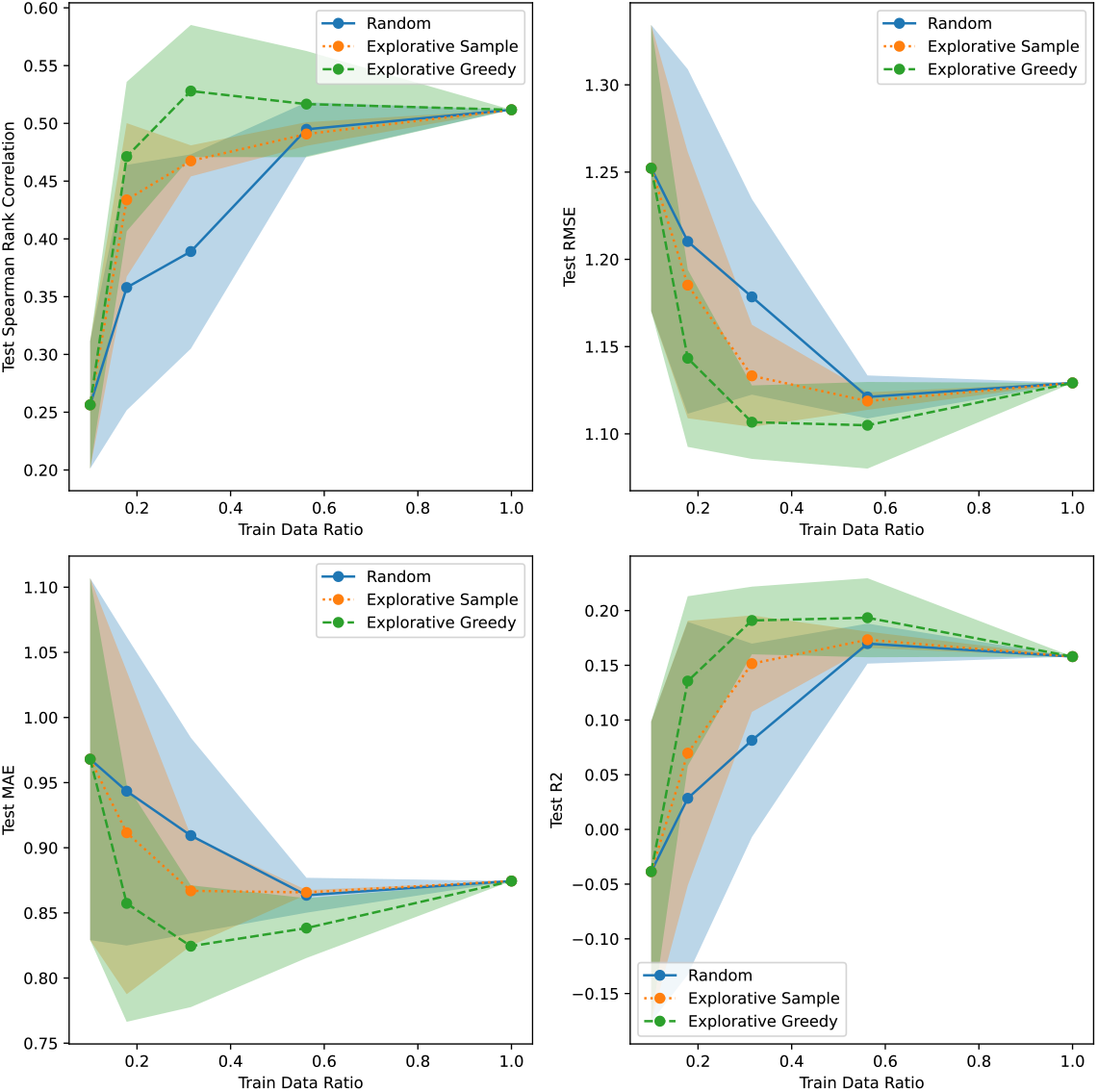
Active learning results for GB1/2 vs. Rest using CNN Dropout uncertainty.

**Figure S45:**
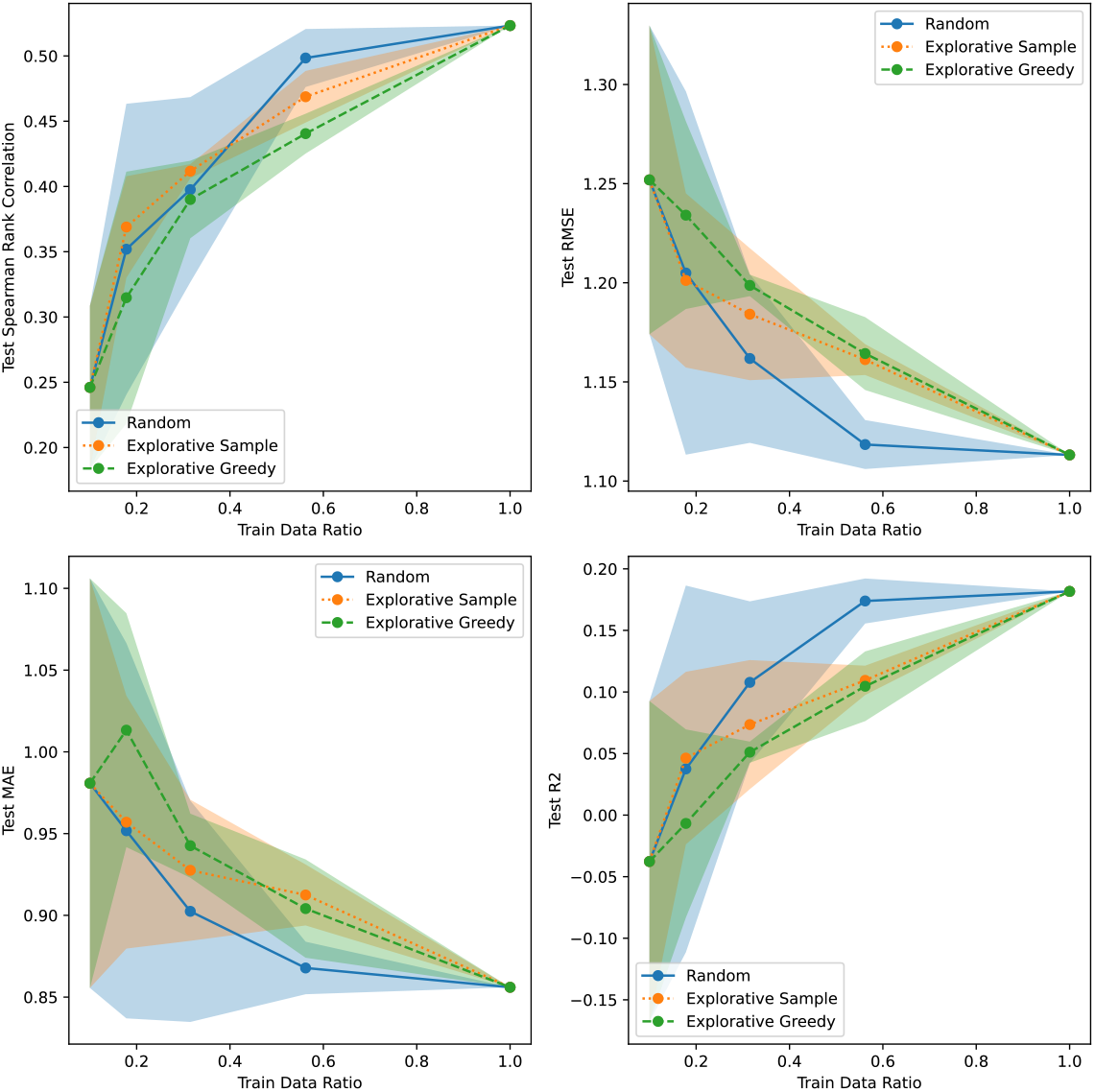
Active learning results for GB1/2 vs. Rest using CNN Ensemble uncertainty.

**Figure S46:**
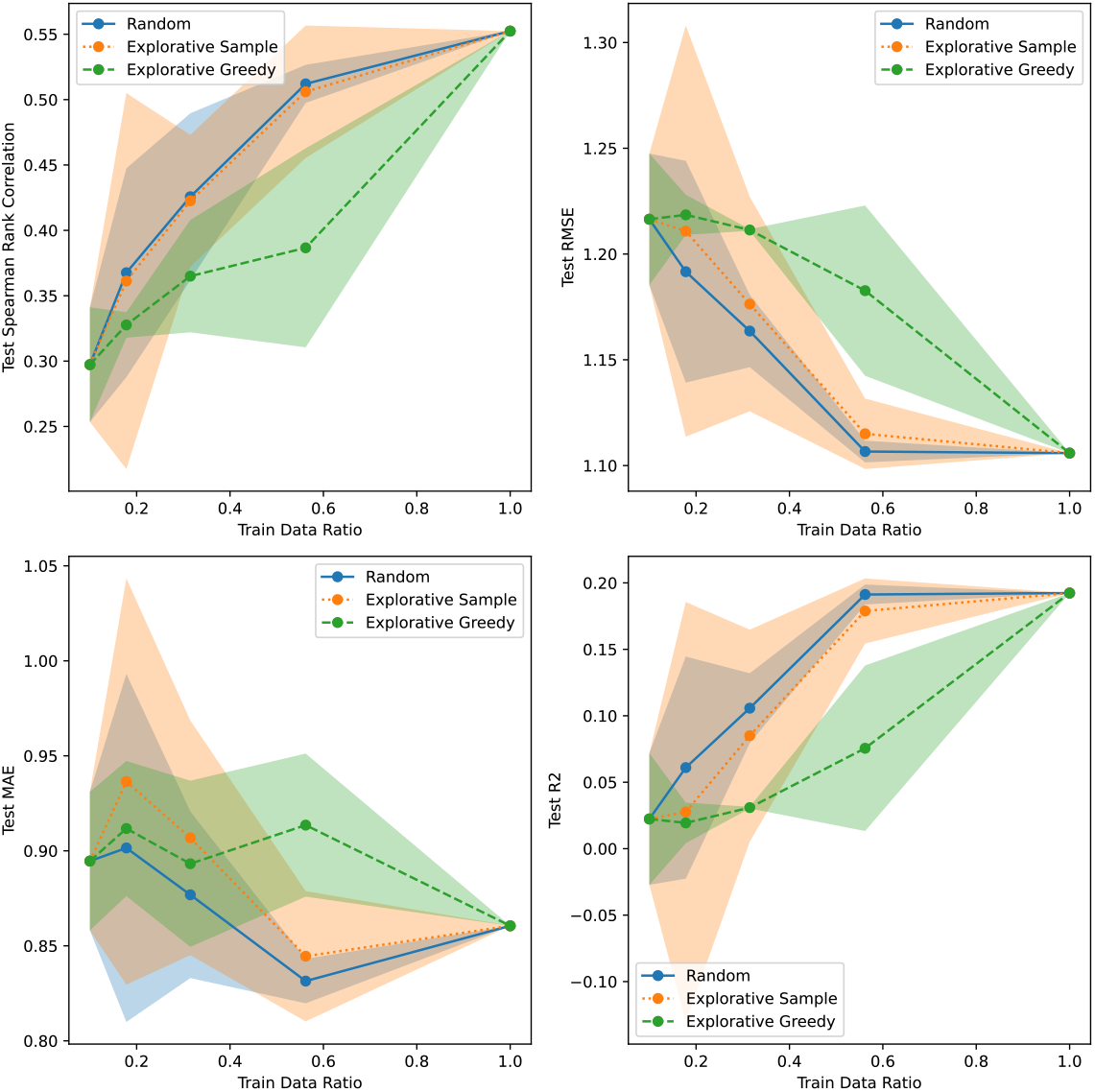
Active learning results for GB1/2 vs. Rest using CNN Evidential uncertainty.

**Figure S47:**
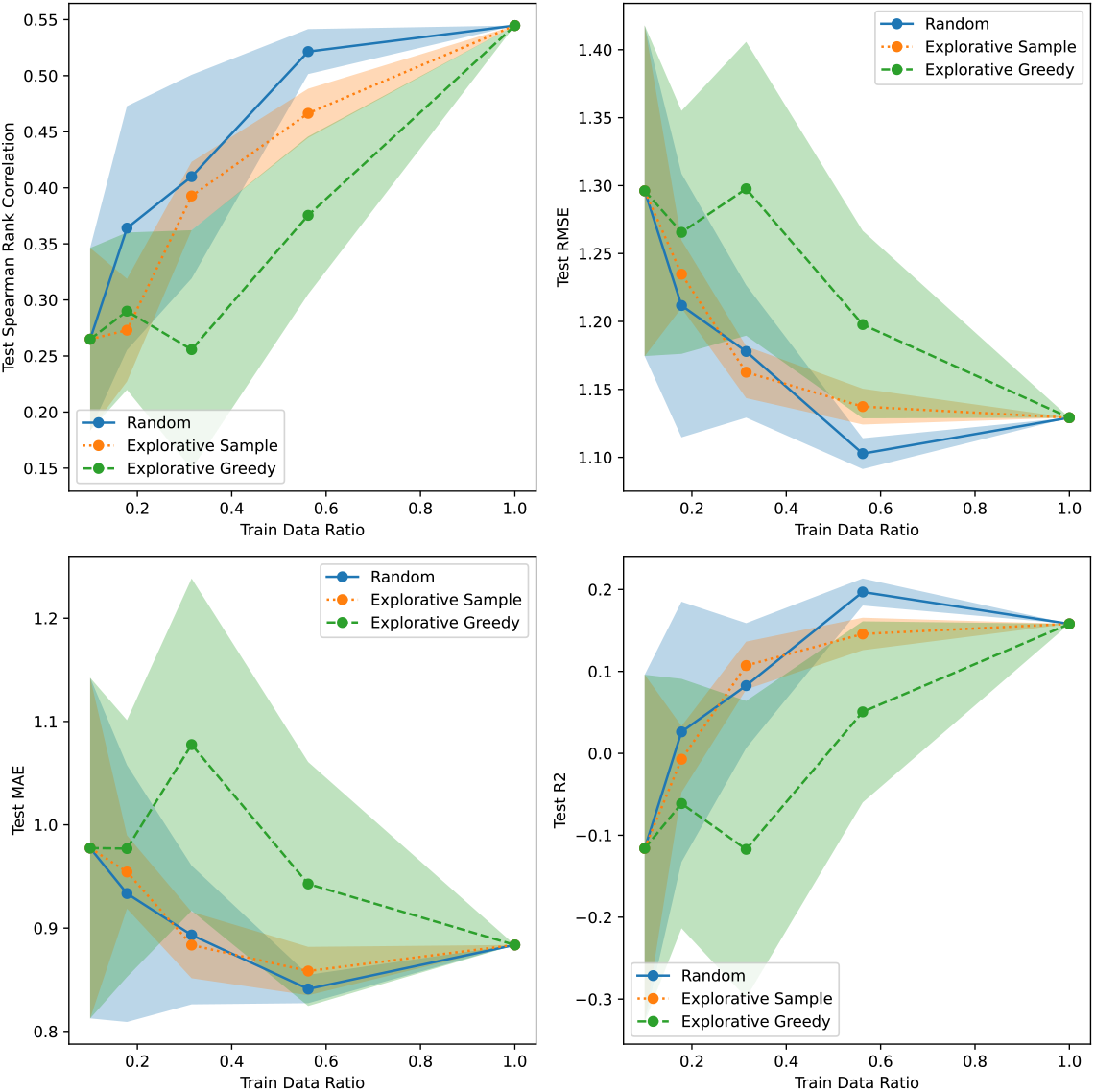
Active learning results for GB1/2 vs. Rest using CNN MVE uncertainty.

**Figure S48:**
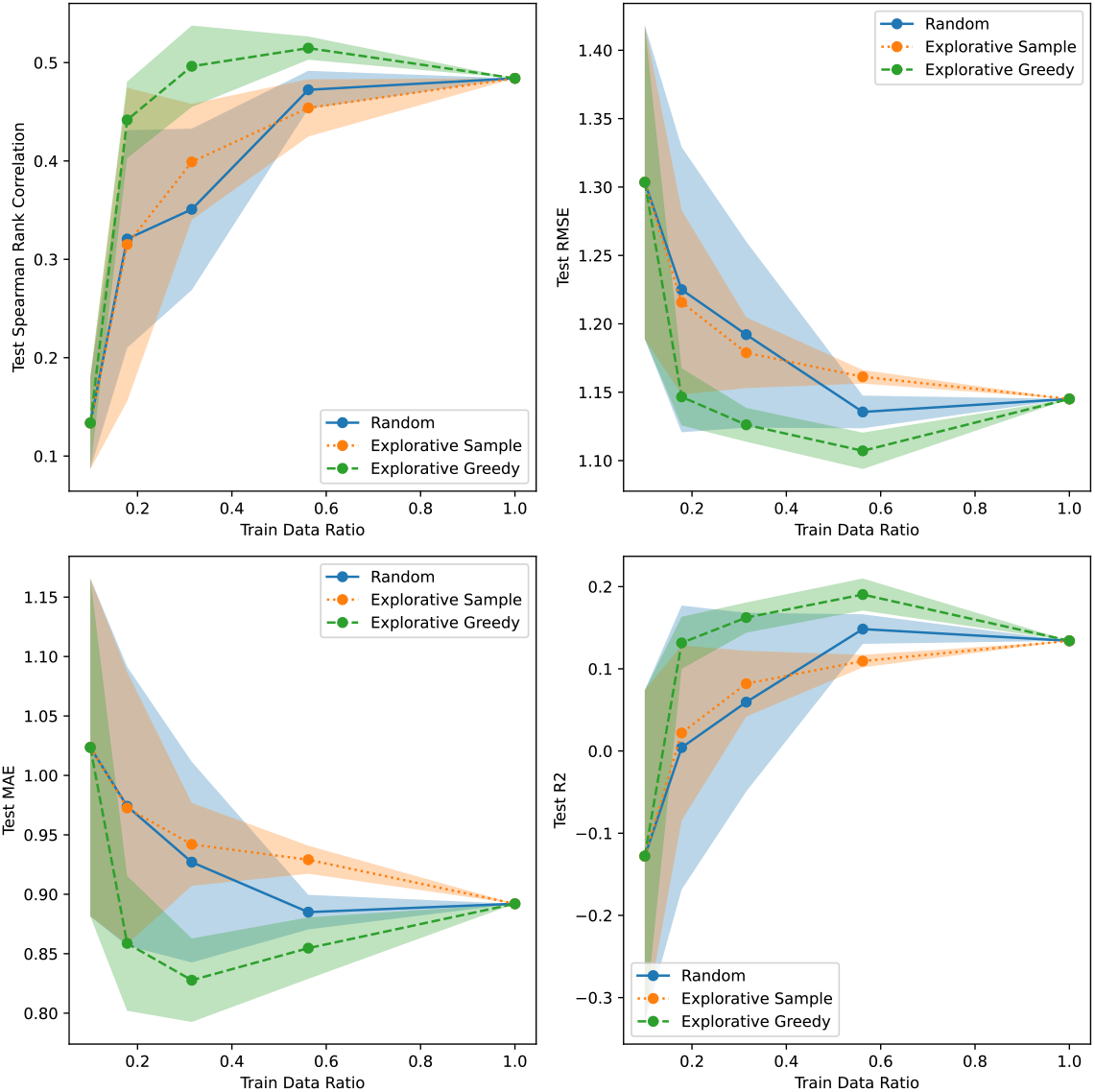
Active learning results for GB1/2 vs. Rest using CNN SVI uncertainty.

**Figure S49:**
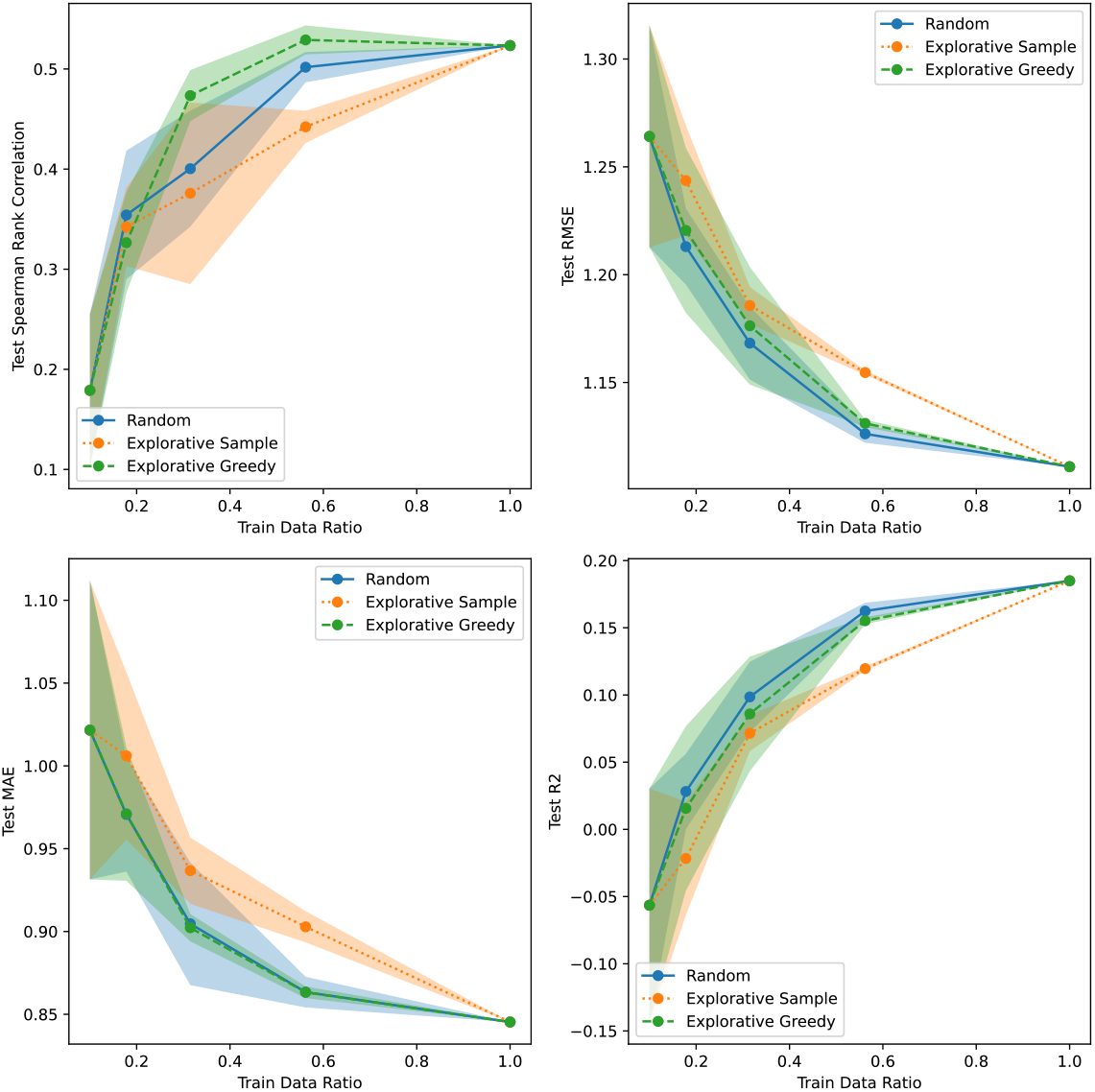
Active learning results for GB1/2 vs. Rest using GP Continuous uncertainty.

**Figure S50:**
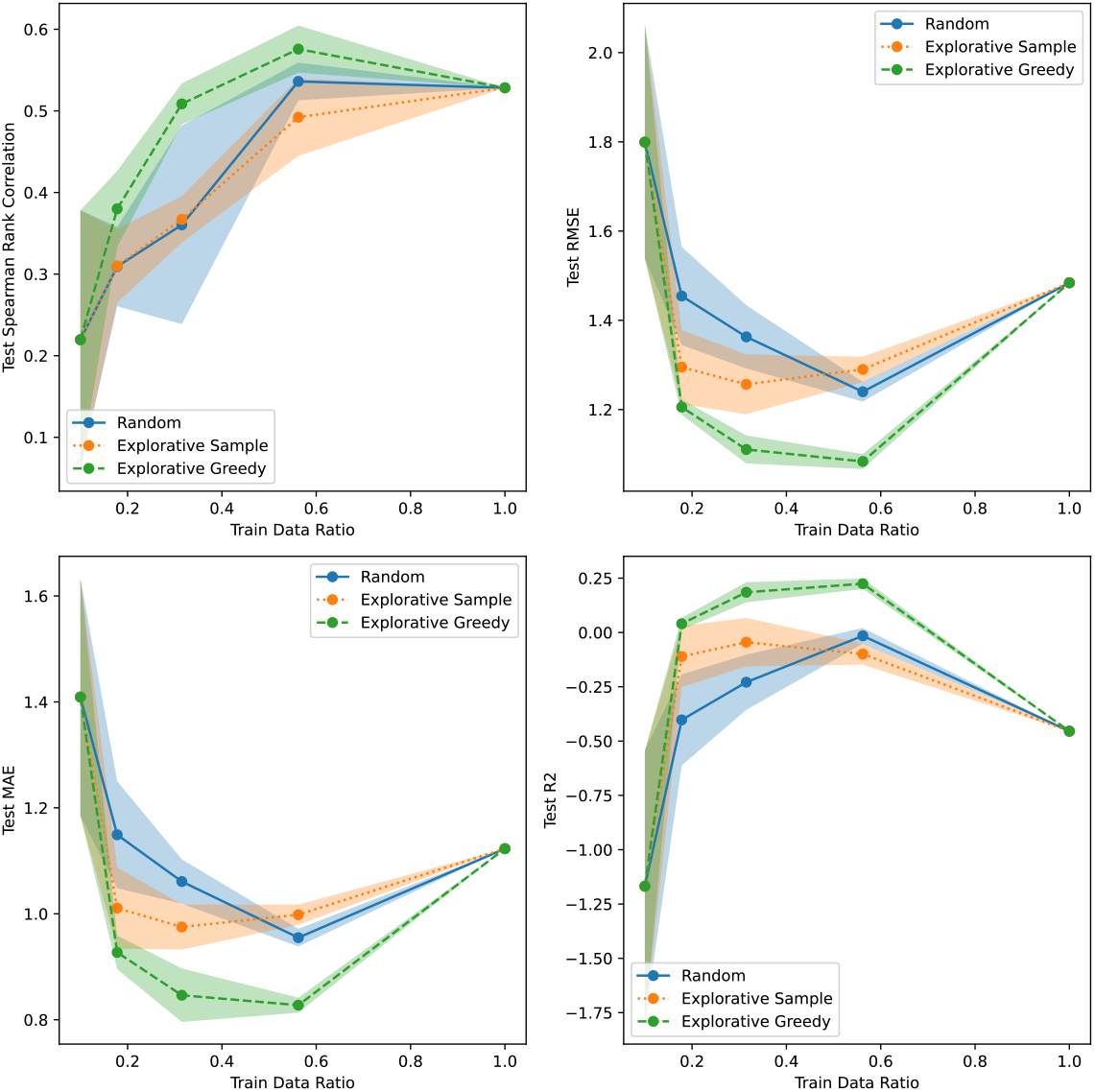
Active learning results for GB1/2 vs. Rest using Linear Bayesian Ridge uncertainty.

**Figure S51:**
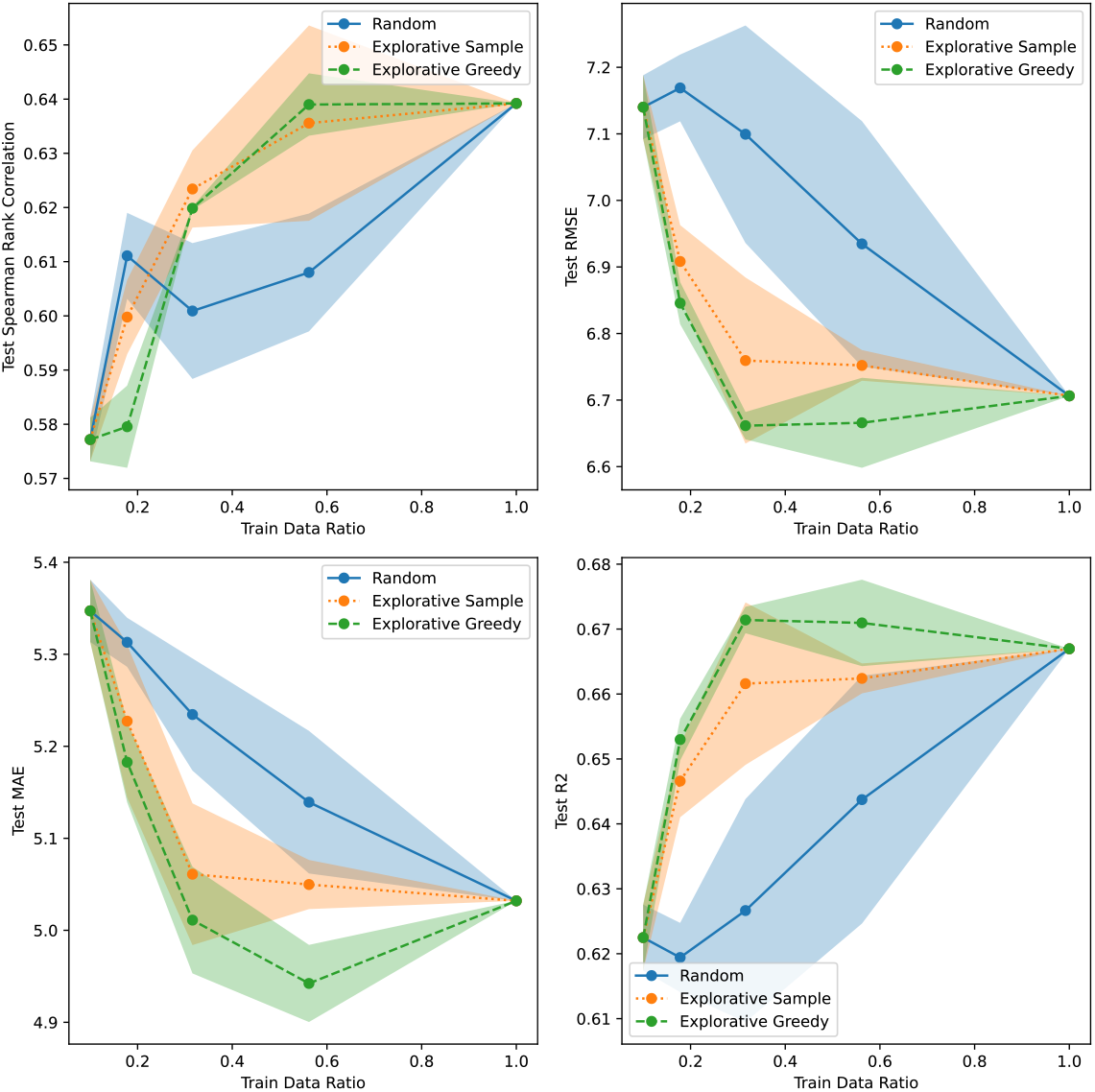
Active learning results for Meltome/Random using CNN Dropout uncertainty.

**Figure S52:**
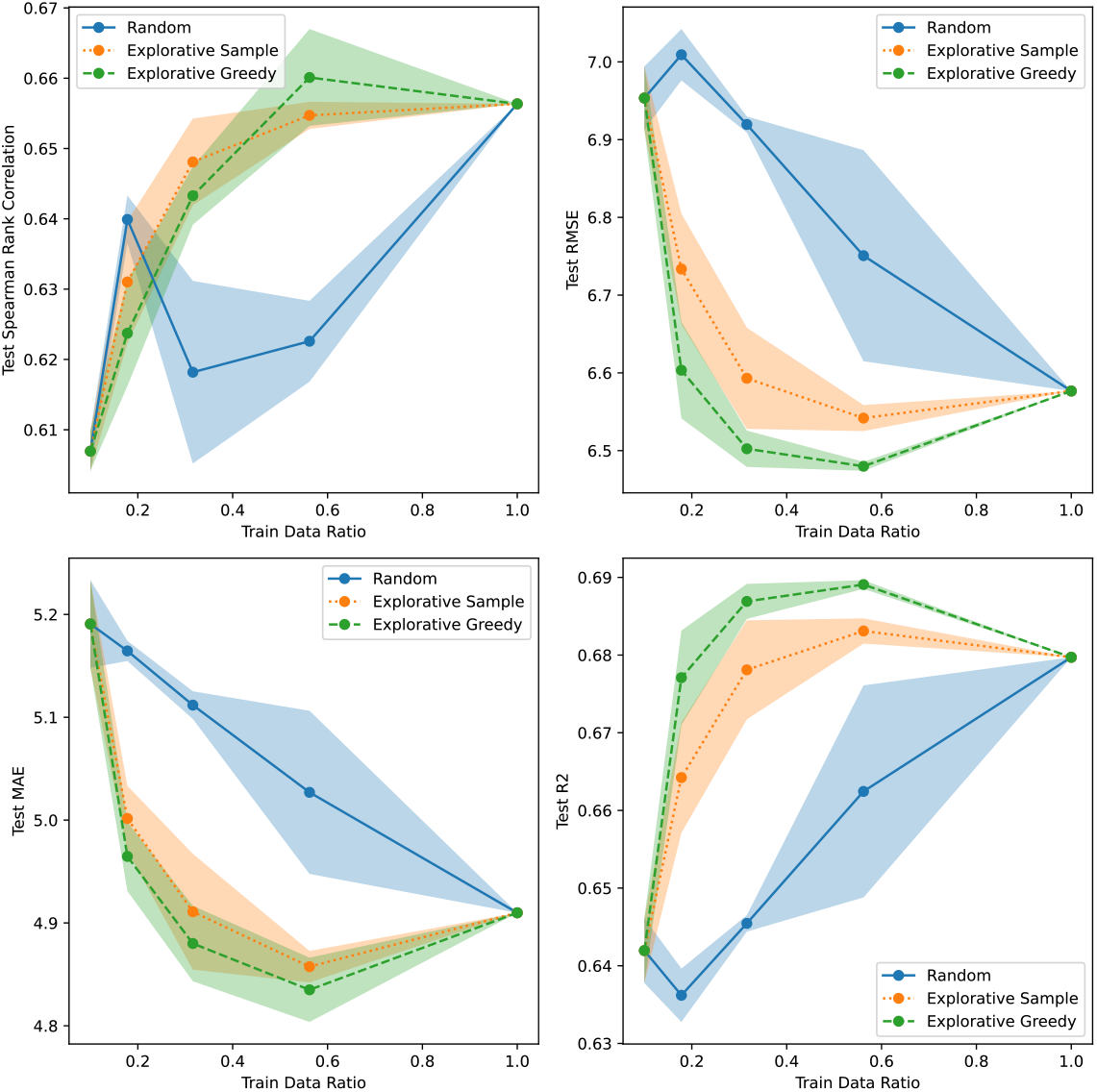
Active learning results for Meltome/Random using CNN Ensemble uncertainty.

**Figure S53:**
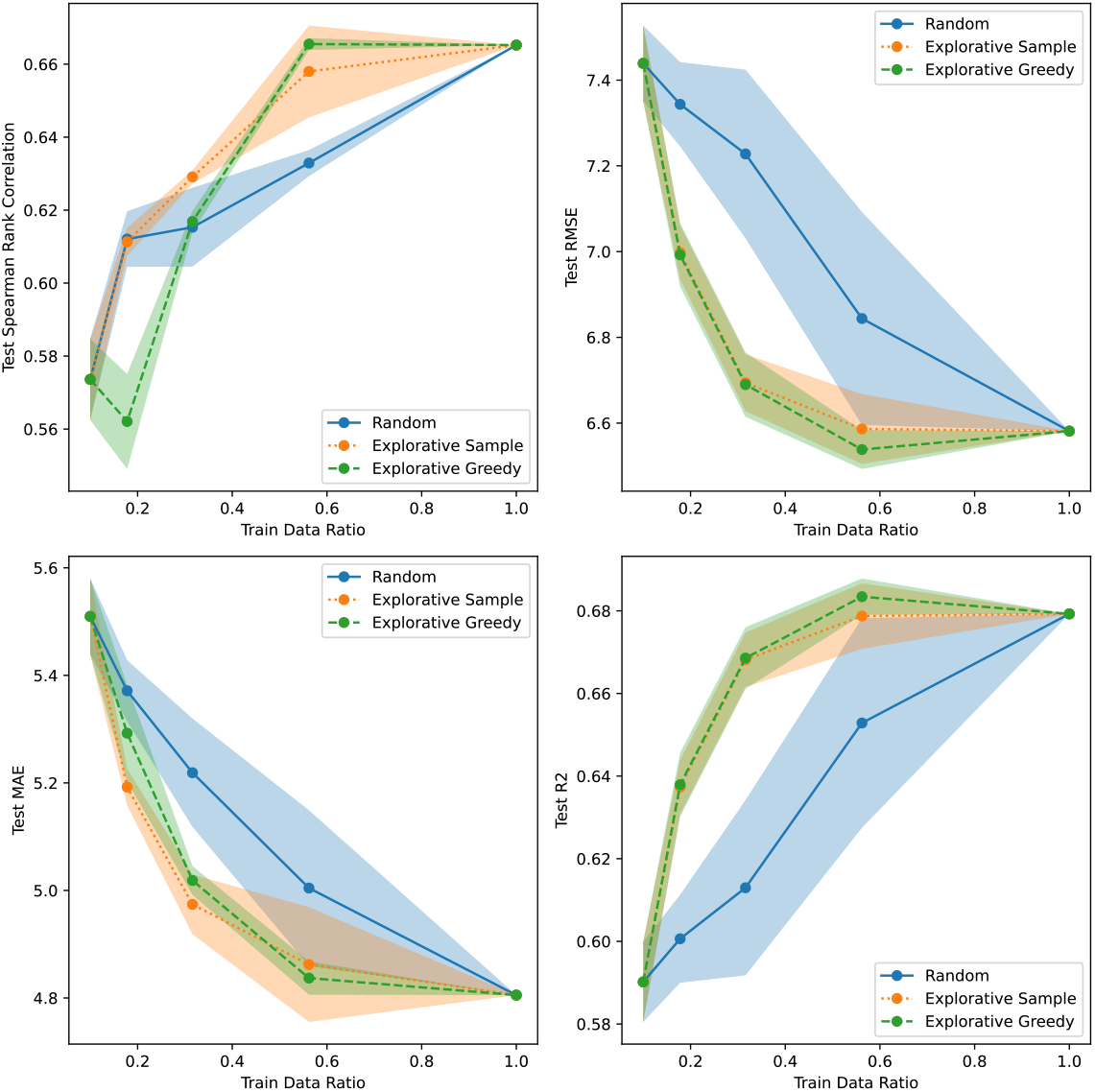
Active learning results for Meltome/Random using CNN Evidential uncertainty.

**Figure S54:**
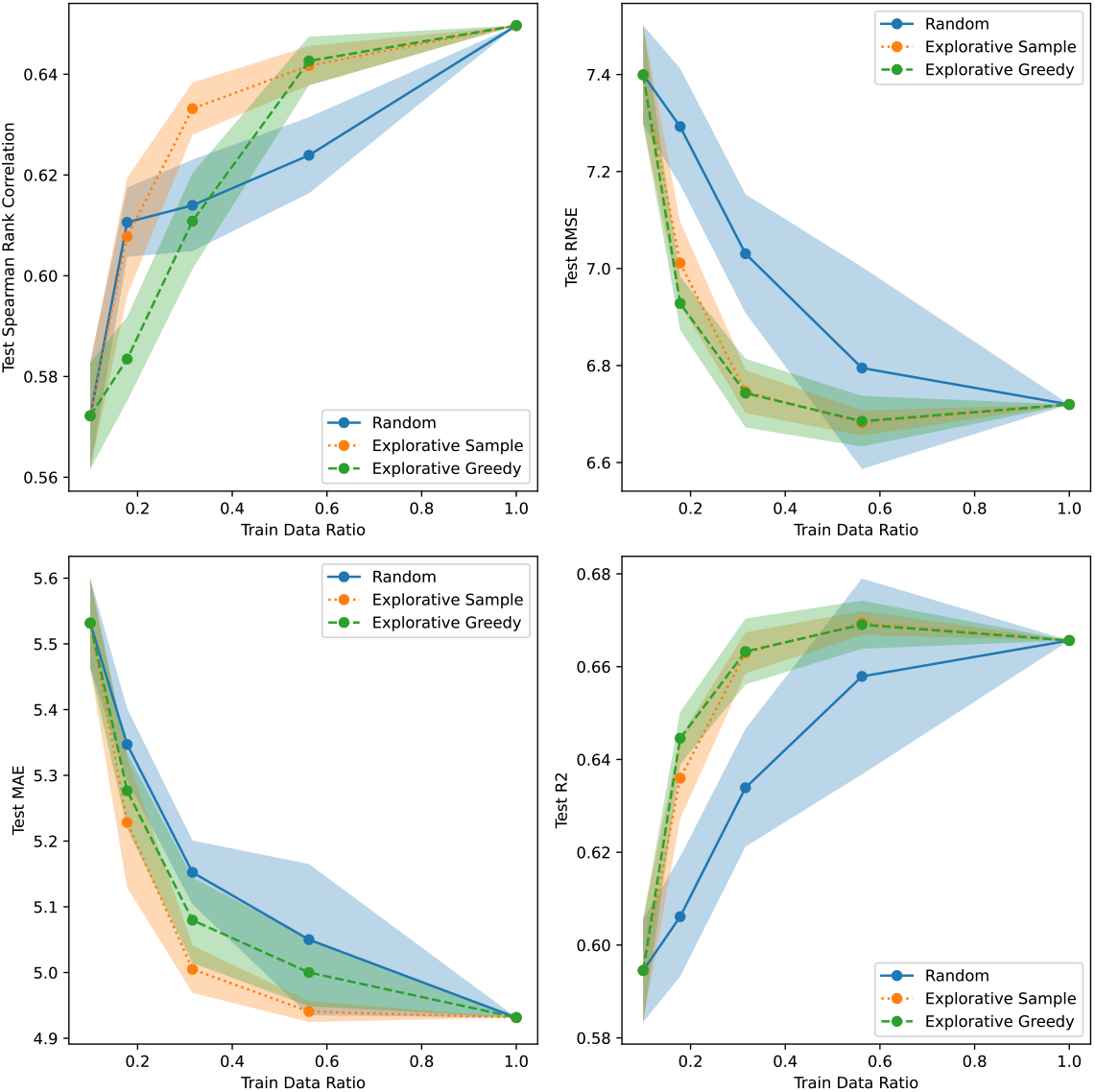
Active learning results for Meltome/Random using CNN MVE uncertainty.

**Figure S55:**
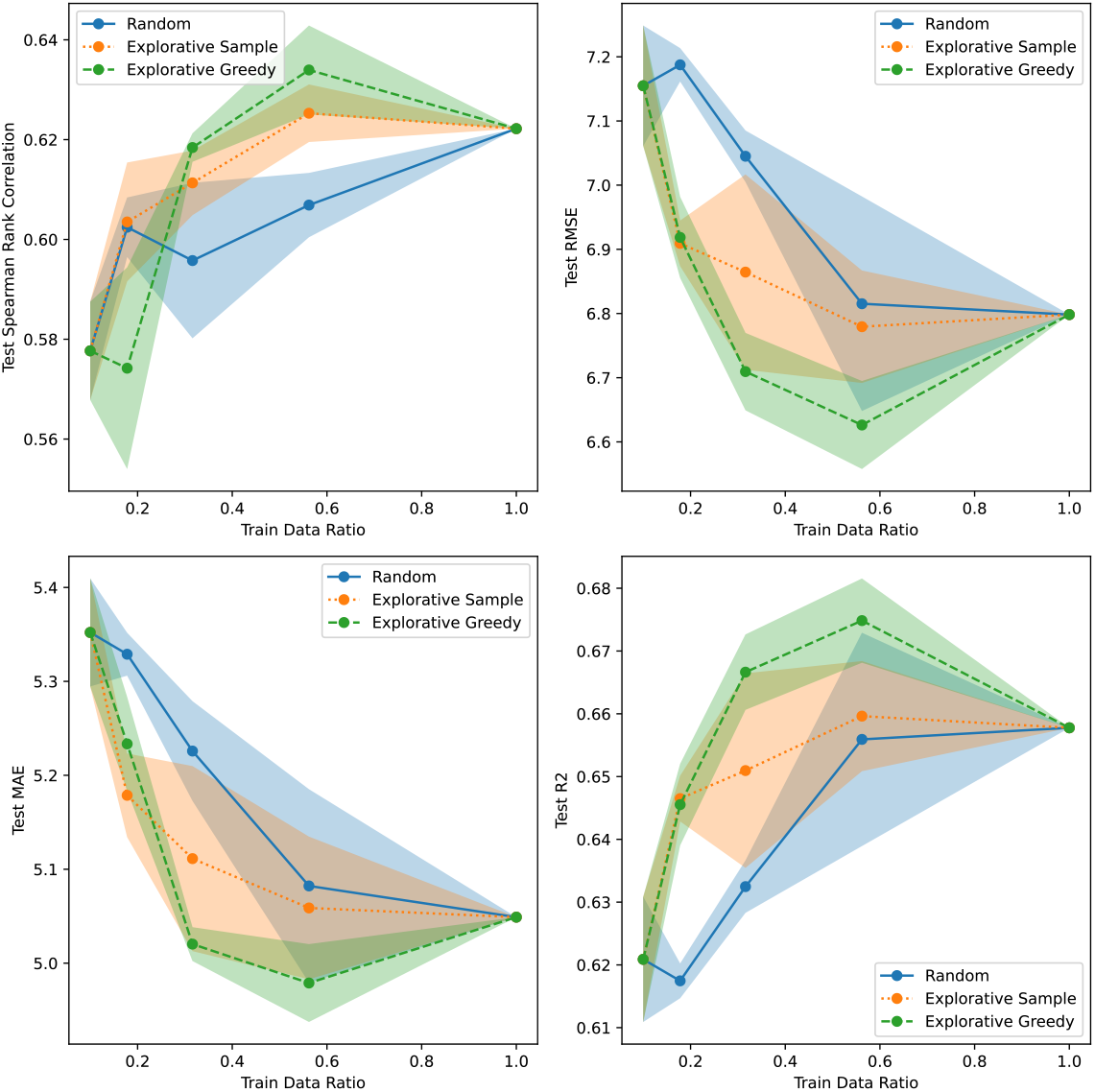
Active learning results for Meltome/Random using CNN SVI uncertainty.

**Figure S56:**
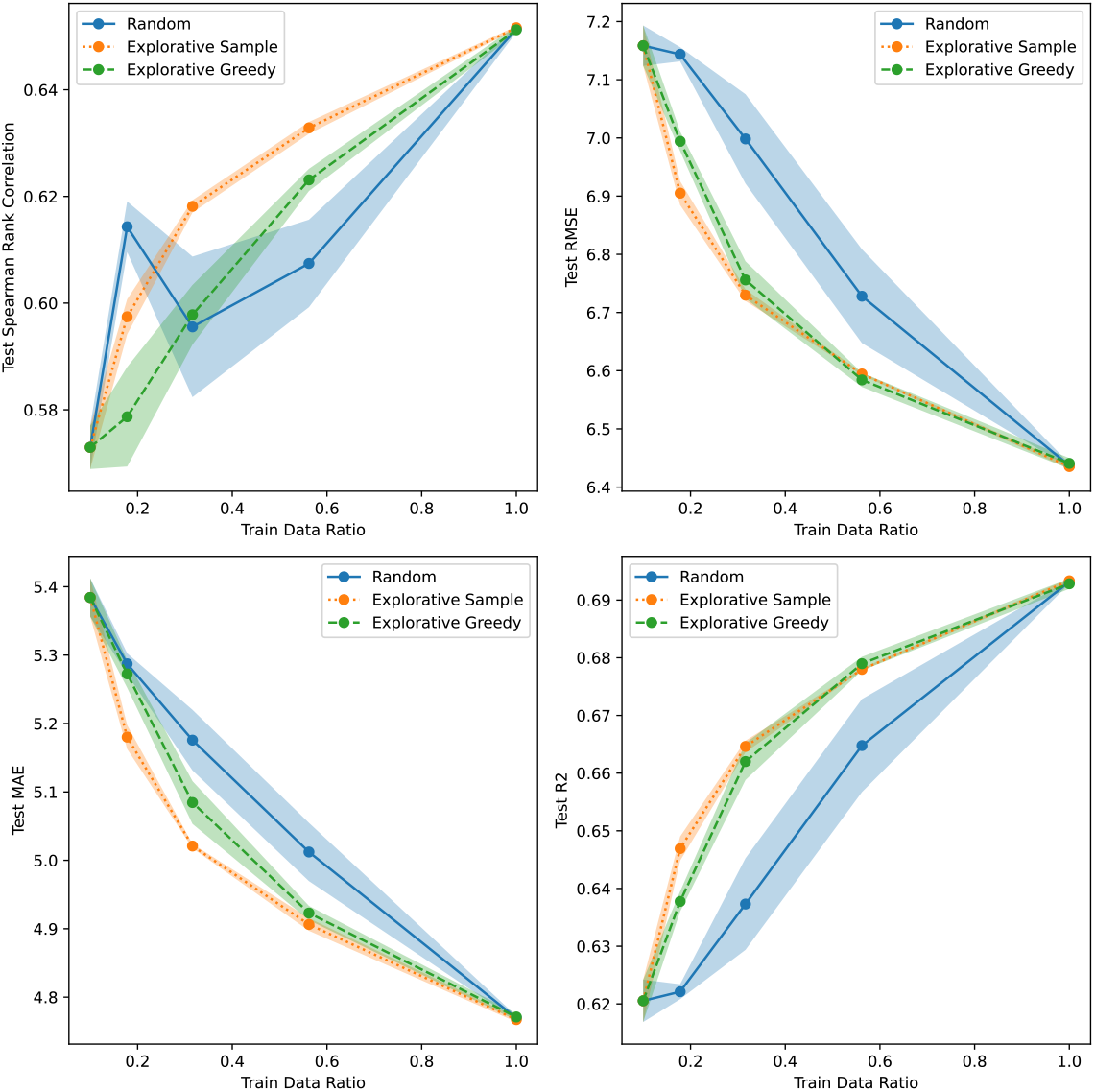
Active learning results for Meltome/Random using GP Continuous uncertainty.

**Figure S57:**
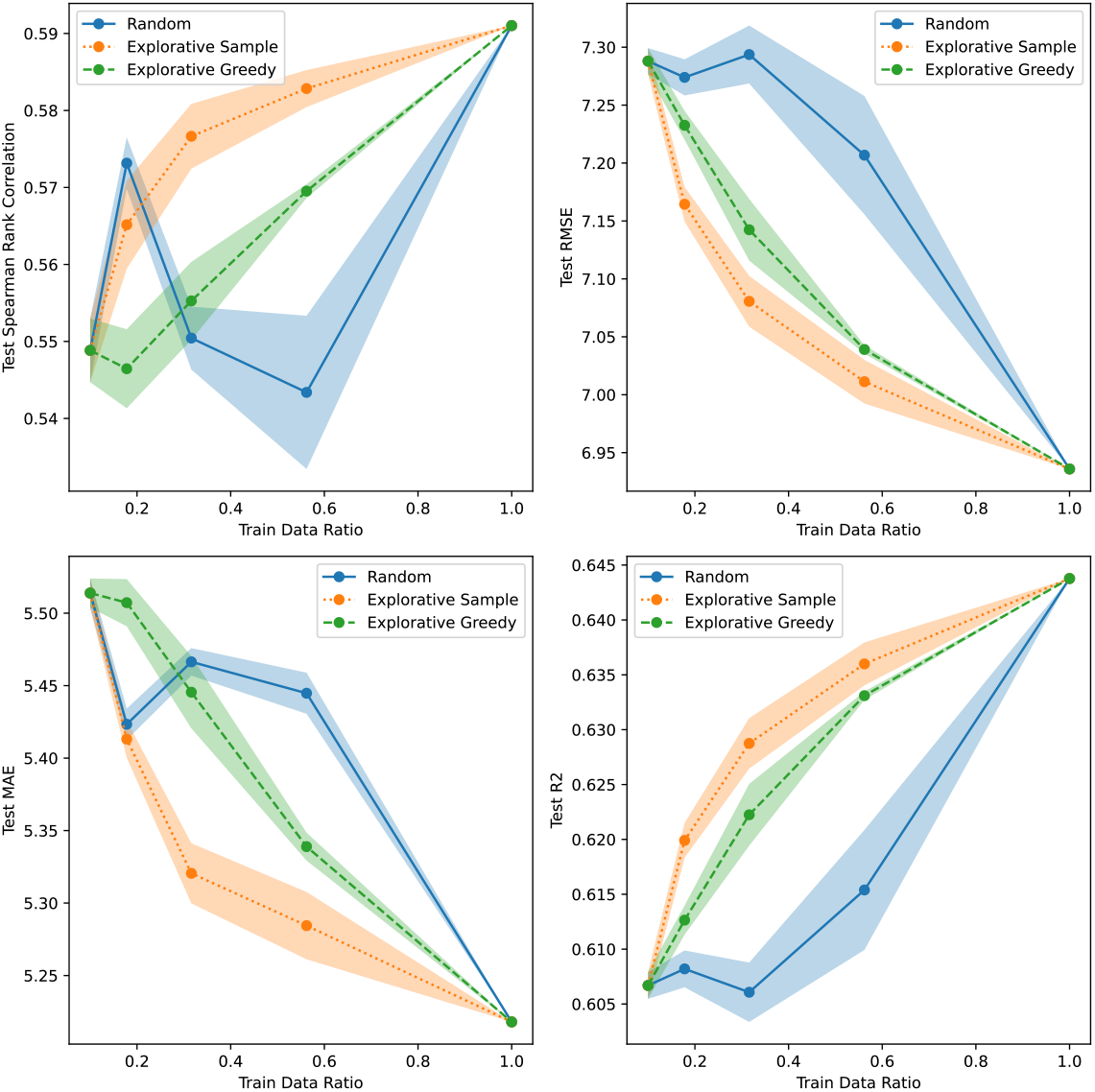
Active learning results for Meltome/Random using Linear Bayesian Ridge uncertainty.

